# To disperse or compete? Coevolution of traits leads to a limited number of reproductive strategies

**DOI:** 10.1101/2022.11.27.518112

**Authors:** Isaac Planas-Sitjà, Thibaud Monnin, Nicolas Loeuille, Adam L Cronin

## Abstract

Reproductive strategies are defined by a combination of behavioural, morphological, and life-history traits. Reproductive investment and offspring propagule size are two key traits defining reproductive strategies. While a substantial amount of work has been devoted to understanding the independent fitness effects of each of these traits, it remains unclear how coevolution between them ultimately affects the evolution of reproductive strategies, and how this might influence the relationship between dispersal and environmental factors. In this study we explore how the evolution of reproductive strategies defined by these two coevolving traits is influenced by resource availability and spatial structuring of the environment using a simulation model. We find three possible equilibrium strategies across all scenarios: a competitor strategy with high reproductive investment (producing large propagules which disperse short distances), and two coloniser strategies differing in reproductive investment (both producing small propagules which disperse long distances). The possible equilibrium strategies for each scenario depended on starting conditions, spatial structure and resource availability. Evolutionary transitions between these equilibrium strategies were more likely in heterogeneous than homogeneous landscapes and at higher resource levels. Transition from coloniser strategy to competitor strategy was usually a two-step process, with changes in propagule size following initial evolution in investment. This highlights how the interaction between the two trait axes affects the evolution of reproductive strategies, particularly where fitness valleys preclude the simultaneous evolution of traits. Our results highlight the need to incorporate trait coevolution into evolutionary models to help develop a more integrative understanding of the structure of natural populations and how the interaction between traits constrains or hinders evolutionary processes.

## Introduction

The central importance of reproduction in ecology and evolution is illustrated by the huge diversity of reproductive strategies exhibited both within and between species. This diversity likely reflects the result of an adaptive balancing of a complex suite of interacting trade-offs; between current and future reproduction, reproduction and somatic maintenance, size and number of offspring, and the relative importance of dispersal and competition (Geritz et al., 1999; Hamilton and May, 1977; Smith and Fretwell, 1974; Tilman, 1994; Weigang and Kisdi, 2015; Williams, 1966). Reproductive phenotype is thus the product of a complex mosaic of integrated traits, potentially including behavioural, physiological, morphological and life-history components (Bonte et al., 2012; Peiman and Robinson, 2017), and with some combinations of traits more beneficial in the eye of selection than others (Ronce and Clobert, 2012). However, the evolution of such complex reproductive strategies is typically modelled as a single parameter or gene locus (which may have multiple traits mapped to it) rather than a complex of interdependently evolving traits (though see Ronce, Perret and Olivieri, 2000). When multiple traits influence fitness interactively, selection can lead to genetic correlations through genetic linkage, pleiotropy and/or linkage disequilibrium (Endler, 1995; Roff and Fairbairn, 2012; Saltz et al., 2017; Sinervo and Svensson, 2002). Hence, while selection on reproductive strategies is imposed by environmental context, the adaptive potential of different strategies in different environments will depend on how the interaction between traits facilitates or constrains evolution (Collar et al., 2008; Dochtermann and Dingemanse, 2013; Lande, 1979; Lande and Arnold, 1983). Whether the coevolution of traits facilitates or constrains their evolution is an unresolved question (Kivelä, 2019). Given the potential importance of interactions between traits in regulating evolutionary responses, investigations of these patterns using models which incorporate multiple trade-offs and the coevolution of traits are needed (Weigang and Kisdi, 2015).

Life-history theory states that investing in reproduction comes at a cost to somatic maintenance, and thus organisms should balance investment in current reproduction against opportunities for future reproduction (Stearns, 1976; Williams, 1966). Reproductive investment can be expected to decrease with reduced availability of resources because priority is shifted to somatic maintenance (Fischer et al., 2009; McNamara et al., 2009), but may increase if these conditions become so extreme as to threaten future investment opportunities (Fischer et al., 2009; Williams, 1966). At the same time, this investment can be allocated to a single offspring or distributed among several offspring of smaller size. Thus, a trade-off between offspring size and offspring number exists for a given quantity of invested resources. In other words, increasing offspring size necessitates a reduction in the total number of offspring produced (Smith and Fretwell, 1974). Offspring size can thus be a strong determinant for survival, and while bigger propagules in general benefit from a higher competitive ability and establishment success (Allen et al., 2008; Coomes and Grubb, 2003; Tilman, 1994), it is typically negatively correlated with dispersal (Levin and Muller-Landau, 2000; see Bonte et al., 2012 for more details). Alternatively, producing a high number of offspring can increase colonisation ability. Therefore, for a given quantity of resources invested in offspring, this results in a trade-off between a focus on competitive ability (low number of larger offspring) or colonisation ability (more offspring with higher dispersal), commonly known as the competition-colonisation trade-off (Geritz et al., 1999; Tilman, 1994). In this context, competitor strategies can be favoured in stable, high resource environments which tend to be saturated and subject to high competition (Allen et al., 2008). Coloniser strategies, on the other hand, can be favoured under strong kin competition (Hamilton and May, 1977) and under conditions of high temporal heterogeneity or unpredictability (Friedenberg, 2003; Levin et al., 1984; Mathias et al., 2001), but selected against by increased spatial heterogeneity (Bonte et al., 2012; Cheptou et al., 2008; Hastings, 1983; Parvinen et al., 2020), though see Cronin et al (2016). This highlights the complexity of the interaction between the evolution of reproductive strategies and ecological context.

Various modelling studies have indicated that the trade-offs outlined above can combine to help define the structure of ecological communities, and can give rise to coexistence of different reproductive strategies in sympatry (Geritz et al., 1999; Mathias et al., 2001; Parvinen et al., 2020). For example, Tilman (1994) showed that any number of strategies could potentially coexist if traits are sufficiently dissimilar between competitors and colonisers (see also Calcagno et al., 2006). However, the relationship between competitive and colonisation ability can differ among species, and can determine their coexistence potential (Geritz et al., 1999). How the shape of the competition-colonisation trade-off affects the evolution of reproductive strategies in sympatry remains unknown.

In this study, we use a simulation modelling approach to elucidate the evolution of reproductive strategies in different spatially explicit environments, when reproductive phenotype is defined by multiple coevolving traits. To incorporate the complex nature of reproductive strategies into our model while maintaining tractability we distil reproductive strategies into two key evolving components: reproductive investment and size of offspring. We assume that these two traits have no genetic linking mechanism but jointly affect reproductive phenotype (Peiman and Robinson, 2017), and thus favourable combinations of traits can be maintained because of their adaptive advantage (Bell and Sih, 2007; Lande, 1979; Lande and Arnold, 1983; Sinervo and Svensson, 2002). Using a simulation modelling approach, we explore how spatial structuring of the environment, environmental quality and size-dispersal relationship, influence the evolution of reproductive strategies defined by coevolving traits.

## Methods

### Reproductive strategies

A reproductive unit produces propagules with a limited ability to disperse depending on physical constraints. Propagule in this study refers to an offspring unit, being either a single organism, or the funding unit in case of social insects (i.e., single queen in case of independent colony foundation, or a queen accompanied by workers in case of dependent colony foundation, budding or fission). We consider dispersal strategies as ranging along a continuum between two extreme strategies specialised on colonisation or competition. We define coloniser strategies as those producing small propagules with high dispersal distance and high number of reproduction attempts, but suffering from high mortality rates associated with dispersal. This strategy allows the exploitation of new habitats and avoids kin competition (Cronin et al., 2013). Alternatively, the competitor dispersal strategy produces big propagules that increase competitive and establishment ability, but reduces the dispersal distance and the number of reproductive attempts (each propagule requires high investment). These strategies also range in terms of parental investment, from a high investment strategy allocating most parental resources into offspring (e.g., terminal strategy) to a low investment strategy which invests minimally in offspring.

### Size-dispersal trade-off

The size of offspring strongly influences the competitive ability and dispersal distance of the reproductive strategy. While dispersal is typically negatively correlated with propagule size (Cronin et al., 2013), the shape of the relationship between dispersal distance and propagule size can vary depending on the dispersal organism, particularly when changes in investment lead to variation in dispersal mode (Calcagno et al., 2006; Eriksson, 2008; Peeters, 2012). For example, in many plants, small seeds can be dispersed by wind, while bigger gravity dispersed seeds grow close to the reproducing tree. In such case, the size-dispersal relationship can form a step-like function (i.e., organisms either disperse or not disperse), as larger seeds can have a dramatically reduced dispersal range after a certain size (Eriksson, 2008; Leslie et al., 2017). On the other hand, zoochory can modify this trade-off, allowing big propagules to disperse longer distances (Eriksson, 2008; Leslie et al., 2017), and thus this size-dispersal trade-off may take a more linear shape. Similarly, in ants, single queens can disperse on the wing, while swarms of queens and workers fission by dispersing dramatically shorter distances on foot; but flying social insects (e.g., Honey bee) or marine invertebrates experience a weaker trade-off between propagule size and dispersal distance (Cheptou et al., 2008; Cronin et al., 2013; Kisdi and Geritz, 2003; Massol and Cheptou, 2011; Tilman, 1994). Here, we define this size-dispersal relationship as strong (a step-like function) or weak (a declining function) trade-off depending on the shape of this relationship (see Fig. 1D), which can affect the evolution of dispersal strategies. We thus generated a set of simulations with a strong size-dispersal trade-off, and another one with a weak size-dispersal trade-off.

**Figure 1:**
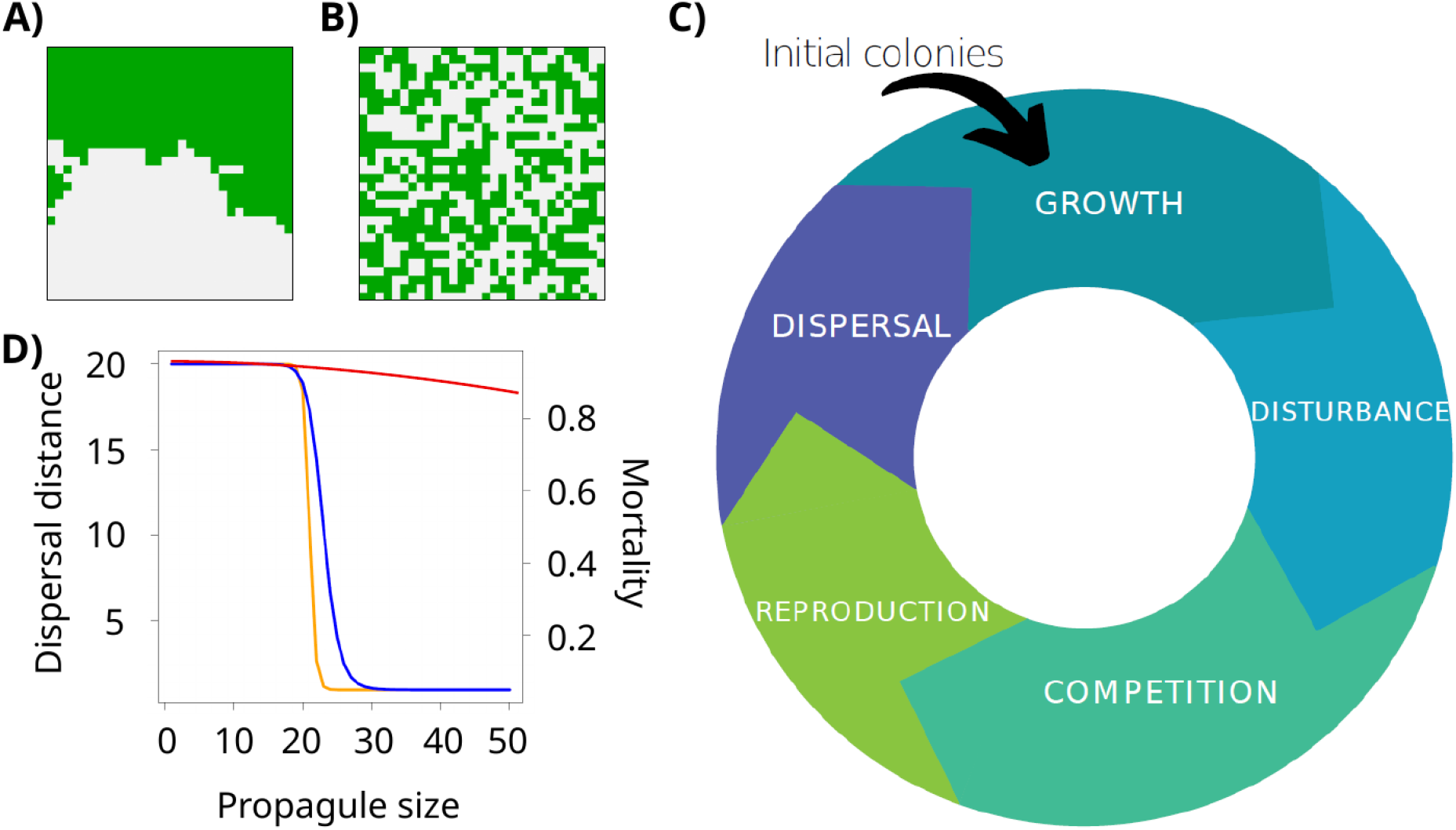
A) Example of Aggregated landscape, where green and gray patches represent high and low quality patches, respectively; B) example of Random landscape. C) flow diagram of the model. Colonies are introduced in the simulation; colonies grow; a stochastic extinction probability (ψ) is applied to each patch; in occupied patches, competition leads to extinction of all but one organism; if their size is higher than their investment threshold (E), they reproduce (if not, they remain in the patch waiting for the growth phase); when organisms reproduce, their offspring can be mutants regarding investment (E) and/or propagule size (S_o_); propagules disperse and land on occupied or empty patches. D) curves for the dispersal distance with strong (Eq. 2; yellow) and weak (Eq. 3; red) size-dispersal trade-off, and dispersal mortality (Eq. 4; blue) as a function of propagule size.

### Model outline

In this model we simulate organisms reproducing in landscapes with different degrees of heterogeneity and different levels of resources. The landscape consisted in a toroid lattice of 30 × 30 patches and we simulated three types of landscapes: Homogeneous, Random and Aggregated. All patches were defined by a quality *K* (available resources or carrying capacity). Homogeneous landscapes consisted entirely of ‘intermediate’ quality patches (*K* * 0.75), while Random and Aggregated landscapes consisted of even numbers of rich (*K* * 1) and poor (*K* * 0.5) patches, thus maintaining the same mean *K* values for comparable scenarios across different landscapes (i.e., same number of good and bad patches). All patches are therefore inhabitable and there is no fragmentation (although see below). High-quality (rich) and low-quality (poor) patches were distributed over the landscape using a fractional Brownian motion neutral landscape model (NLM) algorithm (see Sciaini et al., 2018) with either a high correlation index (1.2 for Aggregated landscapes, Fig. 1A) or low correlation index (0.001 for Random landscapes, Fig. 1B). For each landscape, we simulated four different levels of *K* (see Table 1).

**Table 1.** Parameters used in simulations and range of values tested for each parameter.

| *Parameter* | *Description* | *Value* |
| --- | --- | --- |
| $T$ | Simulation time (timesteps) | $10^6$ |
| $N$ | Initial number of colonies | 900 |
| $K$ | Maximum quality patch | 100, 500, 1000, 2000 |
| $P$ | Number of patch type | 2 |
| $\gamma$ | Spatial correlation index | Random=0.001; Aggregated=1.2 |
| $\psi$ | Stochastic extinction probability | 0.05 |
| $S_{init}$ | Initial organism size | 20 |
| $r$ | Growth rate (Fecundity - Mortality) | 1.2 |
| $\mu$ | Mutation probability | 0.001; 0.05 |
| $\varepsilon$ | Maximal mutation range [0-1] | 0.1 |
| $\beta_M; \delta_M; n_M$ | Minimum mortality probability;<br>maximum mortality probability;<br>steepness | 0.05 ; 0.95; 20 |
| $\beta_D; \delta_D; n_D$ | Minimum dispersal distance;<br>maximum dispersal distance;<br>steepness | Strong trade-off= [1; 20; 50] ;<br>Weak trade-off = [10; 20; 0.002] |
| $T_M; T_D$ | Threshold value (refers to $S_0$) for mortality (M) and dispersal (D) | 23 ; 21 |

Organisms were modelled as agents with reproductive strategies defined by two evolving traits: reproductive investment (or energy, *E*), and propagule size (or size of offspring, *S_o_*). Simulations followed the cycle of events shown in Fig. 1C, which can be summarised as follows. At *t* = 0, the landscape is populated with agents of an initial size *S_init_*, each with the default values of reproductive investment (*E)* and propagule size (*S_o_*) (see Table 1). At each timestep *t*, representing one reproductive season, organisms grow, compete for resources with others within the same patch, and produce new propagules that will disperse over the landscape. During reproduction, these propagules have a probability of mutation in *E* and/or *S_o_* that may affect the dispersal strategy of the next generation.

During the growth phase, organisms grow following the Ricker logistic equation (Ricker, 1954), which has been broadly used for competition models in discrete time:

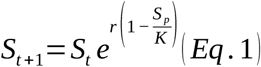

Where *S_t_* is the size of the organism at time *t*, *r* is the growth factor, *K* is the carrying capacity of the patch, and *S_p_* is the cumulative size of all organisms present in the patch. Thus, the growth of an organism depends on the number of agents present in the same patch, as well as their size (larger organisms gather more resources).

Following growth, each patch is subject to a stochastic extinction probability (*ψ*) to model random environmental disturbance. This represents a probability that all organisms in a given patch are destroyed, thus producing empty patches at every timestep and favouring dispersive strategies through increased selection for bet-hedging (Cronin et al., 2016; Kivelä, 2019; Levins, 1969).

At the end of year *t*, we allow each patch to be occupied by only one organism, and thus if more than one organism is present, a ‘winner’ is selected via a multinomial probability competition, where the probability of winning is proportional to colony size. This reflects competition for resources and aggression which usually results in larger organisms outcompeting smaller ones.

During the reproduction phase, organisms large enough to reproduce *(Size* > *E*),produce *N_o_* propagules, with *N_o_* =*E / S_o_* (fraction being rounded to the nearest lower integer). These propagules inherit the reproductive traits (*E; S_o_*) of the reproducing organism, unless a mutation occurs with a probability *μ*, calculated independently for each trait and each new propagule. When a mutation occurs, the value of the new trait (*E*’ and/or *S_o_*’) is drawn from a uniform distribution of *E* ± *E · ε* and/or *S_o_* ± *S_o_* · *ε*, with *ε* being the maximum amplitude of a mutation (range [0 - 1]). Note that the maximum mutation amplitude is therefore proportional to the trait value (as both traits are quantitative). As these mutations are restricted to the new propagule, they do not affect the reproductive agent or the dispersal of the propagule itself, but will define the offspring phenotype of the new propagule. If propagule size mutates to exceed investment (*S_o_* > *E* hence *N_o_* < 1), or investment exceeds the *K* of the patch, the new colony will not be able to reproduce. Such strategies are thus never selected for in the long term, as they are eliminated by stochastic extinction *ψ* or competition.

Propagules disperse in a random direction and at a distance from the reproductive agent obtained from a Poisson distribution centred on the corresponding value of equation 2 & 3. Those which survive become established in a patch. The size of new propagules is defined by the *S_o_* parameter of the reproductive agent. After reproduction, the energy used for reproduction (*N_o_* * *S_o_*) is reduced from the total size of the organism, representing the cost of the reproductive event to the parent organisms.

### Size-dispersal trade-off

To model the relationship between *S_o_* and dispersal, we considered either a strong or a weak size-dispersal trade-off, meaning that competition ability comes at a high or low cost of dispersal, respectively. In both cases, the dispersal distance (*D*; Eq. 2 & 3) is defined as a function of *S_o_* (see also Fig. 1D):

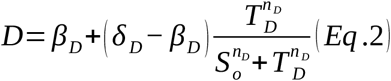

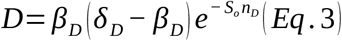

We used equation 2 for the strong dispersal trade-off. This equation generates a step function at a certain threshold value *(T_D_)* with a given steepness *(n_D_)* considering maximum *(δ_D_)* and minimum *(β_D_)* value of dispersal distance (Fig. 1D). This strong tradeoff between size and dispersal means that, at a certain threshold propagule size *T_M_*, propagules switch from long-range dispersal (*δ_D_*) to short-range dispersal *(β_D_*).

For the weak dispersal trade-off, we used equation 3 to model dispersal distance in function of *S_o_*. This equation uses the same parameters described above but with different values (Table 1) to generate a smooth decreasing function representing species without a switch of dispersal mode (e.g., bees, or seeds dispersed by animals).

### Size-mortality trade-off

Mortality probability due to dispersal was modelled in function of *S_o_* too. We used equation 4:

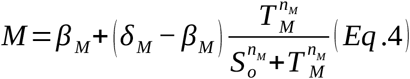

which, similar to equation 2, generates a step function at a certain threshold value (*T_M_*) with a given steepness (*n_M_*) considering maximum (*δ_M_*) and minimum *(β_M_* value of dispersal mortality (Fig. 1D). This step function generates a strong trade-off between propagule size and mortality, so that small propagules (coloniser strategies) have a high mortality risk while big propagules (competitor strategies) have a low mortality risk. Thus, *β_M_* largely determines the mortality probability for large propagules, while *δ_M_* determines the mortality probability of small propagules. In lack of empirical evidence for a general dispersal mortality cost curve (although see Bonte et al., 2012b), we kept the same dispersal mortality function for all simulations. This makes sense as organisms with either strong or weak trade-off may have a drastic decrease of dispersal mortality after a certain size threshold (Beckman et al., 2018; Cronin et al., 2013), and organisms dispersing farther distance have higher probabilities to land in different and unknown environment, which incurs higher mortality. Thus, in the case of a strong size-dispersal trade-off, the combination of equations 2 & 4 generates a high non-linear relationship between dispersal distance and dispersal mortality, as long-range propagules suffer from high mortality, while short-range propagules have low mortality. For a weak size-dispersal trade-off, the relationship between distance and mortality is less extreme, as big propagules with reduced mortality come at a minor cost in dispersal distance.

### Mutation rate

We assume that evolution happens due to random mutations, although we have little empirical information about mutation rates in nature. Therefore, to assess the effect of different mutation rates, we used two different mutation speeds for the co-evolving traits, and we refer to them as a high (0.05) and low (0.001) mutation rates. As both sets of simulations led to comparable outcomes, and simulations with low mutation rate did not reach a stable state in some cases at time *T*, we present the results with a high mutation rate and discuss the little differences observed in the discussion.

### Simulations

To investigate whether different starting conditions lead to divergent equilibrium states or converge to the same final strategy in a particular scenario, we tested four different starting conditions of *E* and *S_o_*. These starting conditions corresponded to three extreme strategies and one intermediate strategy in terms of investment and propagule size, broadly representing two competitor-like and two coloniser-like starting conditions (Fig. 2). Specifically, the starting conditions are: high-investment competitor (high *E* and high *S_o_*), medium investment competitor (intermediate *E* and *S_o_*), high-investment coloniser (high *E* and low *S_o_*), and low-investment coloniser (low *E* and low *S_o_*). In general terms, high and medium investment competitors could be understood as strategies producing one or two relatively large propagules with lower dispersal and low dispersal mortality. On the other hand, high- and low-investment colonisers are strategies producing small propagules with long-range dispersal but high dispersal mortality. To ensure that each starting strategy was viable in the different environmental scenarios tested, the starting values of investment and propagule size in each scenario were based on maximal resources *K_max_* for that scenario, where *K_max_* = *K* for Heterogeneous landscapes (Aggregated and Random) and *K_max_* = *K* * 0.75 for Homogeneous landscapes. Investment *E* for high-investment strategies was set at *K_max_* - 5, while *E* for the low-investment strategy was set at 50. For the medium investment strategy, *E* was set at *K_max_* /2. In all starting conditions, *S_o_* was set at *E* – 45, except for the high-investment coloniser, for which it was set at 5 (see Fig. 2). As a result, starting conditions differed slightly between Homogeneous and Heterogeneous landscapes, especially for low resource environments (see Fig. 3 – 5).

**Figure 2:**
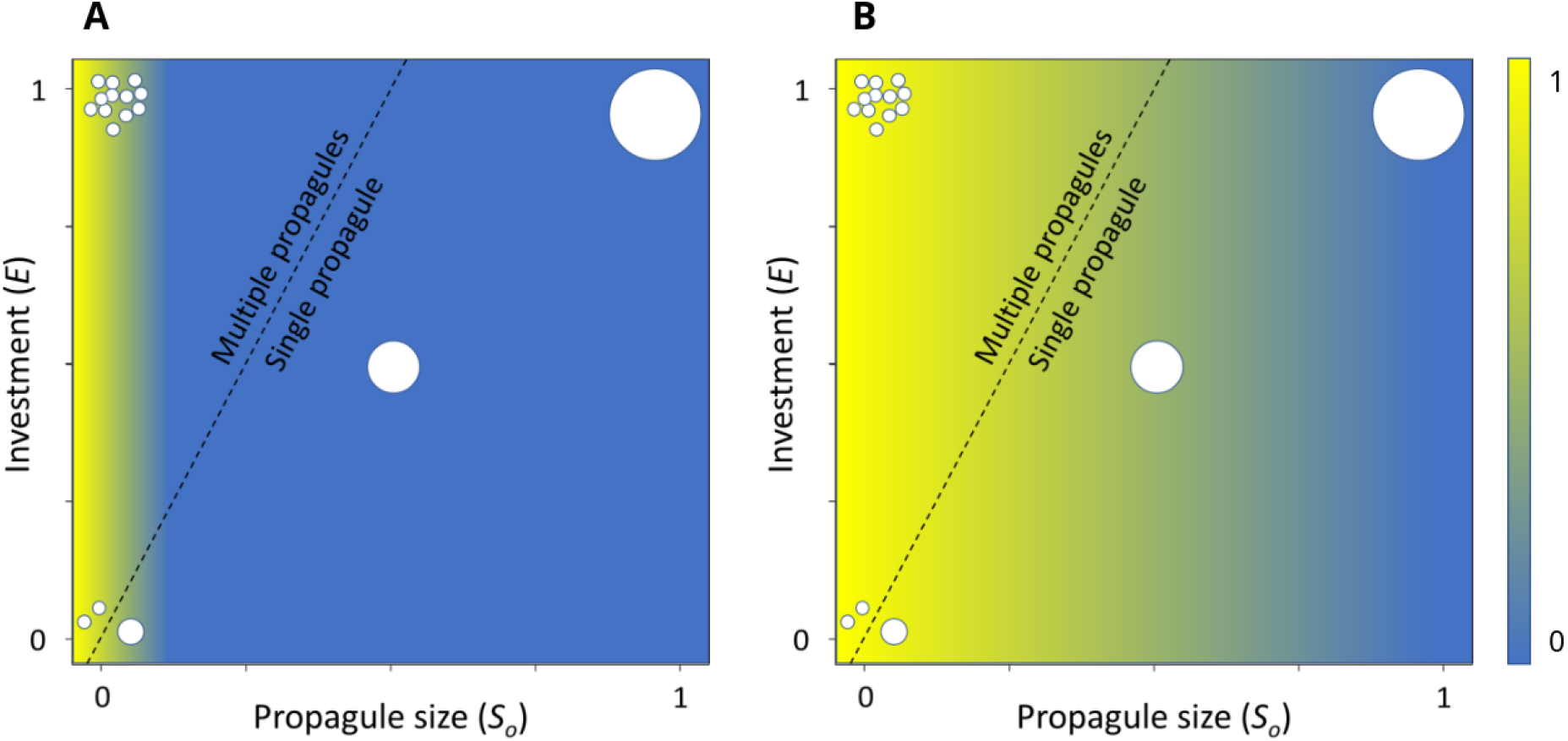
Example of starting conditions used in our simulations. Axes (E and S_o_) are written as proportions of the total resource K available in a given landscape. Dashed line indicates parameter space producing single or multiple propagules, blob size represents the size of the propagules, and colour indicates dispersal distance, for (A) with a strong sizedispersal trade-off or (B) a weak trade-off.

**Figure 3.**
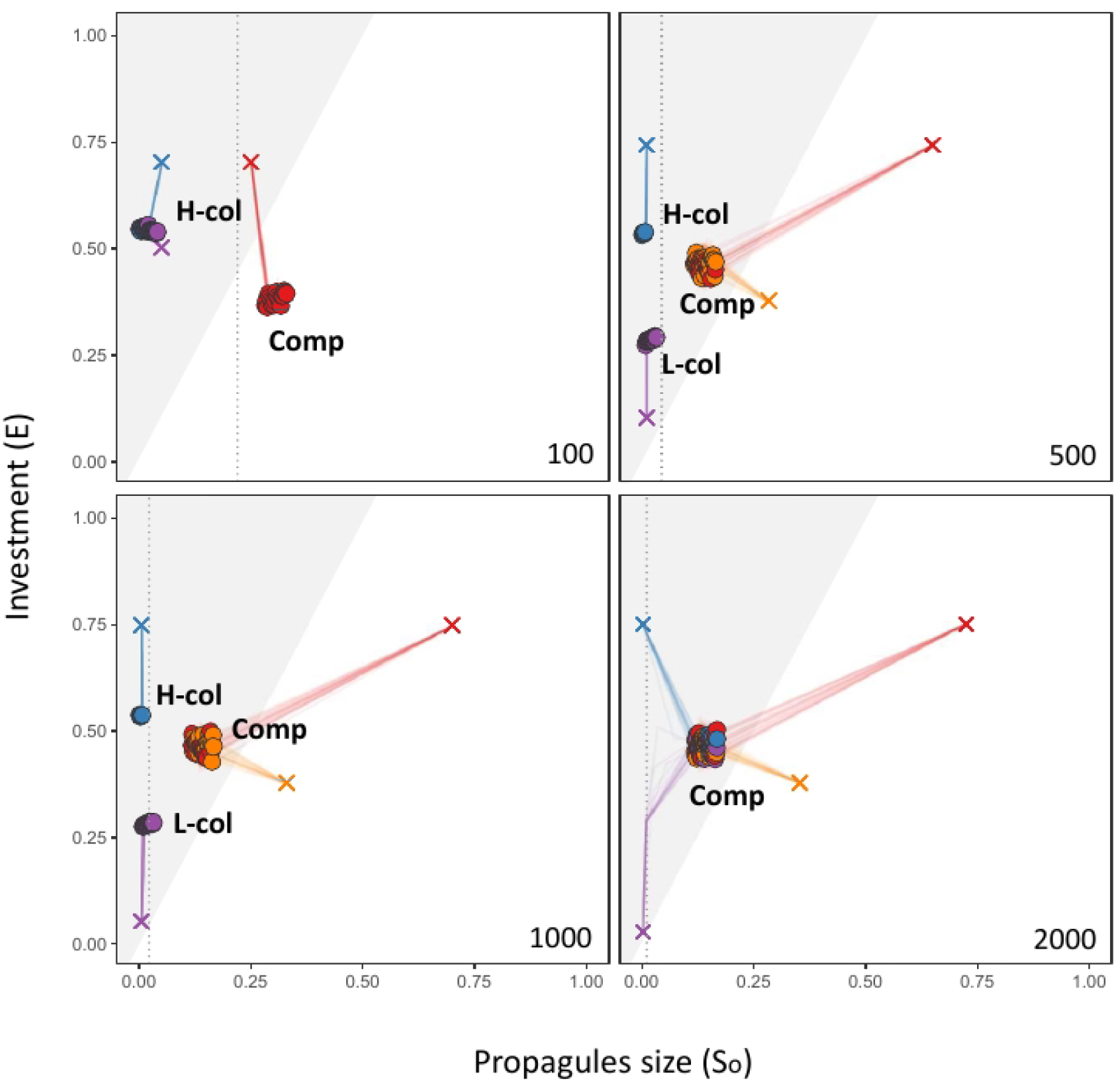
Evolution of strategies in Homogeneous landscapes. Filled circles indicate the final (= mean of all colonies in a simulation) investment (E) and propagule size (S_o_) for different starting conditions for the Homogeneous landscape at different resource levels (K = 100, 500, 1000, 2000). H-col = High-investment coloniser; L-col = Low-investment coloniser; Comp = competitor. Note that axes (E and So) are written as proportions of the resource level K for that landscape. Starting conditions are indicated with a cross and lines are the temporal dynamics of E and S_o_ during simulations (values captured every 10^4^ time-steps). Each coloured straight line corresponds to the mean value of E/S_o_ of all colonies in a given simulation. The shaded area indicates trait combinations producing more than one propagule, while the unshaded region indicates a single propagule. Bold letters designate clusters of final equilibrium strategies. The dotted line indicates the limit between propagules with high dispersal (left of line) and propagules with low dispersal. The position of points has been randomly shifted slightly to aid visualisation (exact values can be extracted from Fig. S2).

**Figure 4.**
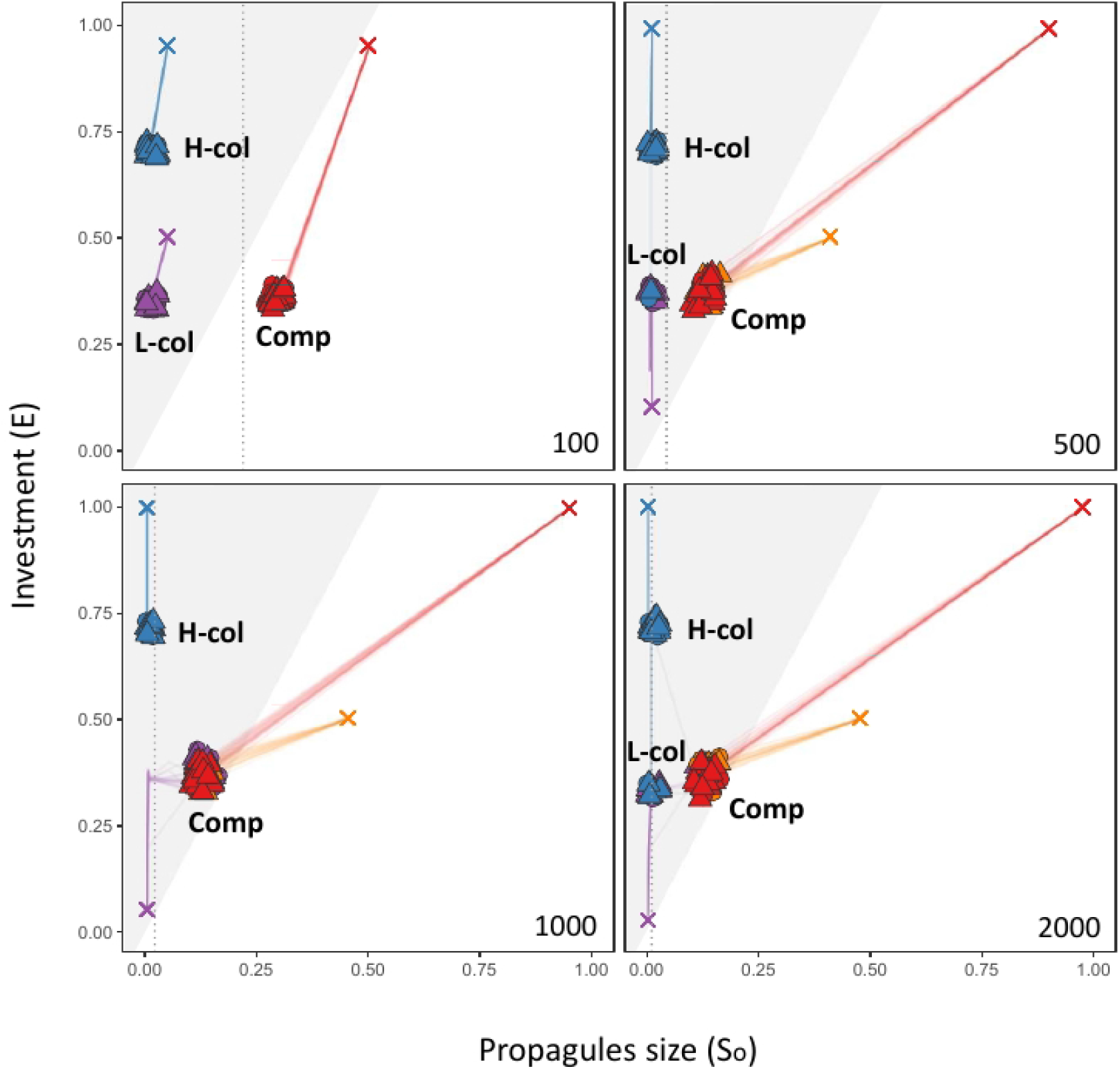
Evolution of strategies in Random landscapes. Details as for Figure 3, except that circles indicate colonies in rich patches and triangles indicate colonies in poor patches.

**Figure 5.**
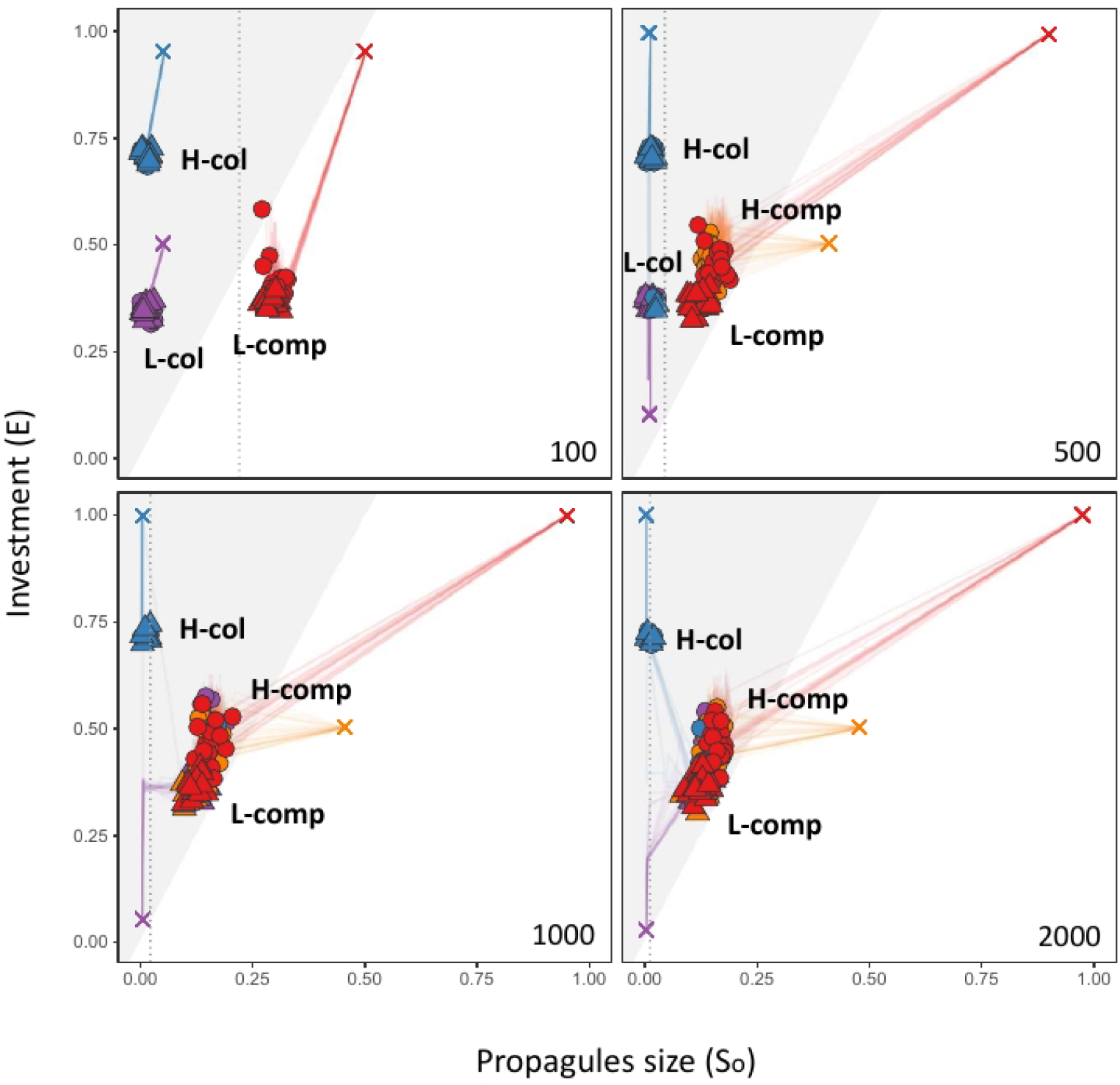
Evolution of strategies in Aggregated landscapes with strong size-dispersal trade-off. Details as for Figure 4. H-comp = High-investment competitor; L-comp = Low-investment competitor.

We performed 20 replicate simulations for each unique set of parameters. We used the same set of 20 different randomly generated maps (available as File S1) for each of the described scenarios, and all simulations had a time limit of *T* = 10^6^. We confirmed that this time limit was enough for simulations to reach equilibrium values of *E* and *So* by visual checks of temporal dynamics (File S2). All parameter values in our simulations (Table 1) were chosen arbitrarily and based on pilot runs as points from which effects of variation could be explored (Bonte et al., 2012; Cronin et al., 2016).

We extracted the following results for each organism at the end of each simulation: position (x,y), patch quality, size, age, lineage (i.e., track descendants from original agents), *E* and *S_o_*. We also captured the number of agents, distribution of *E* and *S_o_*, and mean (± SD) of *E and S_o_* in rich/poor patches every 10^4^ time-steps.

### Invasion analysis

Our evolutionary simulations resulted in several possible equilibrium strategies (see Results). We therefore assessed whether the trait combinations present in these equilibrium conditions represented global evolutionary stable strategies (ESS) using an invasion analysis. Equilibrium strategies were defined as distinct clusters with trait combinations of *E* and *So* remaining at the end of simulations (see Figs. 3–5), the number of which varied among scenarios. For each scenario (landscape type and resource availability), we selected one of the possible equilibrium strategies as a ‘resident’ strategy, and populated the landscape with a randomly selected agent from the focal strategy. At timestep 10, allowing enough time to grow and reproduce, we introduced a single ‘invader’ agent with parameters values taken from one of the other equilibrium strategies for that scenario in the same manner. If one of the equilibrium strategy clusters was absent from that scenario, parameters were taken from an alternative scenario for the same landscape containing such a cluster. Simulations were carried out as outlined above except without mutation. We did 10^3^ invasion simulations for each possible combination of resident and invader equilibrium strategies in each scenario. Simulations were halted when only one strategy remained or after 2·10^4^ timesteps (preliminary simulations indicated that this time was enough for a single invader colony to populate the whole landscape or to reach a stable equilibrium of coexistence). This invasion analysis was done only for scenarios with mutation rate = 0.05, and was done for strong and weak size-dispersal trade-off scenarios separately. Equilibrium strategies were classified as ESSs if they were never successfully invaded by any of the other equilibrium strategies. Simulations in which both invader and resident strategy were present at the end of the simulation were classified as conditions supporting coexistence.

## Results

We first consider the results of simulations using the high mutation rate (0.05) and strong size-dispersal trade-off, then explore the effects of varying these parameters.

### Evolution of reproductive strategies in homogeneous environments

We found three possible equilibrium strategies in Homogeneous landscapes, though outcomes depended on both starting conditions and resource availability. In all cases, only one equilibrium strategy was observed in a single simulation (see Fig. S1 for example of single simulation outcomes). Thus, scenarios with more than one equilibrium strategy indicate conditions in which different replicates using the same parameters produced different equilibrium strategies. Traits evolved away from starting conditions in all cases, though evolution in investment *E* was not always accompanied by evolution in propagule size *S_o_*.

Competitor start-conditions converged on an intermediate competitor strategy (~50%*E*) in all scenarios, while coloniser start-conditions remained either high- or low-investment coloniser strategies in scenarios with intermediate level of resources (Fig. 3). The convergence of coloniser start-conditions on a single high-investment coloniser strategy with extremely low resources (*K* = 100), was likely facilitated by the fact that traits of starting conditions were very similar (Fig. 3). In scenarios with high resources (*K* = 2000), coloniser start-conditions converged on the aforementioned final competitor strategy (Fig. 3). Thus, the evolution from coloniser to competitor strategies was facilitated by the availability of resources. Interestingly, in this case evolution in *E* was necessary before any evolution could occur in *S_o_* for the low-investment coloniser start-condition to evolve to a competitor strategy (Fig. 3, K = 2000). This likely reflects the fact that mortality was a non-linear function of propagule size but was a linear function of investment (through competition effects), and illustrates how achieving the final competitor reproductive strategy is dependent on allowing two-dimensional evolutionary dynamics.

### Evolutionary outcomes in heterogeneous environments

In Random and Aggregated landscapes, we once again observed three equilibrium strategies, with only one equilibrium outcome per replicated simulation. The two competitor start-conditions again converged on a final competitor strategy in all scenarios, whereas outcomes for the coloniser starting conditions varied depending on resource availability (Figs. 4 and 5). Evolution in *S_o_* for both coloniser start-conditions was preceded by evolution in *E* in all cases. Convergence of coloniser start-conditions on the competitor strategy was less common in Random landscapes (27 simulations) than in Homogeneous landscapes (40), but more common in Aggregated landscapes (51) than for either of the other landscapes. Propagule size for all clusters in Random landscapes was comparable to that observed in Homogeneous landscapes, although investment for the competitor strategy was lower in Random landscapes, and higher for coloniser strategies. In Aggregated landscapes (Fig. 5A), the equilibrium competitor cluster spanned a notably broader range of trait values than in Homogeneous or Random landscapes, with poor-patch colonies having lower *E* and, to a lesser extent, *S_o_*.

To summarize, high resource quality increased the rate of evolution from coloniser to competitive strategies, while resource distribution affected the stabilizing selection of dispersal strategies. Competitive strategies were favoured in less fragmented landscapes (Homogeneous and Aggregated landscapes) with high level of resources, while colonisers were favoured in landscapes with low-intermediate levels of resources and heterogeneous environments.

### Effect of mutation rate

Transitions among final strategies were facilitated by the high mutation rate (0.05), which increased the probability of convergence, and reduced the number of final strategies in rich scenarios (high *K*). Results with the low mutation rate also highlight the two-step process required to evolve from coloniser start-conditions to competitor final condition, where evolution in *S_o_* is preceded by evolution in *E* (see File S3). For competitor strategies, low mutation rate (0.001) constrained the evolution towards the production of two propagules, instead of one, produced a less directional selection and had low stabilising selection (File S3).

### Effect of size-dispersal trade-off

The strength of the dispersal trade-off had a quantitative, but not qualitative, effect. The strong trade-off resulted in a slight reduction of investment in most scenarios, except for the poorest landscape (*K* = 100) where colonies had higher investment compared to simulations with a weak dispersal trade-off (see also File S4).

The weak dispersal trade-off produced more stabilising selection (i.e., no differences between good and bad patches) in Aggregated landscapes (Fig. 5B), and there were not differences between Aggregated or Random landscapes (Fig. 5 and S4).

### ESS and coexistence conditions

As we never observed the emergence of multiple equilibrium strategies within a given simulation, it remains unclear which outcomes, if any, represent evolutionary stable strategies. We therefore tested the potential for each equilibrium strategy to resist invasion by other equilibrium strategies in all scenarios. For Homogeneous and Random landscapes, we used the three final equilibrium clusters (see Fig. 3–4), while for Aggregated landscapes, the competitor strategy was further divided into low-investment competitor and high-investment competitor, with these defined as competitor colonies inhabiting poor and rich patches respectively (Fig. 5).

ESSs were found in all landscapes, though none of the equilibrium strategies was an ESS across all scenarios (Fig. 6). Outcomes were resource dependent in Homogeneous and Aggregated landscapes, but consistent across resource levels in Random landscapes. In Random and Homogeneous landscapes the competitor strategy was always an ESS, and coexistence of two strategies was possible through invasion in Aggregated landscapes. The potential for coexistence depended on resource level and strength of size-dispersal trade-off. In general, a weak size-dispersal trade-off decreased the chances of coexistence between final strategies (File S4). Coloniser (low-investment) and competitor strategies coexisted across resource levels only with a strong trade-off (Fig. 6), while both coloniser (high-/low-investment) strategies coexisted in intermediate resources (*K* = 1000), no matter the strength of the trade-off. Finally, a high-investment coloniser strategy resisted the invasion of a low-investment coloniser only in poor (*K* = 100) Homogeneous landscape, otherwise the low-investment strategy was selected.

**Figure 6.**
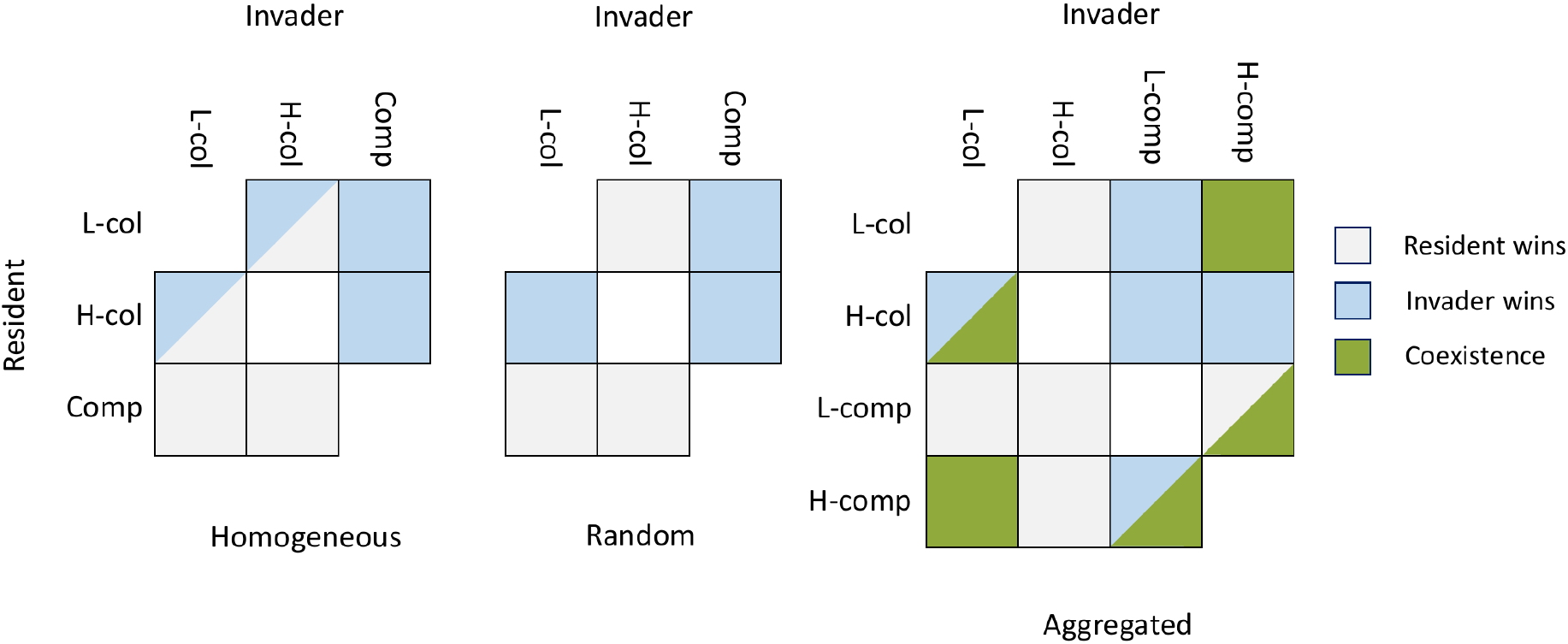
Invasion analysis for final strategies with strong trade-off. Green squares indicate scenarios in which coexistence of strategies was found in at least one simulation; grey indicates that no coexistence was observed and the resident strategy persisted, while blue indicates that the invading strategy replaced the resident. Split cells indicate scenarios where the outcome varied depending on resource level (see File S5 for details). L-col = Low-investment coloniser; H-col = High-investment coloniser; Comp = Competitor; L-comp = Low-investment competitor; H-comp = High-investment competitor.

## Discussion

In this study, we investigated the coevolution of traits defining reproductive strategies under different resource levels, patterns of spatial structuring and for differing strengths of the dispersal trade-off. In our simulations we observe three possible equilibrium strategies: a high-investment coloniser, low-investment coloniser, and competitor. While no equilibrium strategy in our model was an ESS in all conditions, the competitor strategy (with ~50% investment) was never excluded in invasion analyses, and coexisted with other strategies when not an ESS. The high- and low-investment coloniser strategies produced numerous small propagules but differed in levels of reproductive investment (~70% and ~25-40% of resources respectively). Energy invested in adult survival is implicit in our model, as energy not invested in reproduction defined the size of the parent and therefore the probability of surviving competitive challenges from new propagules. The two equilibrium coloniser strategies may therefore represent alternative investment optima along the trade-off between adult survival and fecundity (Endler, 1995; Sinervo, 2000; Williams, 1966; Winkler & Wallin, 1987): a fast-growing, short life span (‘semelparous-like’) strategy focussing on immediate reproduction or ‘terminal investment’ (high-investment coloniser), and slow-growing, long life-span (‘iteroparous-like’) strategy emphasising adult longevity (low-investment coloniser) (Salguero-gómez et al., 2016). [Nico mentioned “Stable environment may give a head-start to semelparity”]--> this idea could be introduced here too

Start conditions strongly influenced the possible final equilibrium strategies. Competitor start-conditions always evolved to the competitor strategy, and never to a coloniser strategy, whereas coloniser start-conditions could evolve to coloniser or competitor strategies. Evolution between the possible equilibrium strategies was unidirectional, either from high-investment coloniser to low-investment coloniser, or from high or low-investment coloniser to competitor. These patterns match the outcomes of invasion analysis in that winners of contests were the end-points of these transitions. One possible explanation for the overall success of the competitor strategy is that the low number of empty patches (stochastic extinction probability = 0.05) generated only weak selection for dispersing morphs (Comins et al., 1980; Duputié and Massol, 2013). The unidirectional evolution from high-to low-investment coloniser equilibrium strategies may be explained by high mortality of dispersing propagules favouring reduced investment in reproduction (Law, 1979; Reznick et al., 1990; Williams, 1966), which can also allow more repeat breeding attempts (Martin, 2014). This directional selection agrees with evolution of dispersal in social insects, in which a coloniser strategy is thought to be ancestral but has repeatedly given rise to competitor strategies (Cronin et al., 2013; Eriksson, 2008; Peeters, 2012), and in angiosperms, in which small seeds gave rise to larger seeds and fruits during the Tertiary period (Eriksson, 2008). Evolutionary transitions from competitor to coloniser strategy have not been documented to our knowledge in social insects, though we might expect long-term costs of such low-dispersal strategies (Hamilton and May, 1977) to favour this transition or simply lead to extinction, and this awaits further study.

Evolution for each start-condition was limited to local fitness optima in low resource environments, although transitions to other fitness optima became more likely in higher resource environments, and in heterogeneous landscapes. Transitions from coloniser starting conditions to the competitor strategy occurred only at intermediate-high resource levels (1000+). Indeed, low-investment strategies, for colonisers or competitors, could be more adaptive in rich environments because increased resource availability can annul selection for terminal investment strategies, favouring more balanced investment (Fischer et al., 2009). Additionally, this increased likelihood of transition is likely linked to higher mutation amplitudes, as mutation amplitude was proportional to trait values, which were themselves resource dependent. Higher mutation amplitude in higher resource environments may thus have facilitated the crossing of fitness valleys between local fitness optima which were unsurpassable in other landscapes. Accordingly, transitions were also constrained by lower mutation rates, which potentially obstructed the crossing of these fitness valleys. Finally, transitions from the high-investment coloniser to the low-investment coloniser strategy occurred only in heterogeneous landscapes and at intermediate-high resource levels (500+), suggesting resource-independent spatial effects.

Trait values of equilibrium strategies were largely consistent across scenarios, with two exceptions. Firstly, propagule size of the competitor strategy was largest in the lowest resource environment, matching predictions of increased investment in offspring to improve offspring survival in poor environments (Armbruster et al., 2001; Fox and Czesak, 2000). This also agrees with the outcome of the invasion analysis, in which a coloniser strategy with high-investment, instead of low-investment, was selected in poor Homogeneous environments. Secondly, while spatial distribution had no effect on final strategies under a weak size-dispersal trade-off, there was a broadening of equilibrium trait values in Aggregated landscapes with a strong size-dispersal trade-off, particularly for investment *E*, suggesting weaker stabilising selection here. This broader range of investment values was linked to patch occupation, with lower investment associated with occupation of low-quality patches, and that was true for high/low mutation rates. Interestingly, trait values in rich patches of the Aggregated landscapes were comparable to those observed for the competitor strategy in Homogeneous landscapes, while trait values in poor patches were comparable to those observed in Random landscapes (Figs. 3–5). This may suggest that trait evolution in Random landscapes is constrained by the quality of poor patches, whereas this limitation is locally relaxed in clusters of good patches in Aggregated landscapes when a strong trade-off is at work. As reproduction is only possible when investment *E* is lower than the *K* of the habited patch, the maintenance of low investment for coloniser strategies in random landscape allow propagules to colonise empty patches, whether rich or poor (Geritz et al., 1999; Weigang and Kisdi, 2015).

Ecological models have shown that competitor strategies can be favoured in stable, high resource environments while coloniser strategies can be favoured under strong kin-competition and high temporal heterogeneity, but selected against by increased spatial heterogeneity (Bonte et al., 2012; Cheptou et al., 2008; Hamilton and May, 1977; Hastings, 1983; Mathias et al., 2001; Parvinen et al., 2020). However, the evolutionary consequences of environmental heterogeneity in the context of coevolving traits remain unclear (Duputié and Massol, 2013; Kivelä, 2019; Massol et al., 2010). While our results support these predictions in indicating that competitors were favoured in uniform and higher resource environments, we did not observe selection against colonisers under conditions of high spatial heterogeneity (Random landscapes). One possible explanation for these patterns is that while the competitor strategy represents a broad fitness optimum, evolution to this strategy was precluded in some scenarios (see above). The competitor strategy may also suffer from high kin-competition within the rich patches of Random landscapes, as rich patches are surrounded by poor patches in which competitor colonies cannot initially reproduce (i.e., their investment *E* is higher than resources in the patch). Thus, the combination of the competition-colonisation trade-off imposed in our simulations and the extreme spatial heterogeneity of this landscape could favour the selection of dispersal phenotypes (Cronin et al., 2016; Gross, 2008). Alternatively, the persistence of colonisers under high spatial heterogeneity could reflect differences between the coevolutionary approach we implemented and single trait evolution models. If spatial heterogeneity modifies the energy invested in reproduction *E*, which in turn affects the propagule size *So* (and thus dispersal strategy), this could have inhibitory or synergistic effects on evolution.

In our simulations, evolution from coloniser starting conditions to the competitor strategy was usually a two-step process, with change in propagule size following initial evolution in investment. This clearly highlights the interactive process existing between the two trait axes. The different evolutionary dynamics this introduces may help explain the coexistence of several dispersal strategies in heterogeneous landscapes (Cronin et al., 2016; Massol et al., 2010; Weigang and Kisdi, 2015), and may explain why evolution from coloniser start-conditions to the competitor equilibrium strategy (requiring sequential evolution of traits) were common, but evolution from the competitor start-conditions to either coloniser equilibrium condition (requiring simultaneous evolution of traits) did not occur. Coevolution of traits is thought to have the potential to both facilitate and constrain evolution (Collar et al., 2008; Dochtermann and Dingemanse, 2013; Lande, 1979; Lande and Arnold, 1983). Evolution of multiple traits can facilitate adaptation to new niches which cannot be ‘reached’ through evolution in a single trait alone (e.g., Collar, Wainwright and Alfaro, 2008). At the same time, the viable trait-space may be restricted by coevolution because conditions supporting evolution of a single trait are rare (e.g., Díaz et al., 2016). That we found only three possible equilibrium strategies in our broad parameter space suggest strong stabilising selection acts on combinations of the two traits defining reproductive strategy in our model.

We only observed small differences between outcomes of scenarios considering a strong or weak size-dispersal trade-off. This suggests that the obtained results may be applicable to a broad range of species using diverse dispersal strategies. Additionally, it could also indicate that the size-mortality trade-off has a higher impact on the evolutionary outcome of dispersal strategies than the size-dispersal trade-off. Further studies could investigate the relationship between both trade-offs in order to assess the relative importance of each one. Finally, our approach considers that all patches are inhabitable, although in real world habitats may be fragmented, with inhabitable patches separated by uninhabitable patches. Further studies could investigate whether fragmentation accentuates or decreases differences between different dispersal trade-offs.

In this study, coexistence between different reproductive strategies only arose through subsequent invasions, highlighting the potential importance of immigration from other populations in maintaining coexistence of strategies in a population (see Hanski, 1985). These results suggest that either natural environments include contexts more conducive to the evolution of such polymorphisms (eg. temporal fluctuation of resources, higher differences in resource quality) than in our model, or the intriguing possibility that such intraspecific polymorphisms arise via subsequent invasion. Our study shows that coevolution of traits may limit the number of possible complex phenotypes, although further analyses of coevolutionary dynamics in organisms with different life-histories will enable us to assess the veracity of this finding. Finally, we show that the consideration of multiple trait axes and coevolutionary interactions introduces different evolutionary dynamics in reproductive strategies, which may help develop a more integrative understanding of the structure of associated populations.

## Supporting information

Example of simulations

Homogeneous landscape supplementary figure

Rasters used

Dynamics for Homogeneous, Random and Aggregated landscapes with strong trade-off and high mutation rate

Dynamics for Homogeneous, Random and Aggregated landscapes with strong trade-off and low mutation rate

Dynamics for Homogeneous, Random and Aggregated landscapes with weak trade-off

Invasion analysis

## Notes

### Competing Interest Statement

The authors have declared no competing interest.

