## Supplementary material for "To disperse or compete? Coevolution of traits leads to a limited number of reproductive strategies": Example of simulations

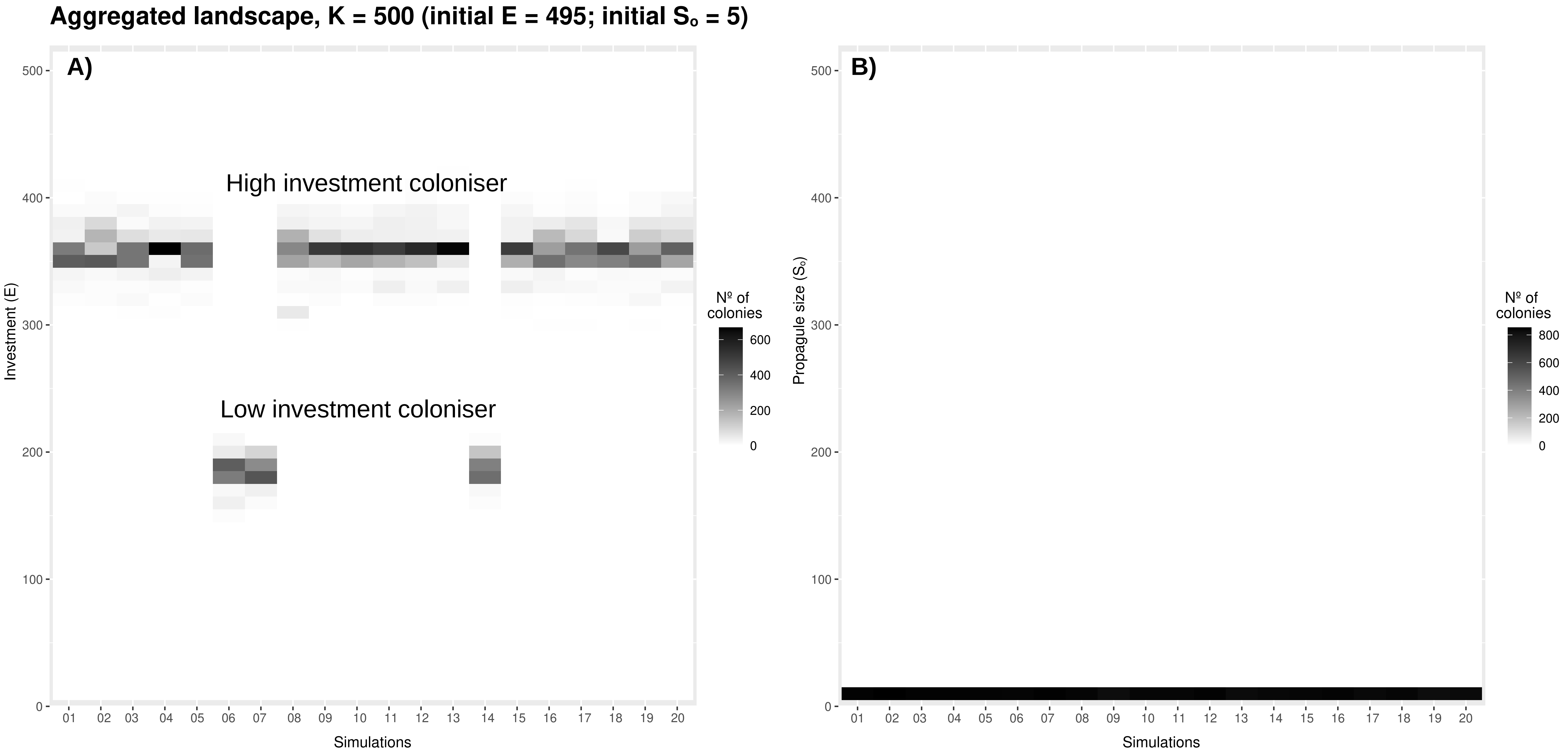


**Figure S1.** Example of simulations giving rise to high or low investment coloniser equilibrium strategies, but not both in a given simulation, in Aggregated landscape with K = 500. A) Indicates the distribution of final investment of colonies (~900) within a simulation and B) indicates the distribution of propagule size.
