## Supplementary material for "To disperse or compete? Coevolution of traits leads to a limited number of reproductive strategies": Homogeneous landscape supplementary figure

### Supplementary Figure S2:

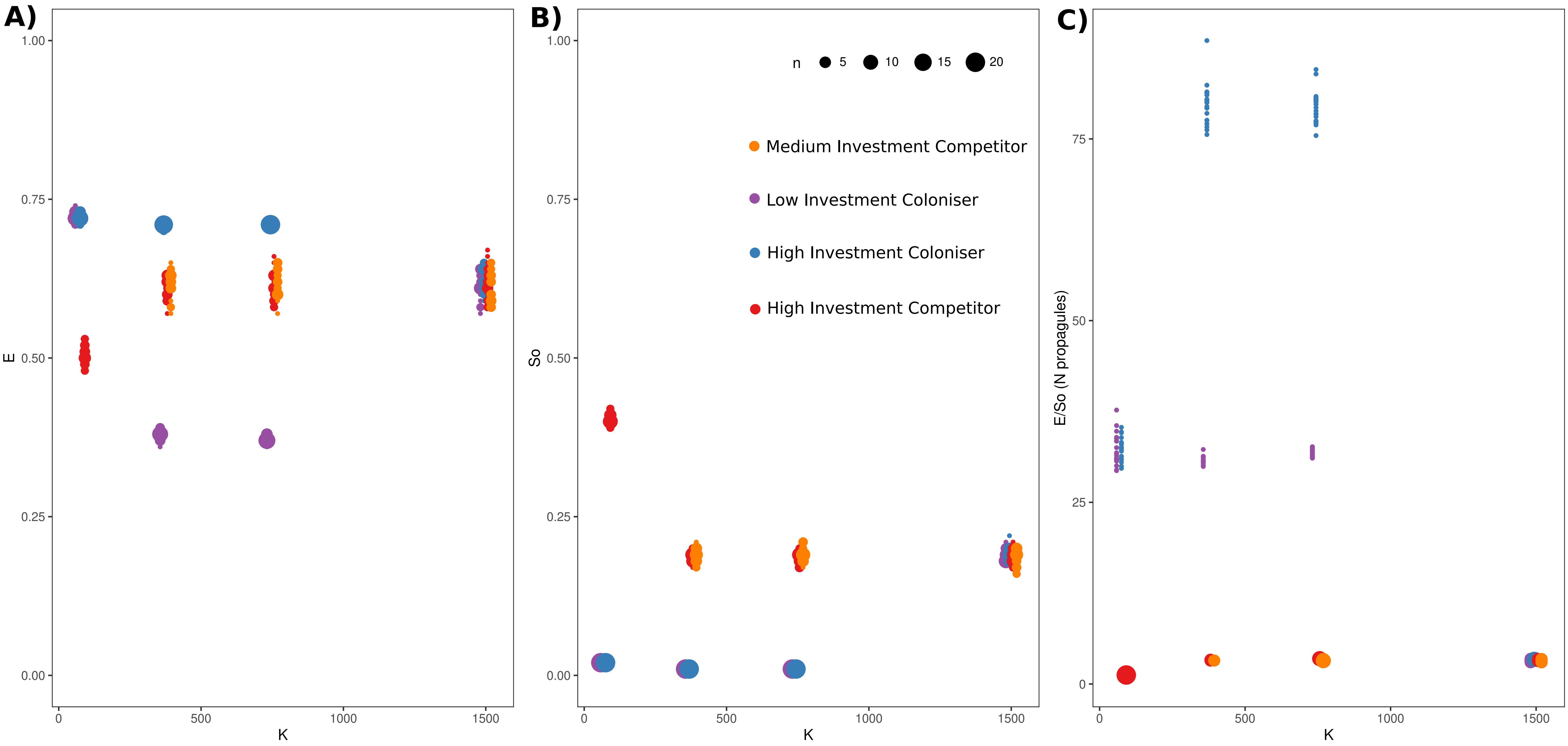


Comparison of final values of a) investment (E), b) propagule size (SO), and c) number of propagules (E/SO) for different resource levels (K) in Homogeneous landscapes. Size of dots corresponds to the number of observations with the same value, while colour indicates the starting condition.
Simulations performed with strong size-dispersal trade-off and fast mutation rate (0.05).
