## Supplementary material for "To disperse or compete? Coevolution of traits leads to a limited number of reproductive strategies": Rasters used: Coev-traits_maps.pdf

### Aggregated landscape nº 1

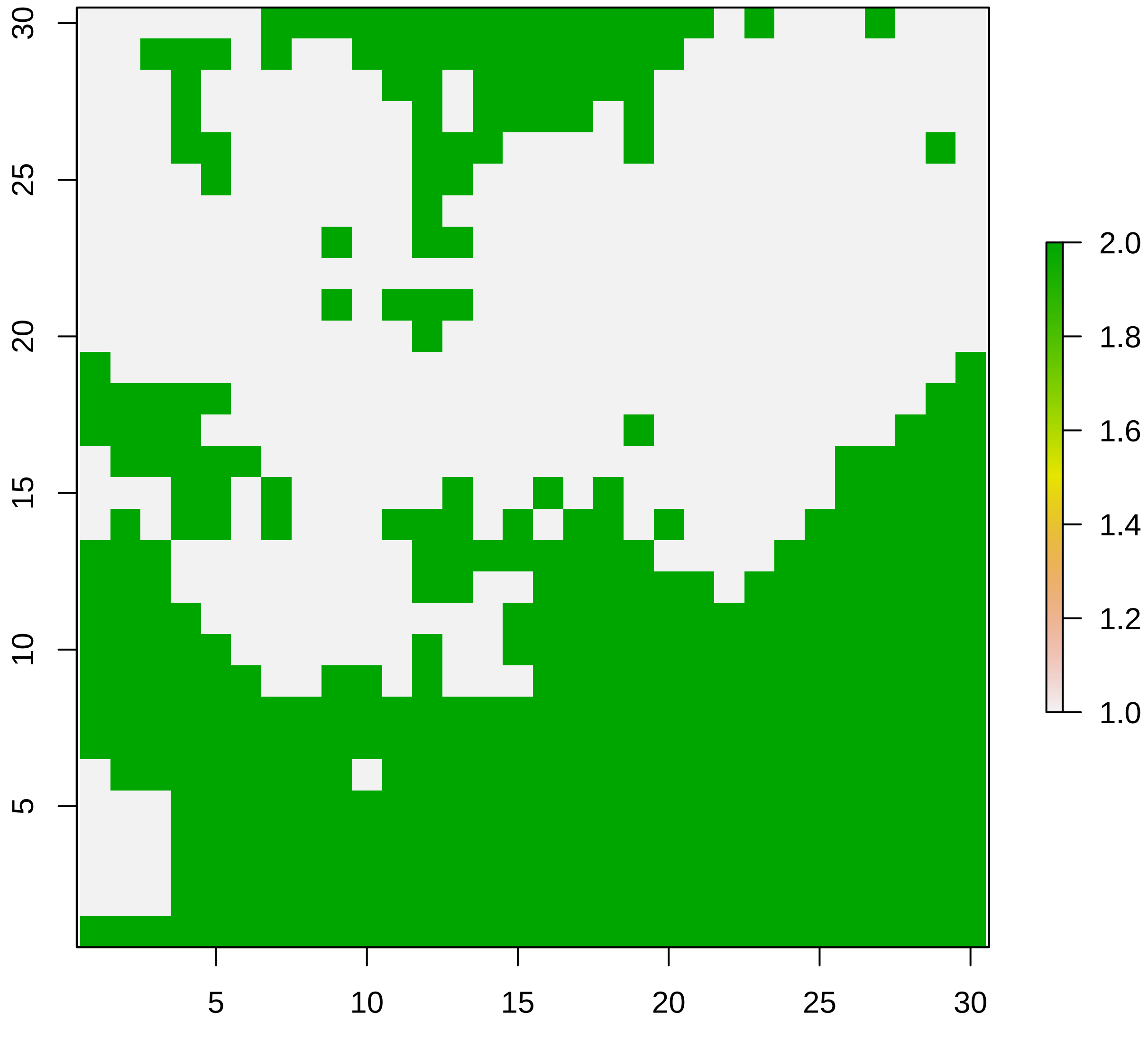

#### Aggregated landscape nº 2

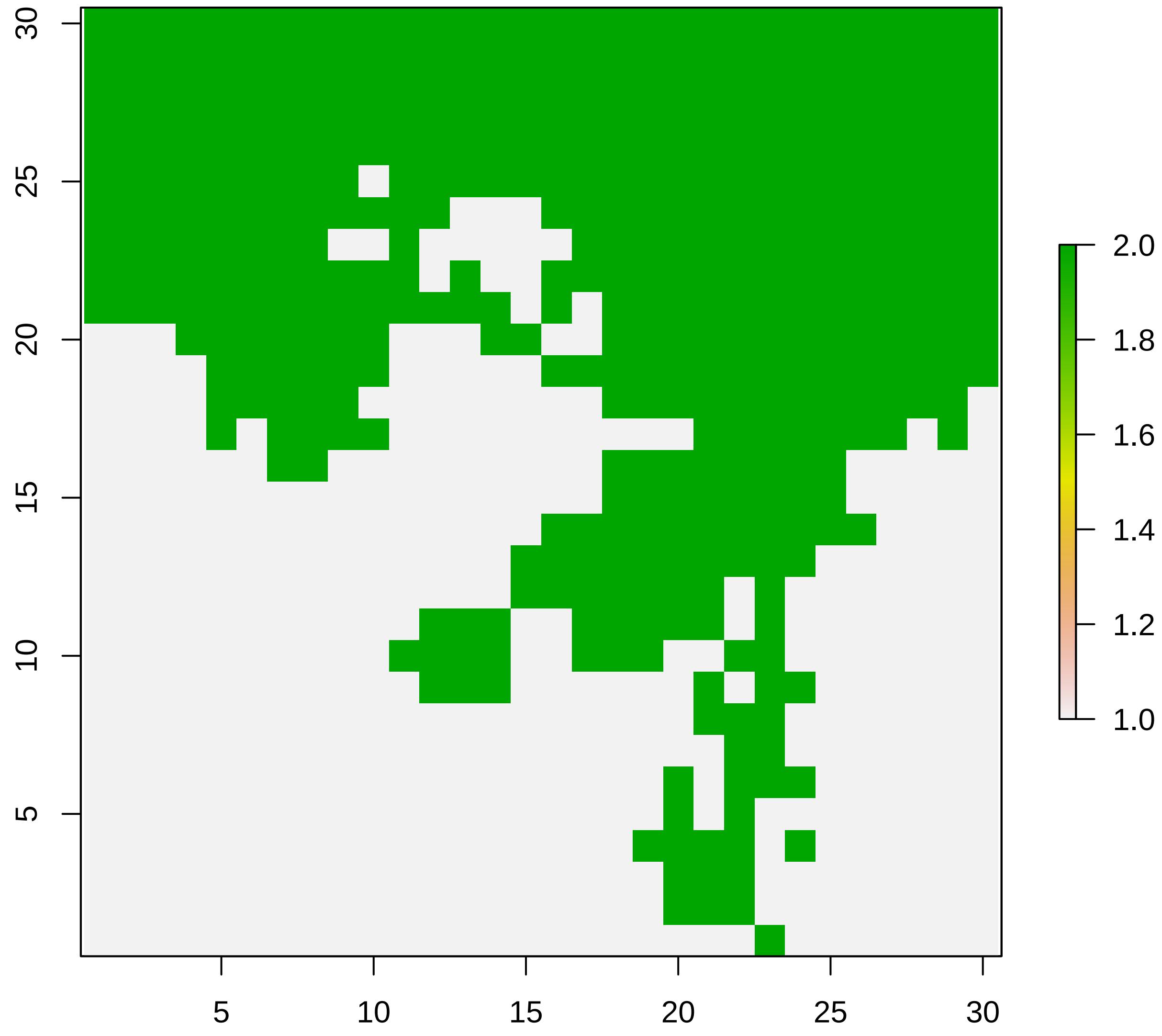

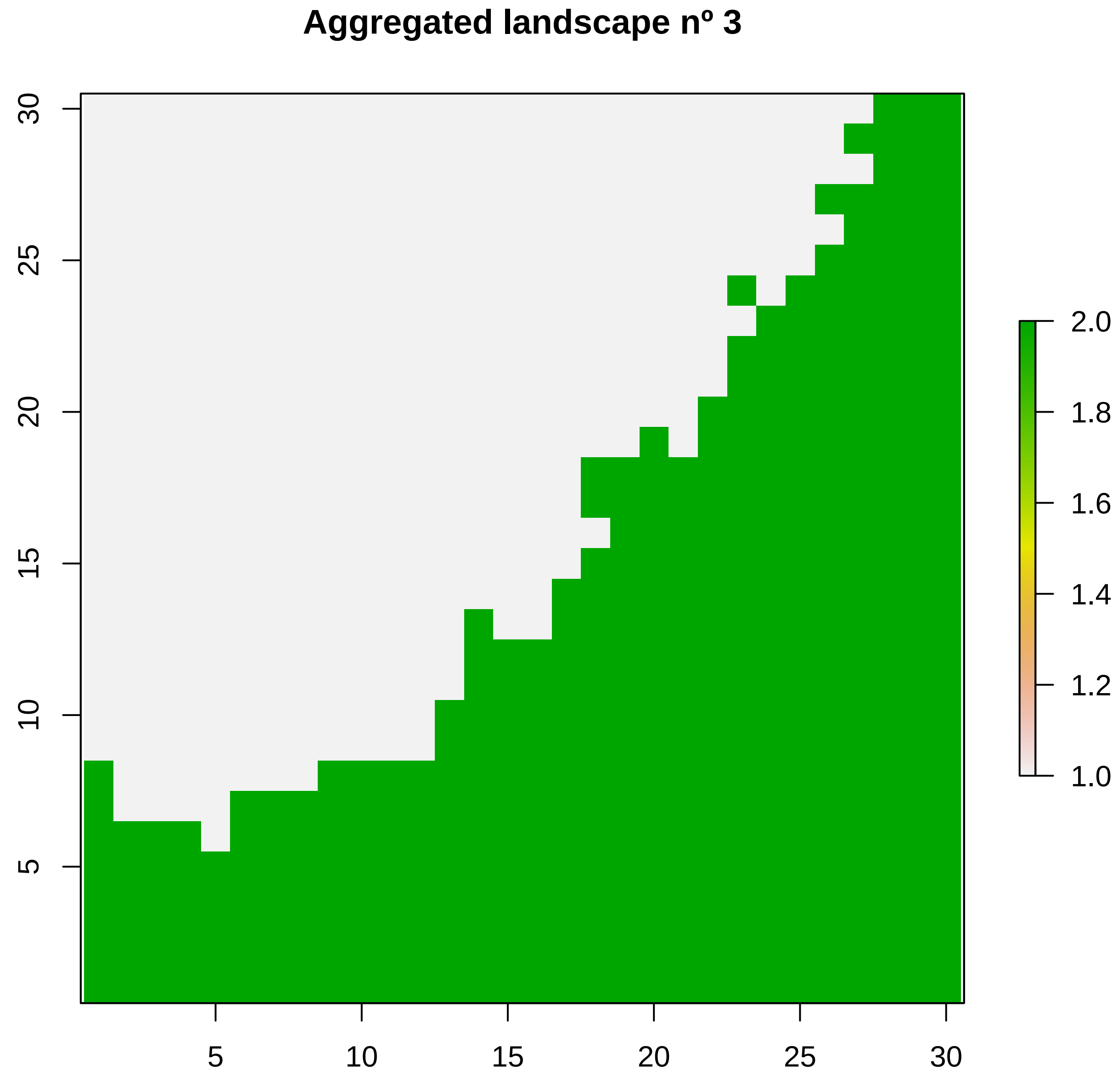

Aggregated landscape nº 3
30
25
20
15
10
5
5
10
15
20
25
30
2.0
1.8
1.6
1.4
1.2
1.0

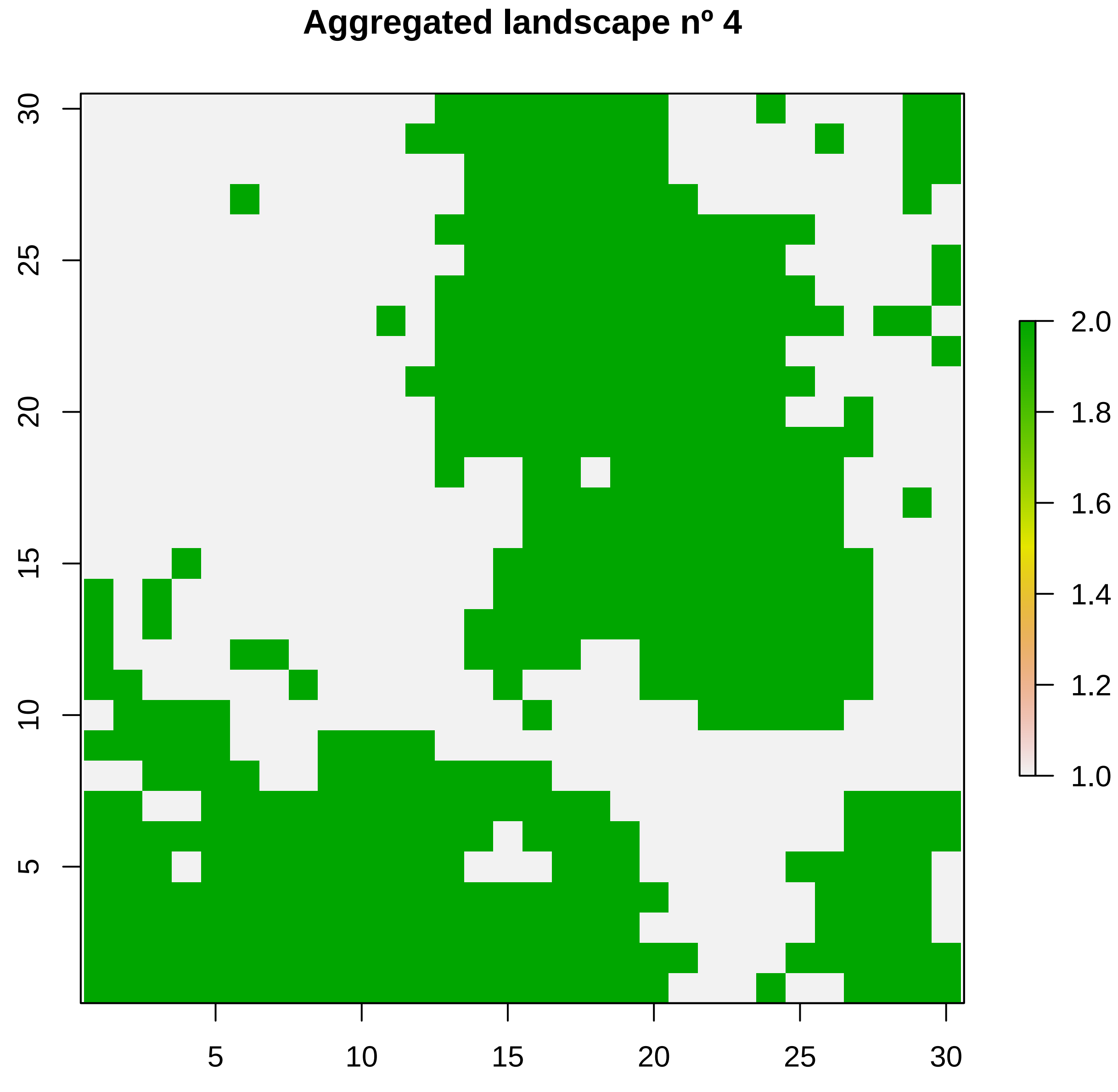
Aggregated landscape nº 4
5
10
15
20
25
30
2.0
1.8
1.6
1.4
1.2
1.0

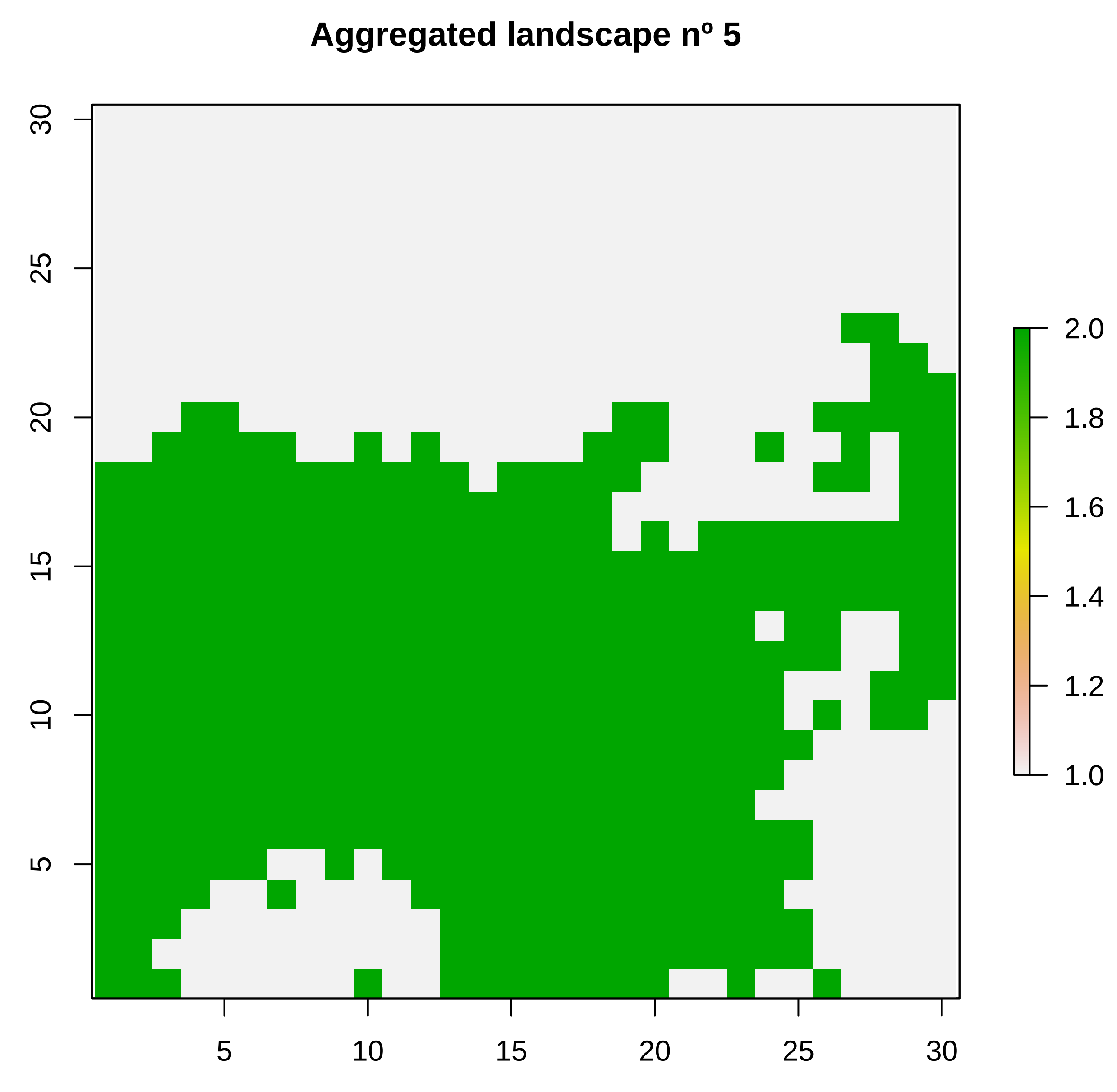

Aggregated landscape nº 5
30
25
20
15
10
5
5
10
15
20
25
30
2.0
1.8
1.6
1.4
1.2
1.0

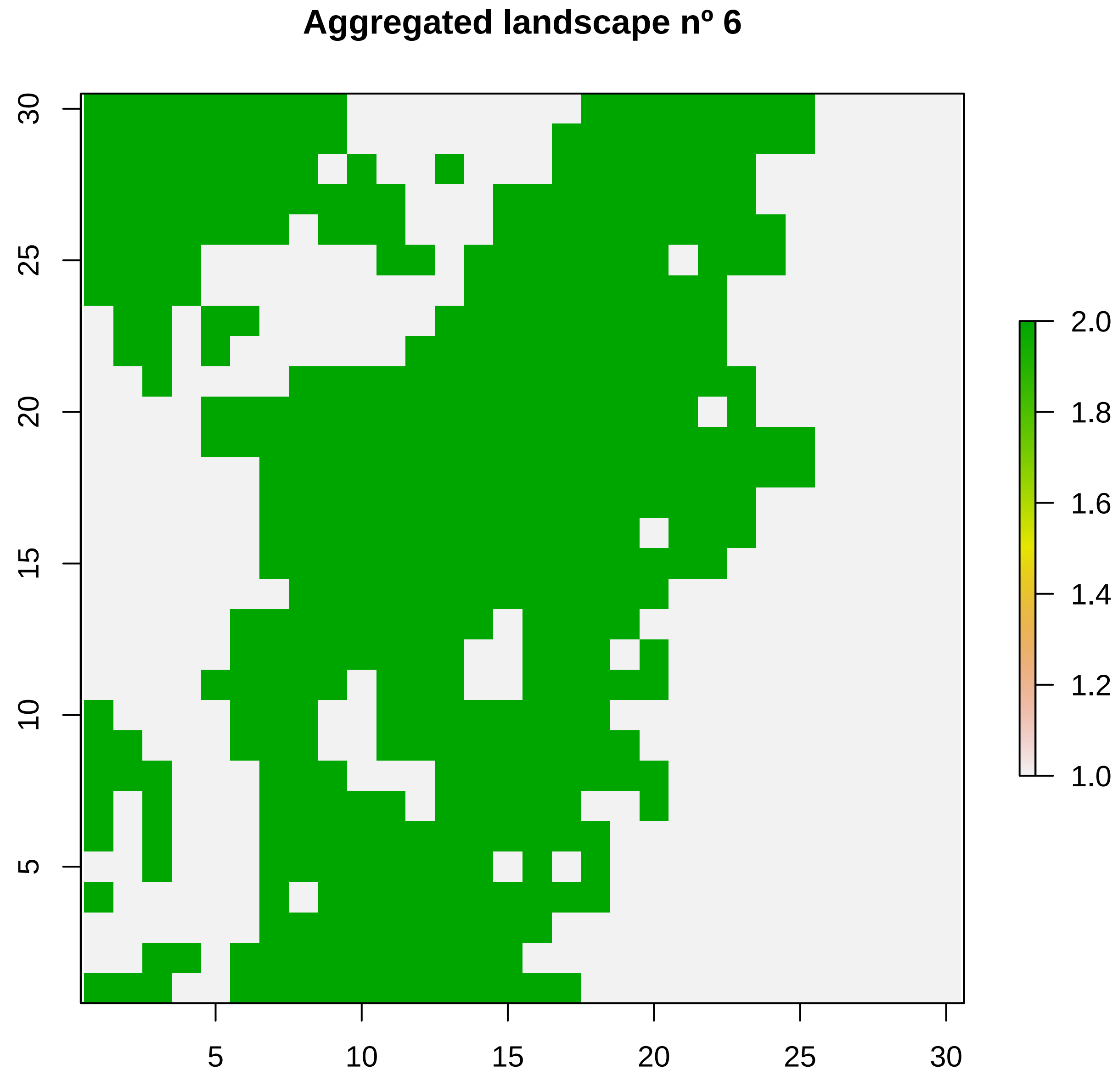

Aggregated landscape nº 6
5
10
15
20
25
30
5
10
15
20
25
30
2.0
1.8
1.6
1.4
1.2
1.0

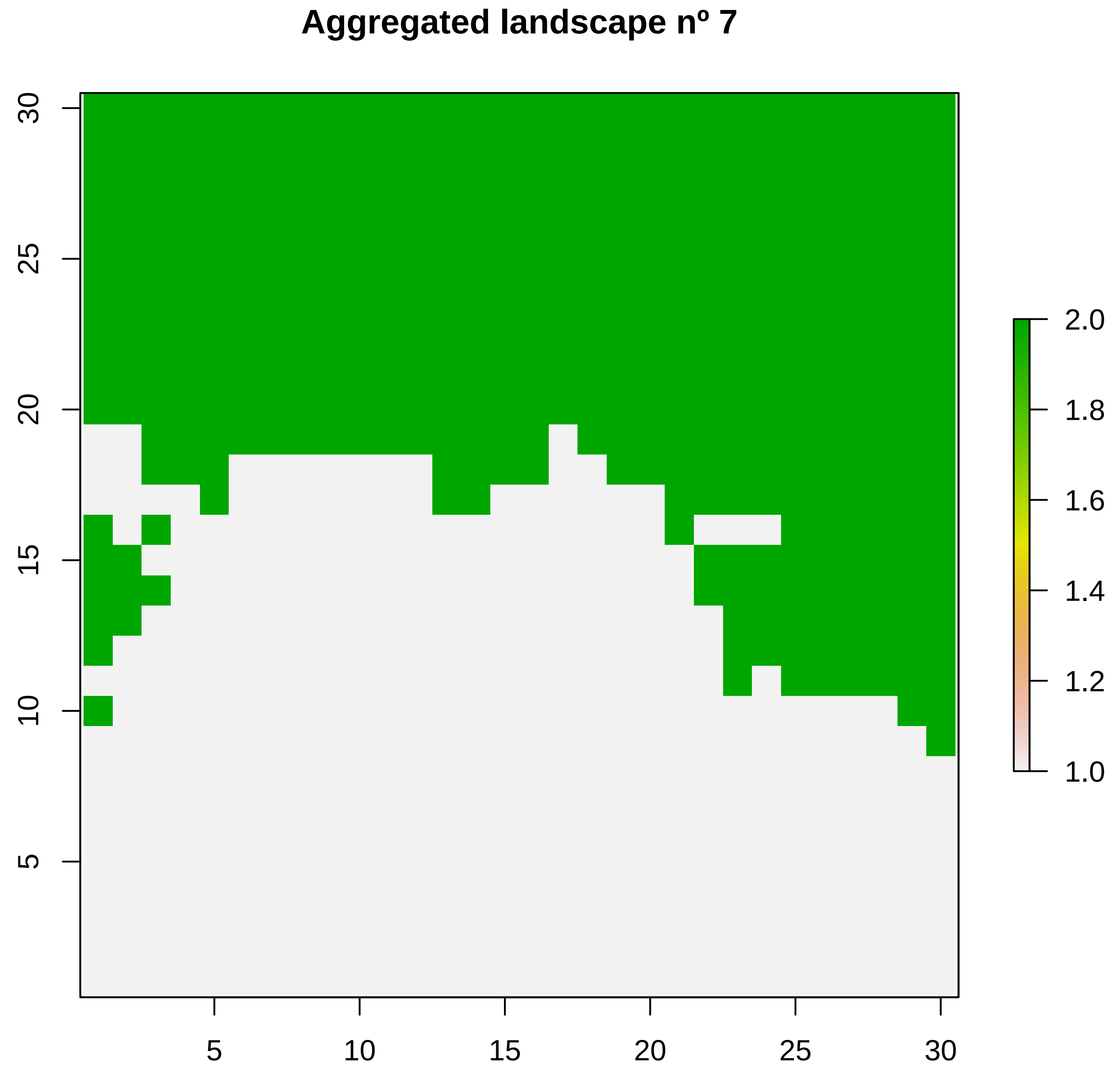

Aggregated landscape nº 7
30
25
20
15
10
5
5
10
15
20
25
30
2.0
1.8
1.6
1.4
1.2
1.0

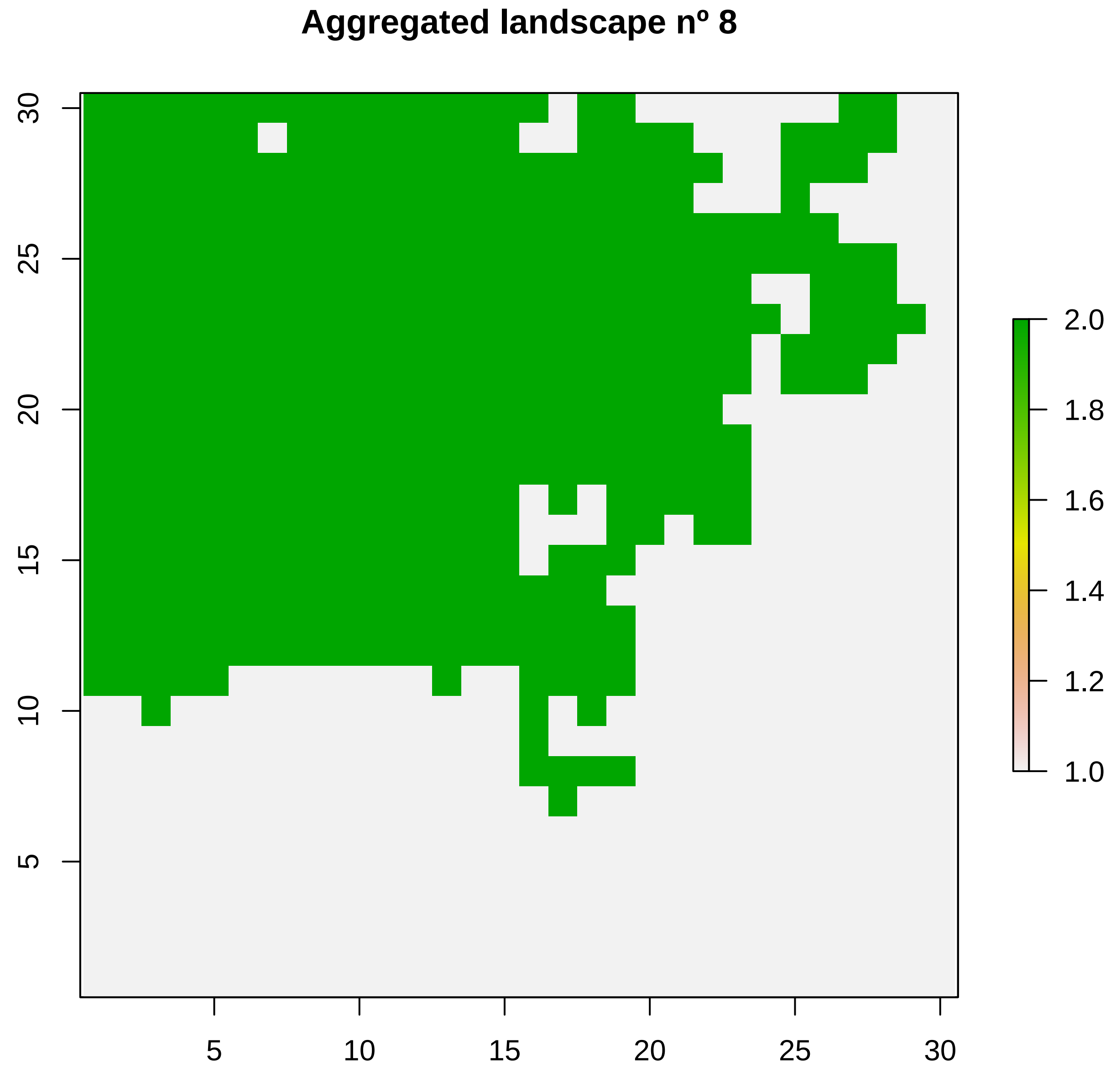

Aggregated landscape nº 8
30
25
20
15
10
5
5
10
15
20
25
30
2.0
1.8
1.6
1.4
1.2
1.0

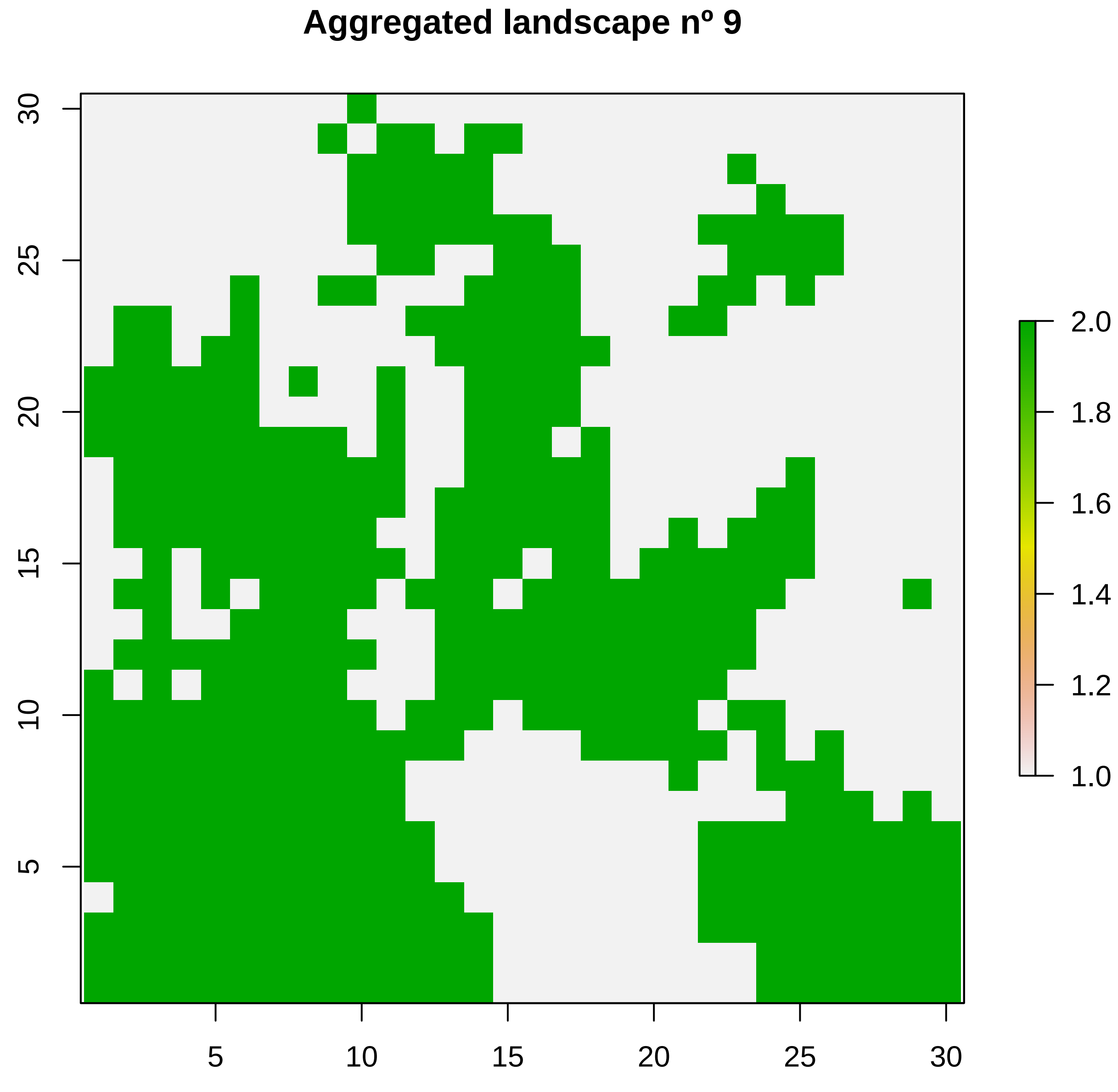
Aggregated landscape nº 9
5
10
15
20
25
30
5
10
15
20
25
30
1.0
1.2
1.4
1.6
1.8
2.0

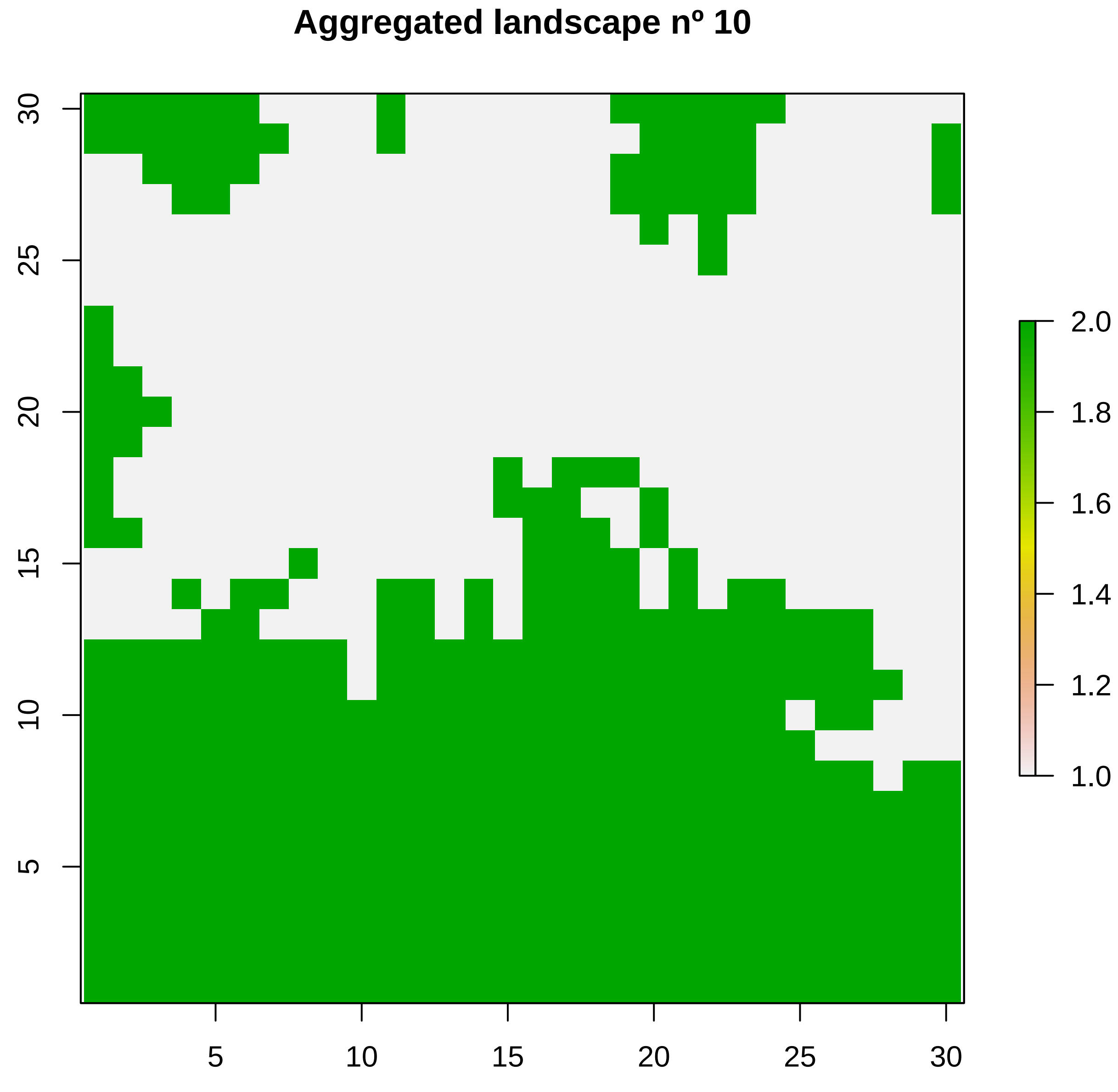

Aggregated landscape nº 10
5
10
15
20
25
30
5
10
15
20
25
30
2.0
1.8
1.6
1.4
1.2
1.0

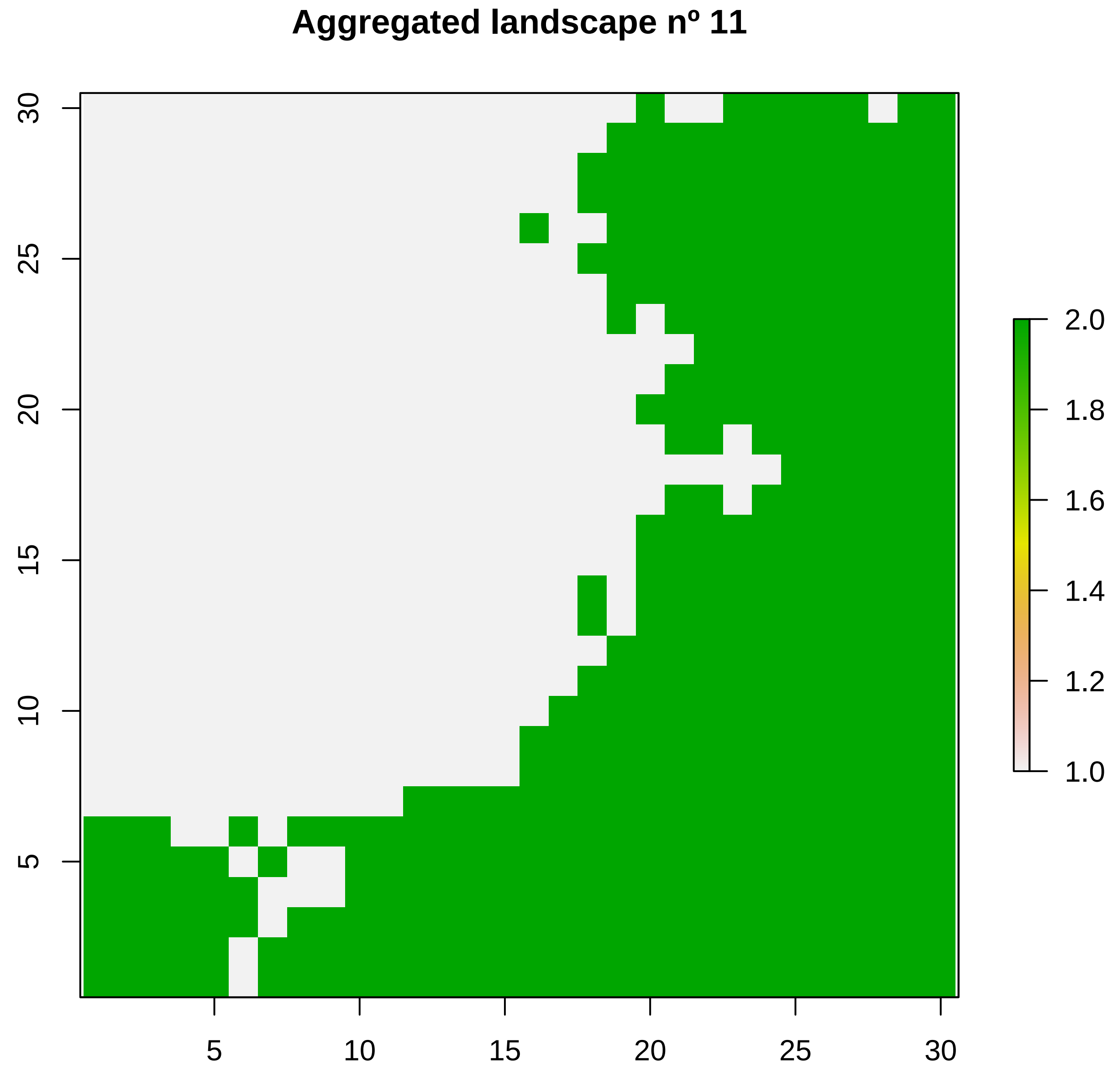
Aggregated landscape nº 11
5
10
15
20
25
30
5
10
15
20
25
30
2.0
1.8
1.6
1.4
1.2
1.0

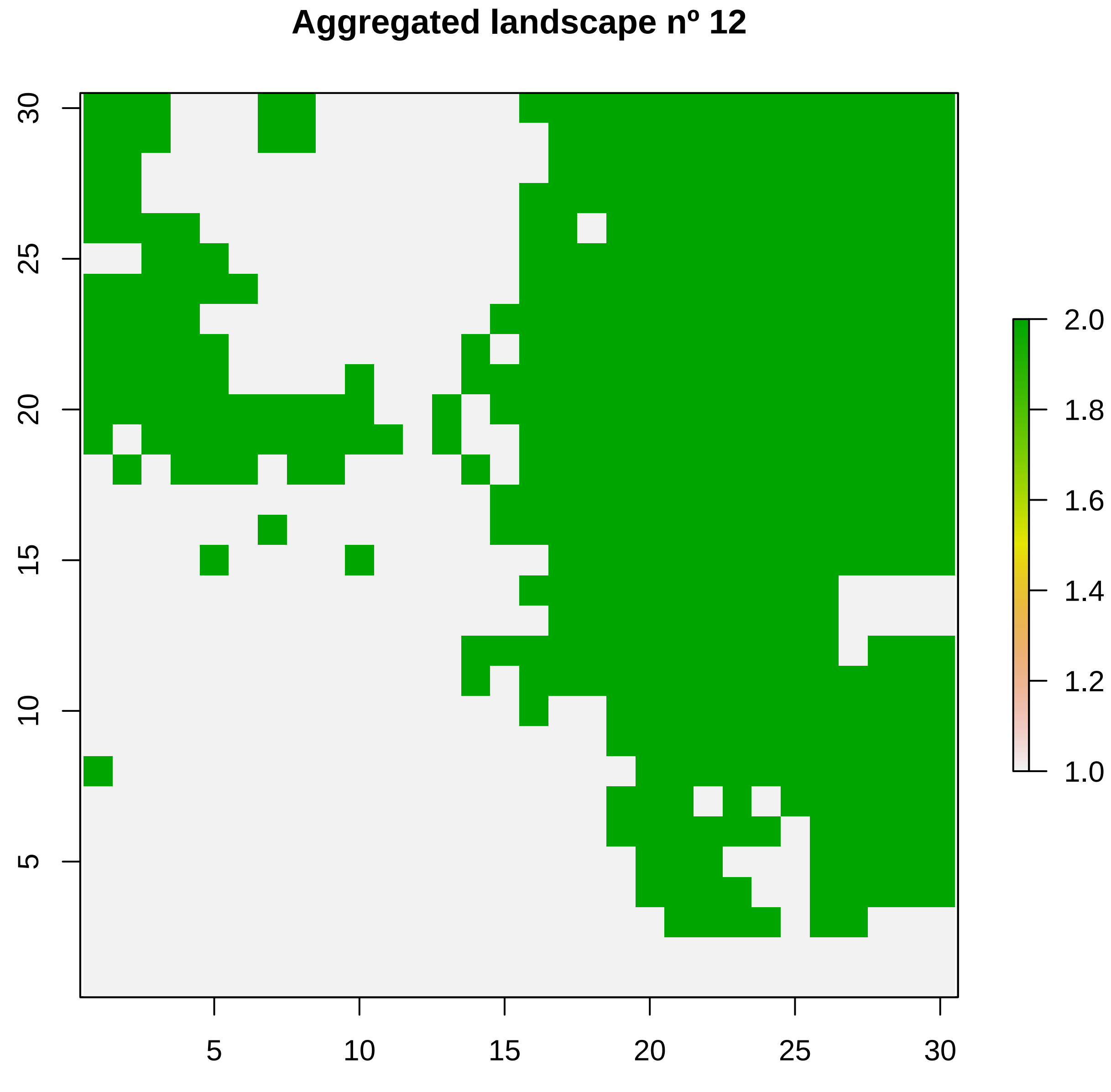
Aggregated landscape nº 12
30
25
20
15
10
5
5
10
15
20
25
30
2.0
1.8
1.6
1.4
1.2
1.0

**Aggregated landscape nº 13**

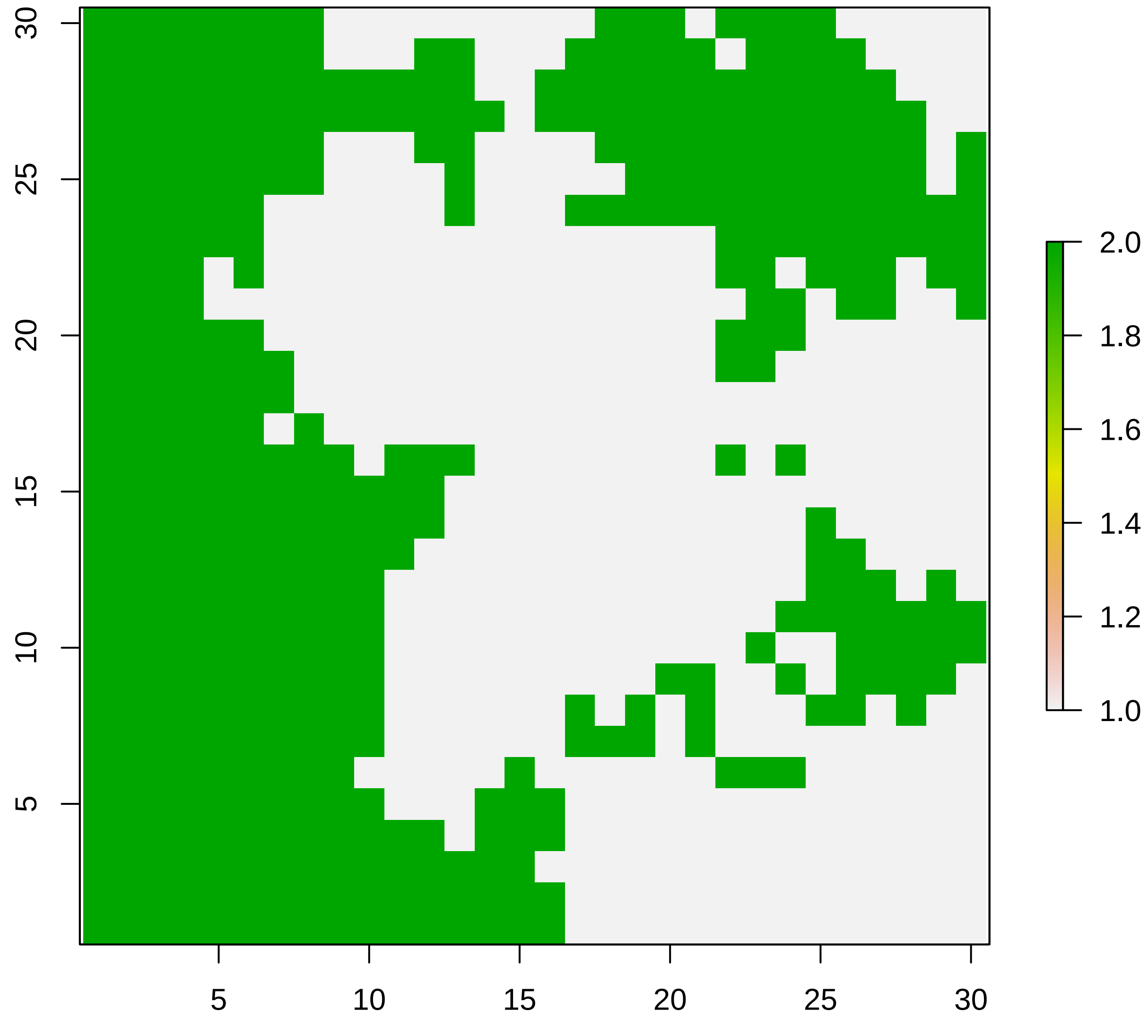

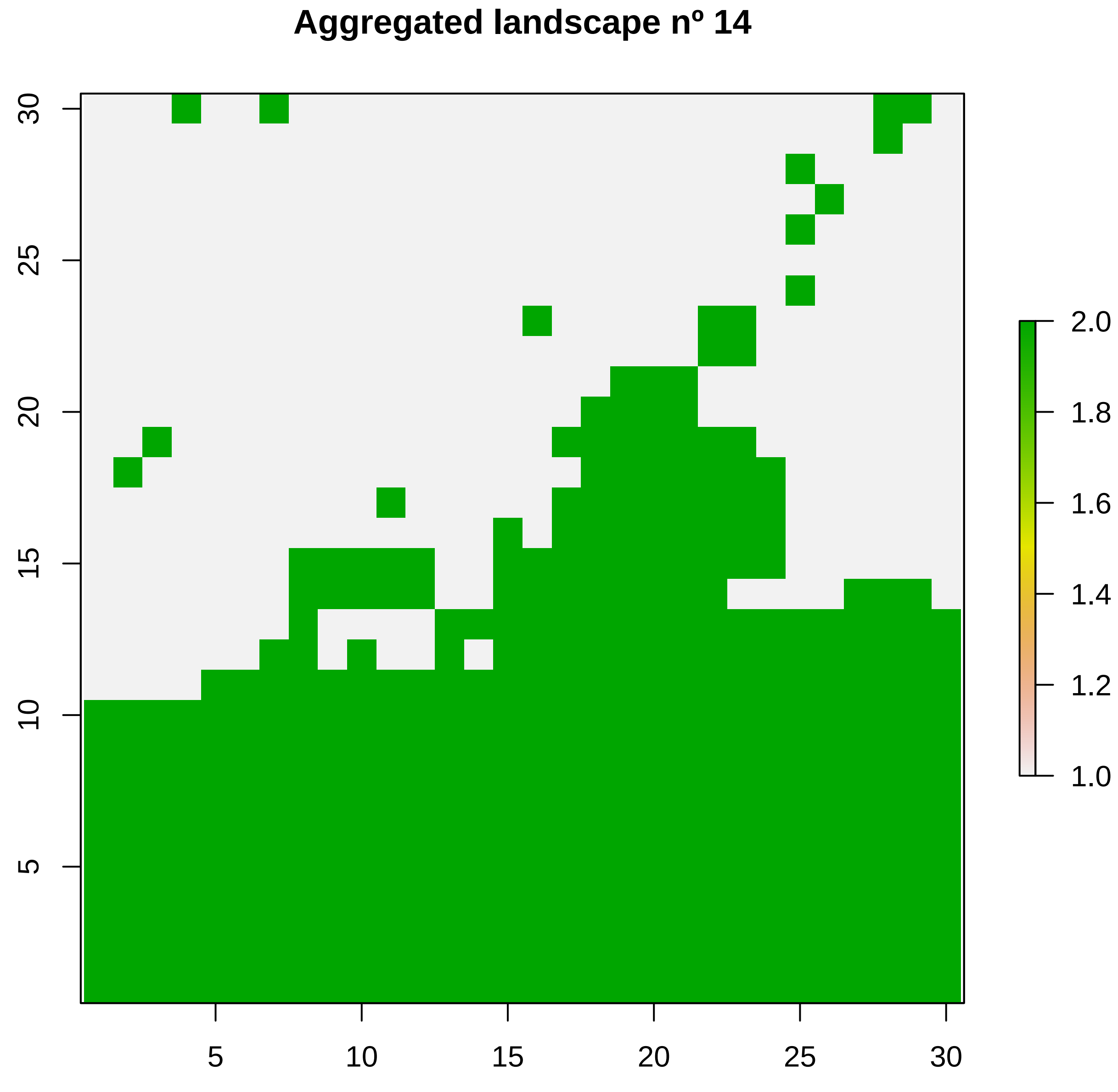
Aggregated landscape nº 14
5
10
15
20
25
30
5
10
15
20
25
30
1.0
1.2
1.4
1.6
1.8
2.0

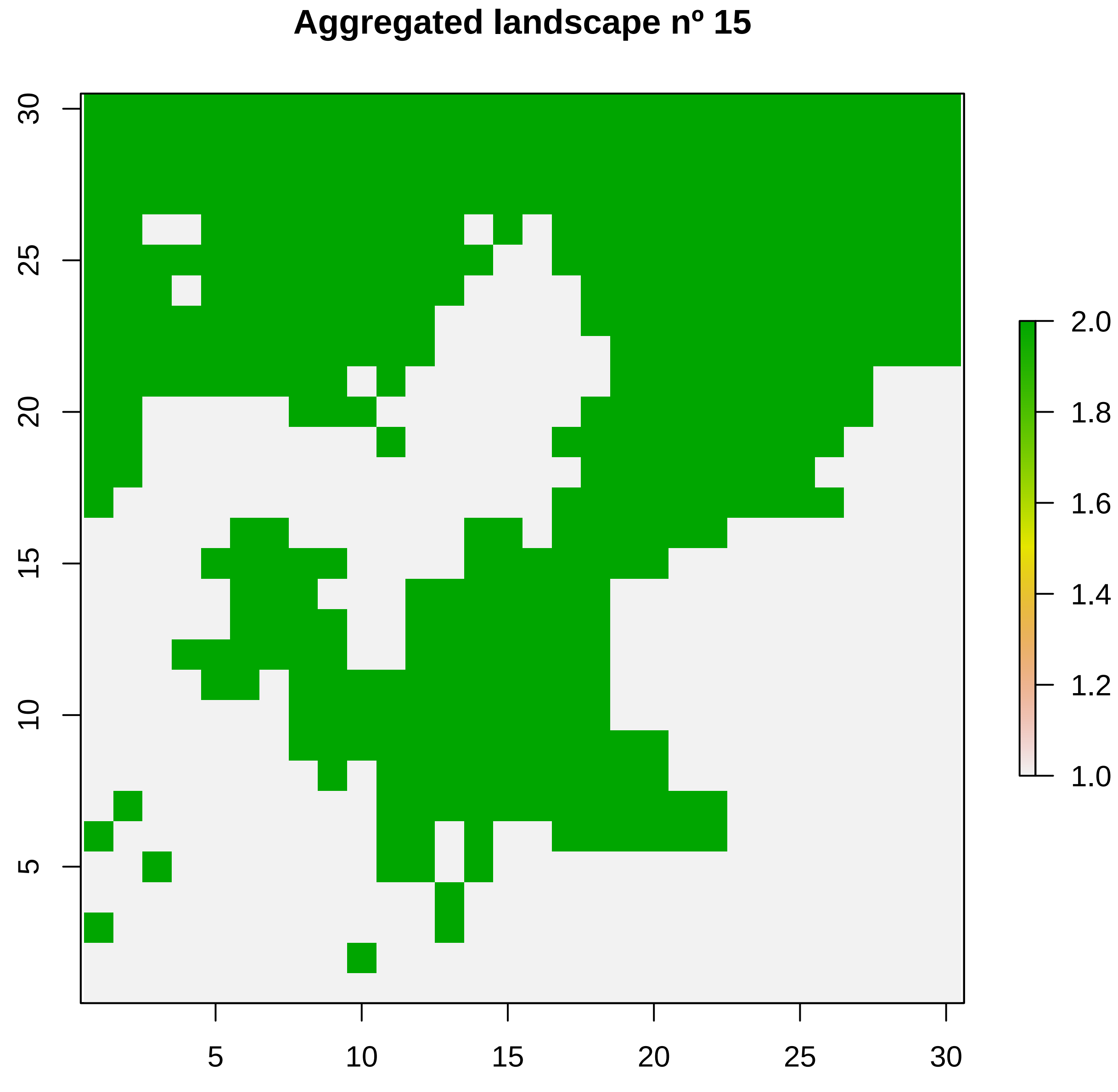
Aggregated landscape nº 15
5
10
15
20
25
30
5
10
15
20
25
30
2.0
1.8
1.6
1.4
1.2
1.0

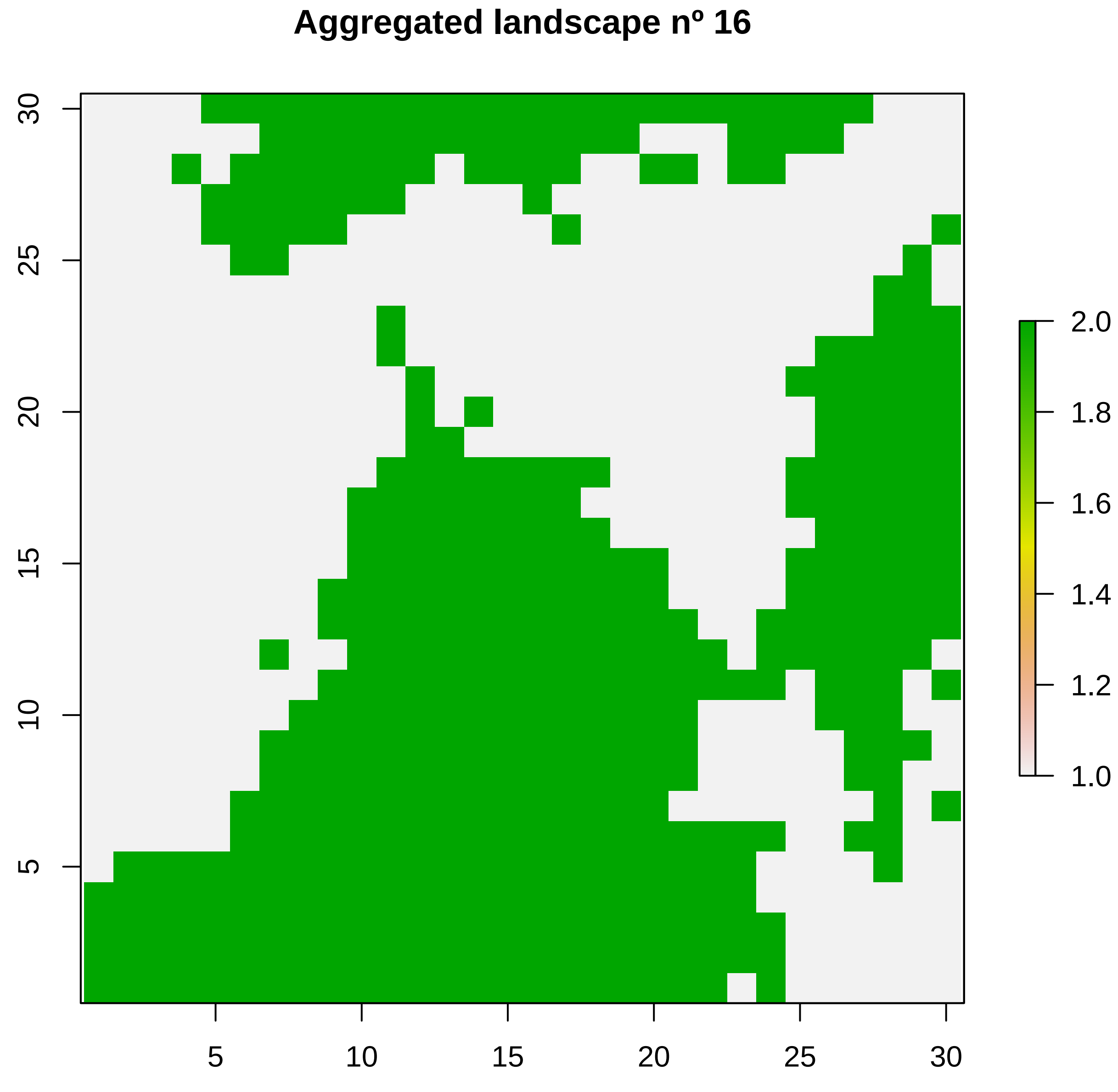

Aggregated landscape nº 16
30
25
20
15
10
5
5
10
15
20
25
30
2.0
1.8
1.6
1.4
1.2
1.0

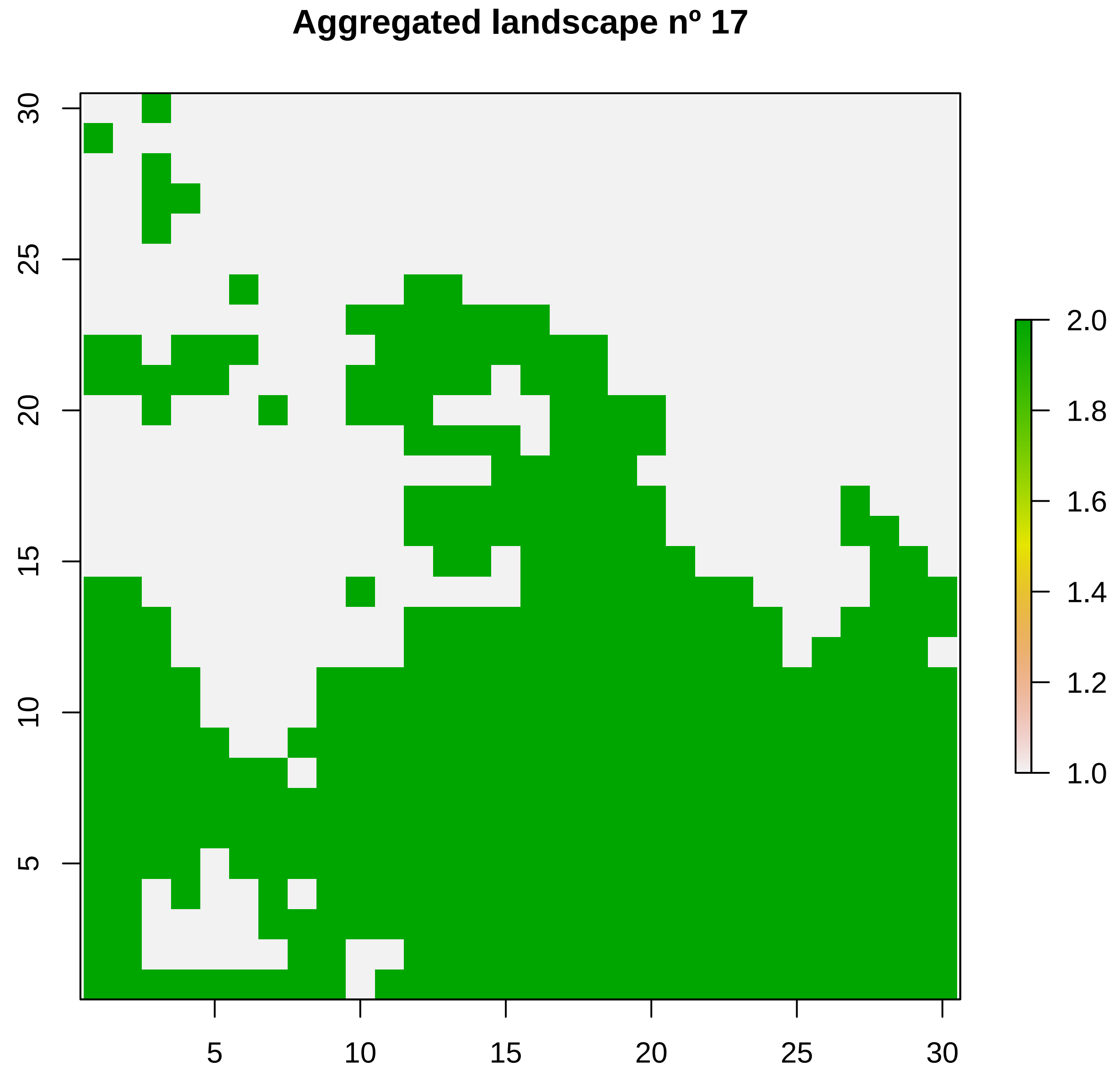
Aggregated landscape nº 17
5
10
15
20
25
30
5
10
15
20
25
30
2.0
1.8
1.6
1.4
1.2
1.0

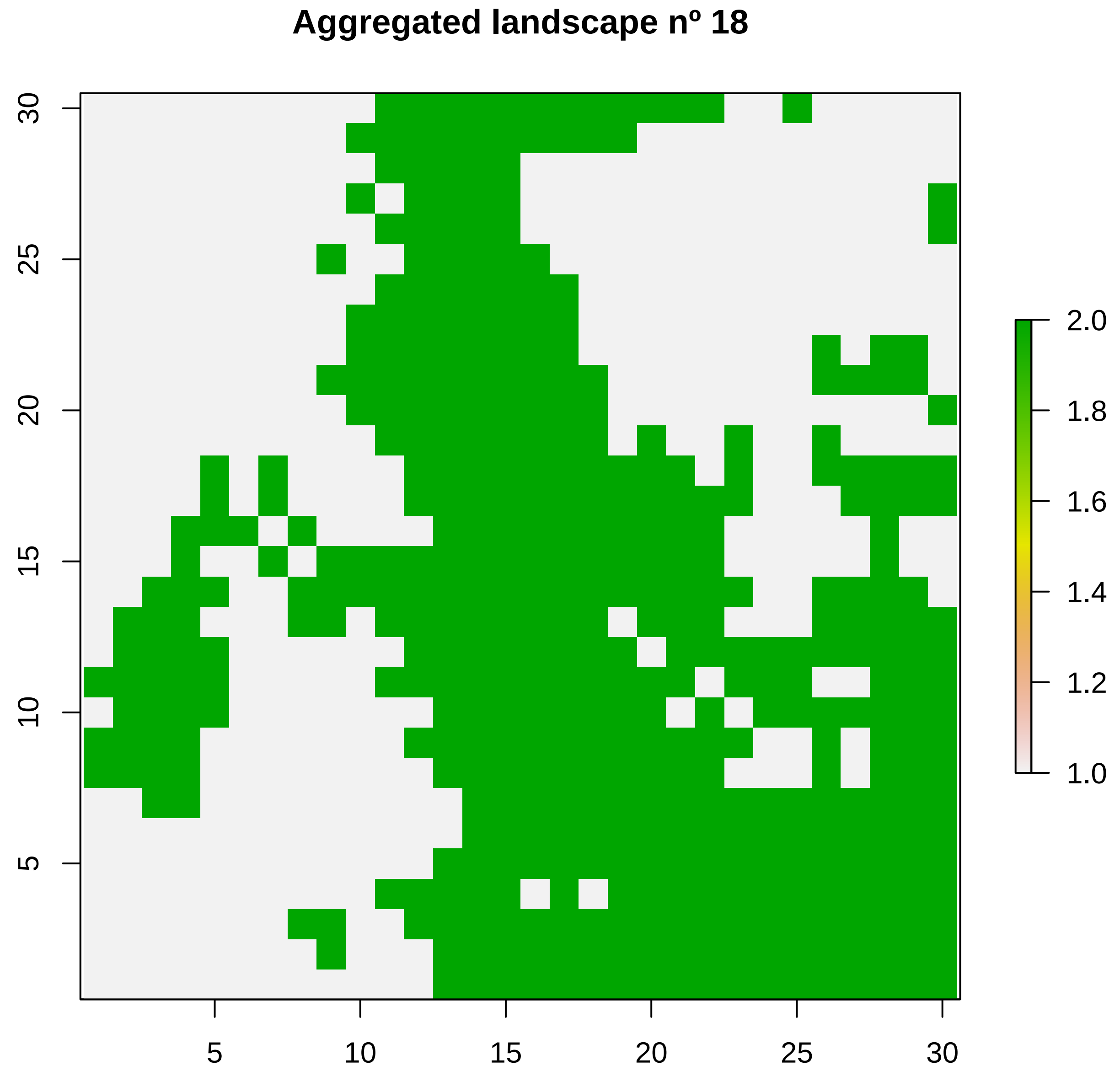
Aggregated landscape nº 18
5
10
15
20
25
30
5
10
15
20
25
30
1.0
1.2
1.4
1.6
1.8
2.0

### Aggregated landscape nº 19

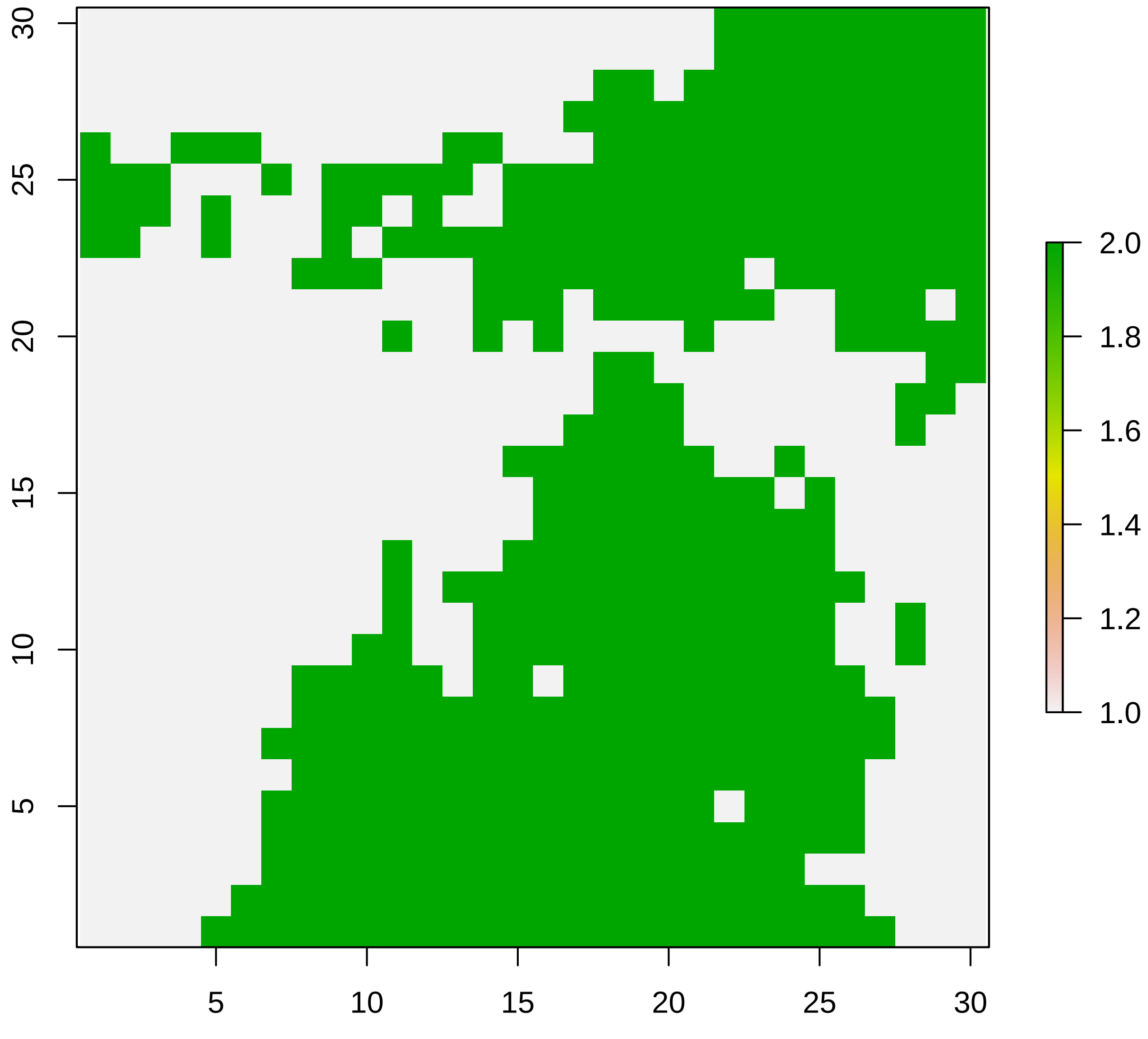

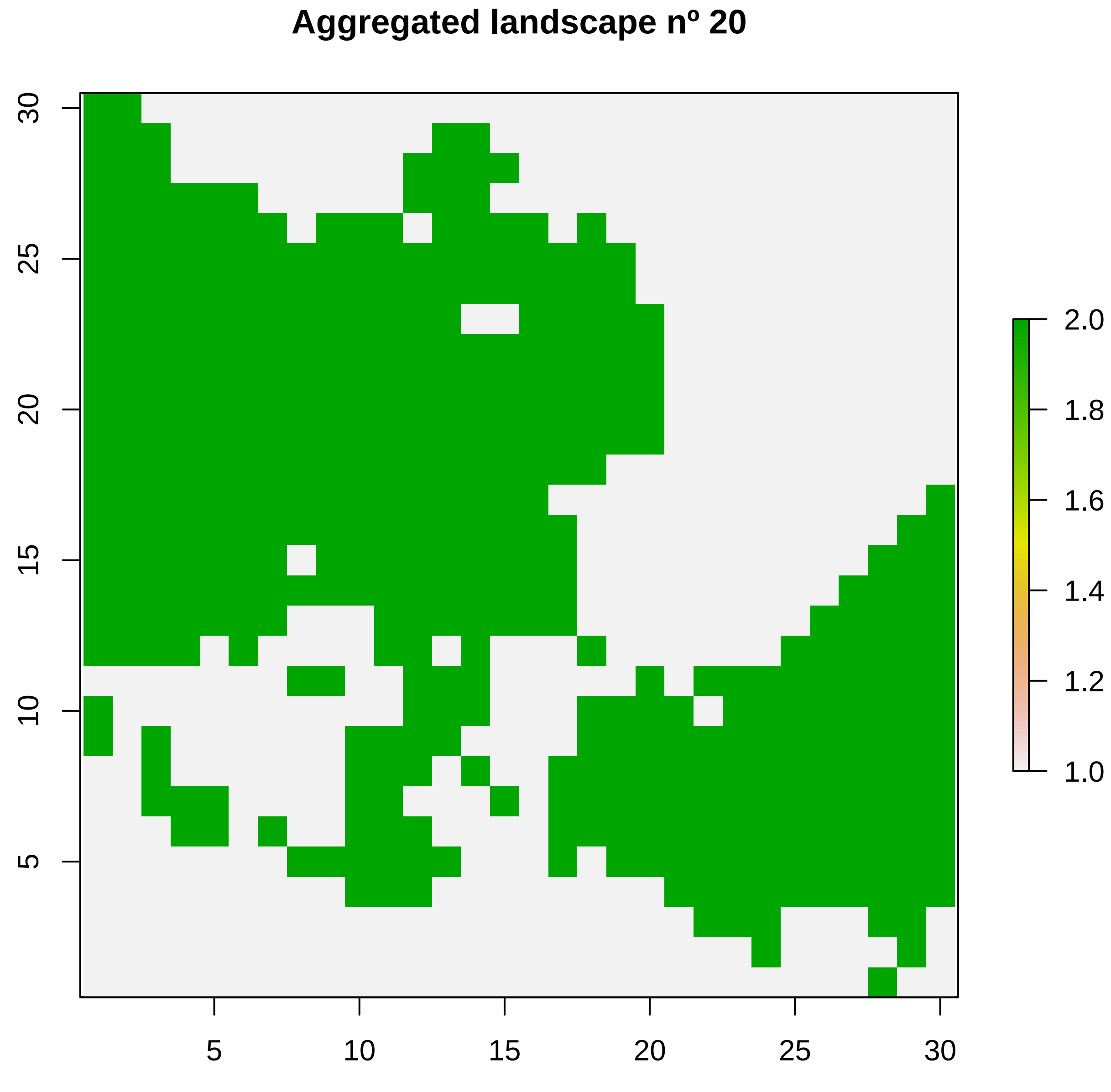
Aggregated landscape nº 20
5
10
15
20
25
30
5
10
15
20
25
30
2.0
1.8
1.6
1.4
1.2
1.0

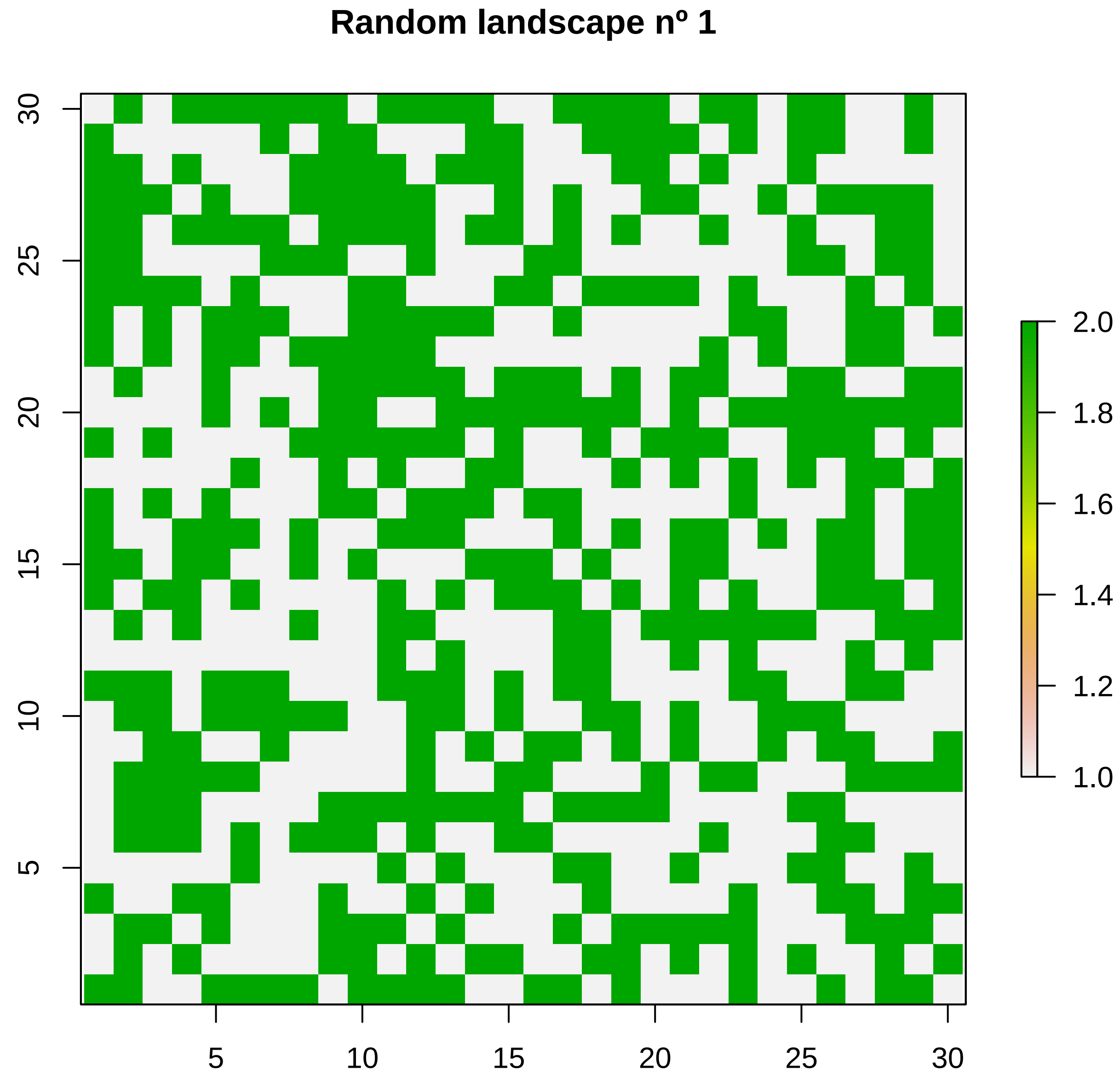
Random landscape nº 1
5
10
15
20
25
30
5
10
15
20
25
30
2.0
1.8
1.6
1.4
1.2
1.0

**Random landscape nº 2**

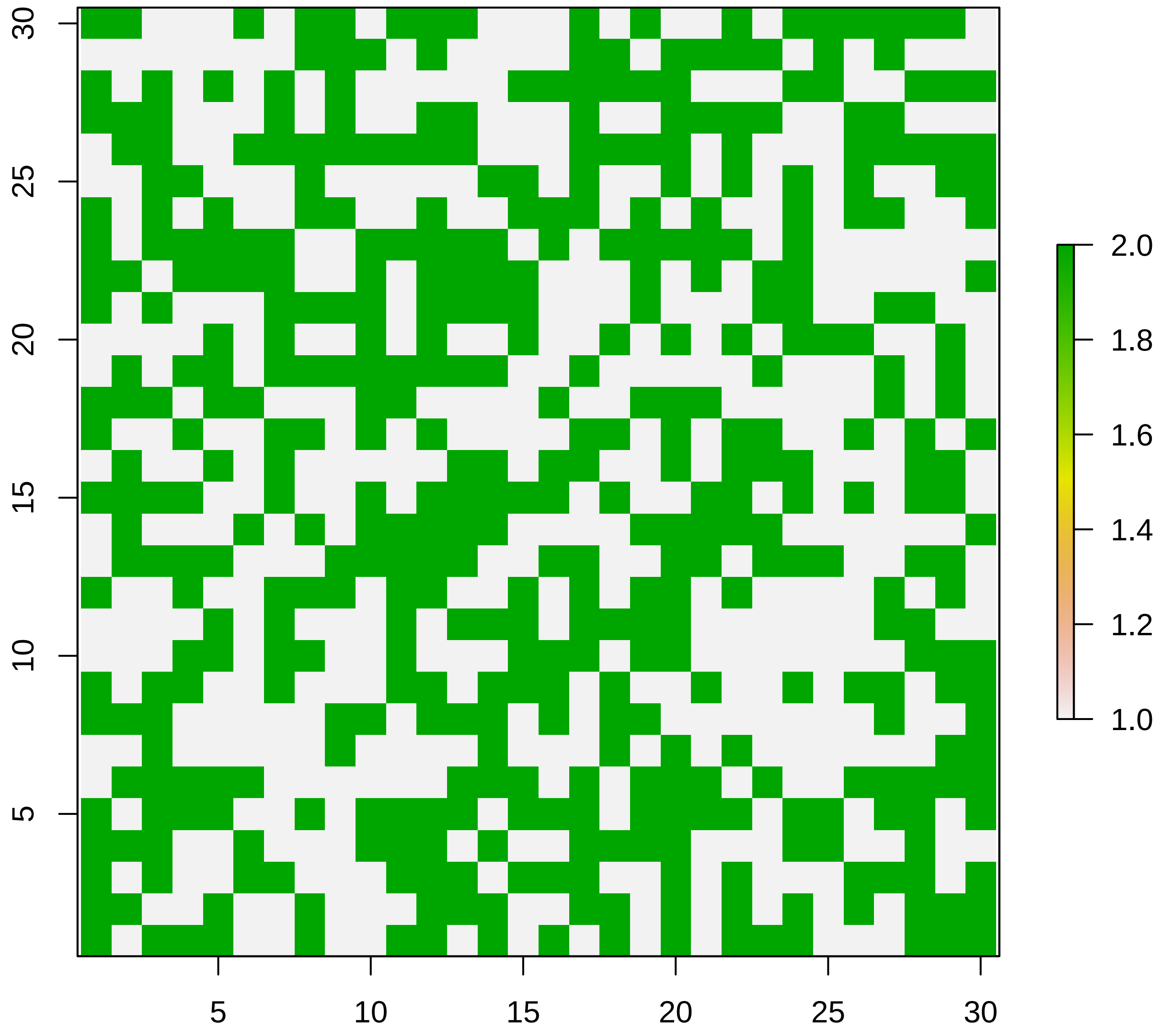

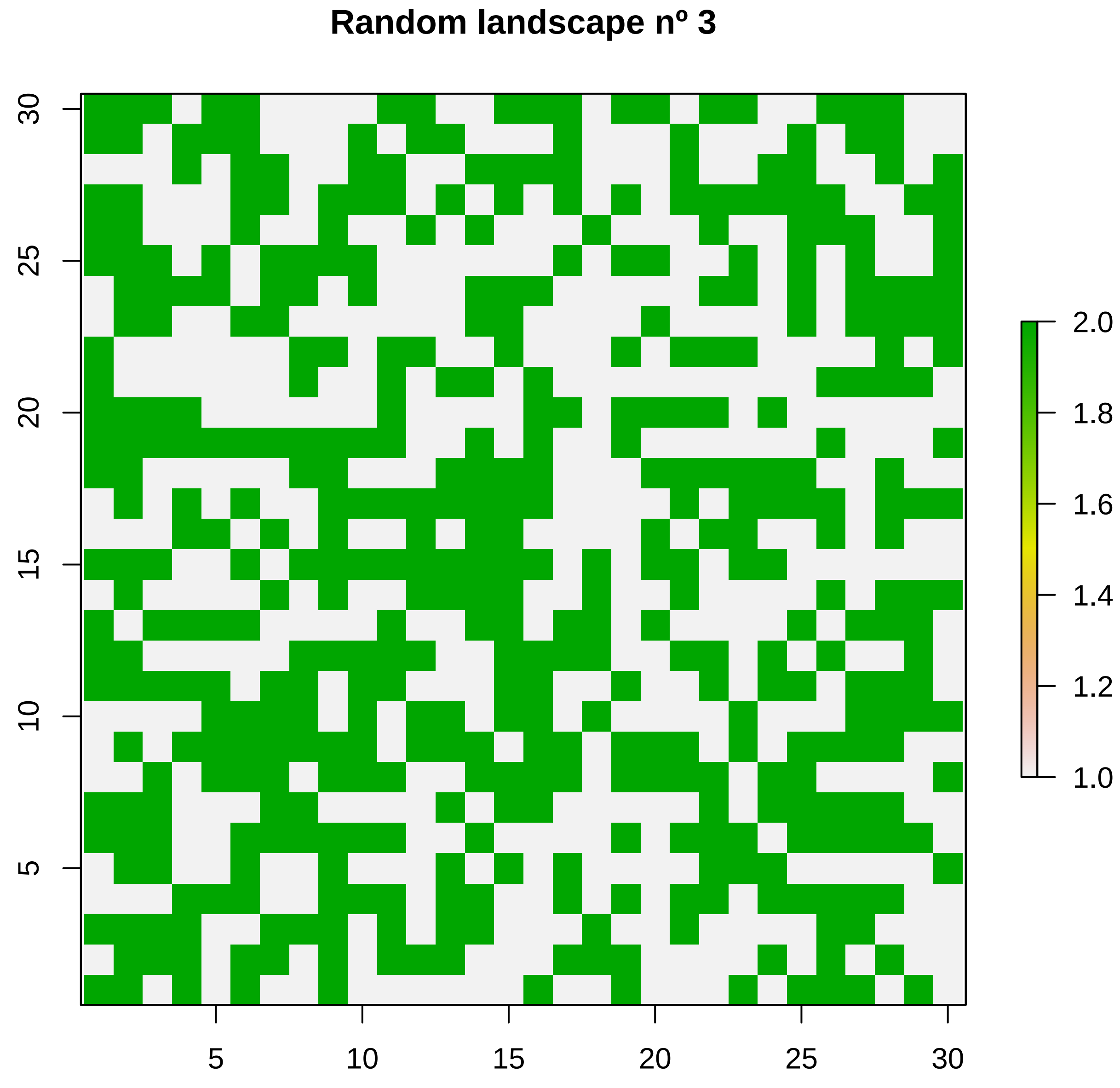
Random landscape nº 3
5
10
15
20
25
30
5
10
15
20
25
30
1.0
1.2
1.4
1.6
1.8
2.0

#### Random landscape nº 4

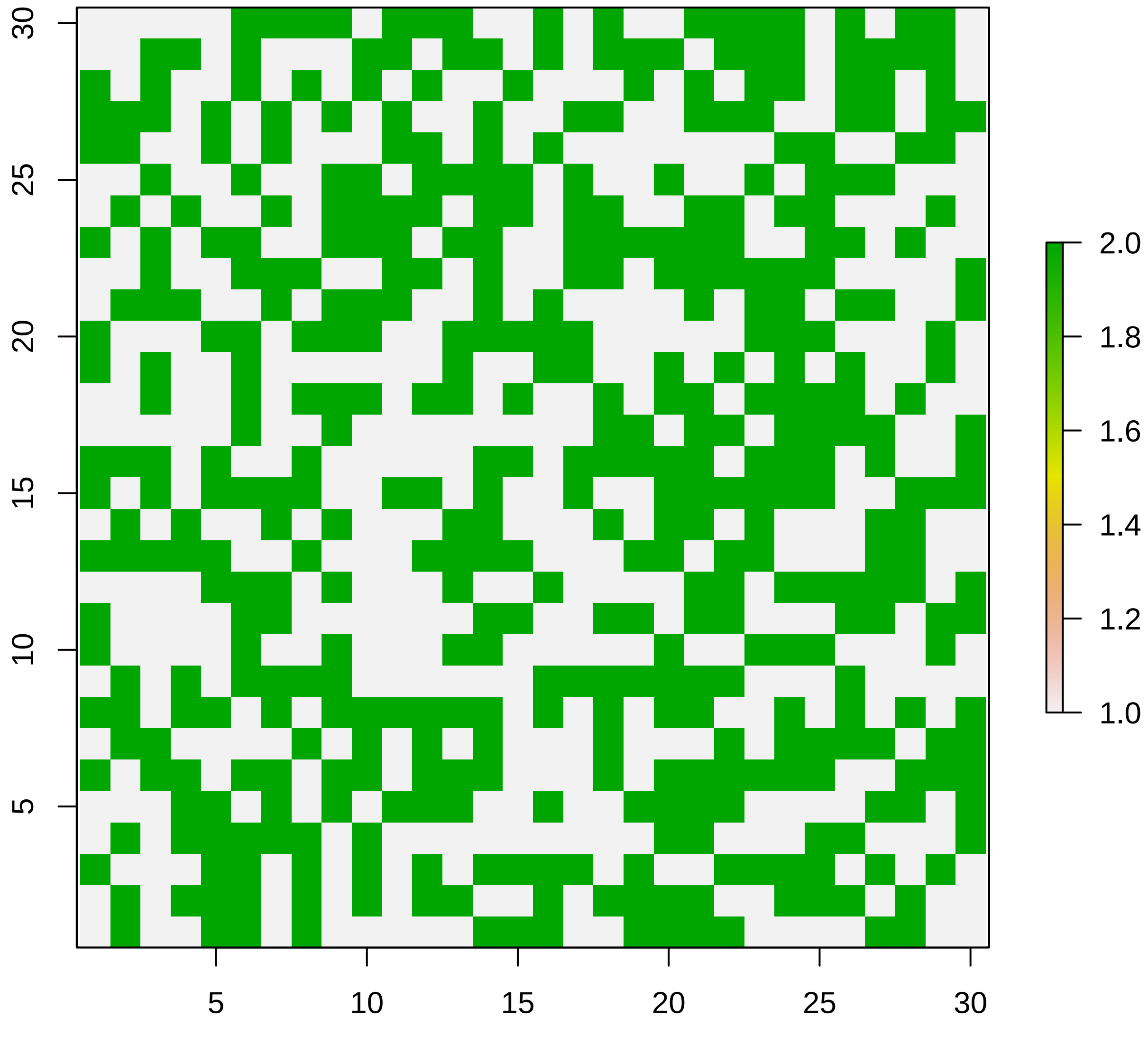

#### Random landscape nº 5

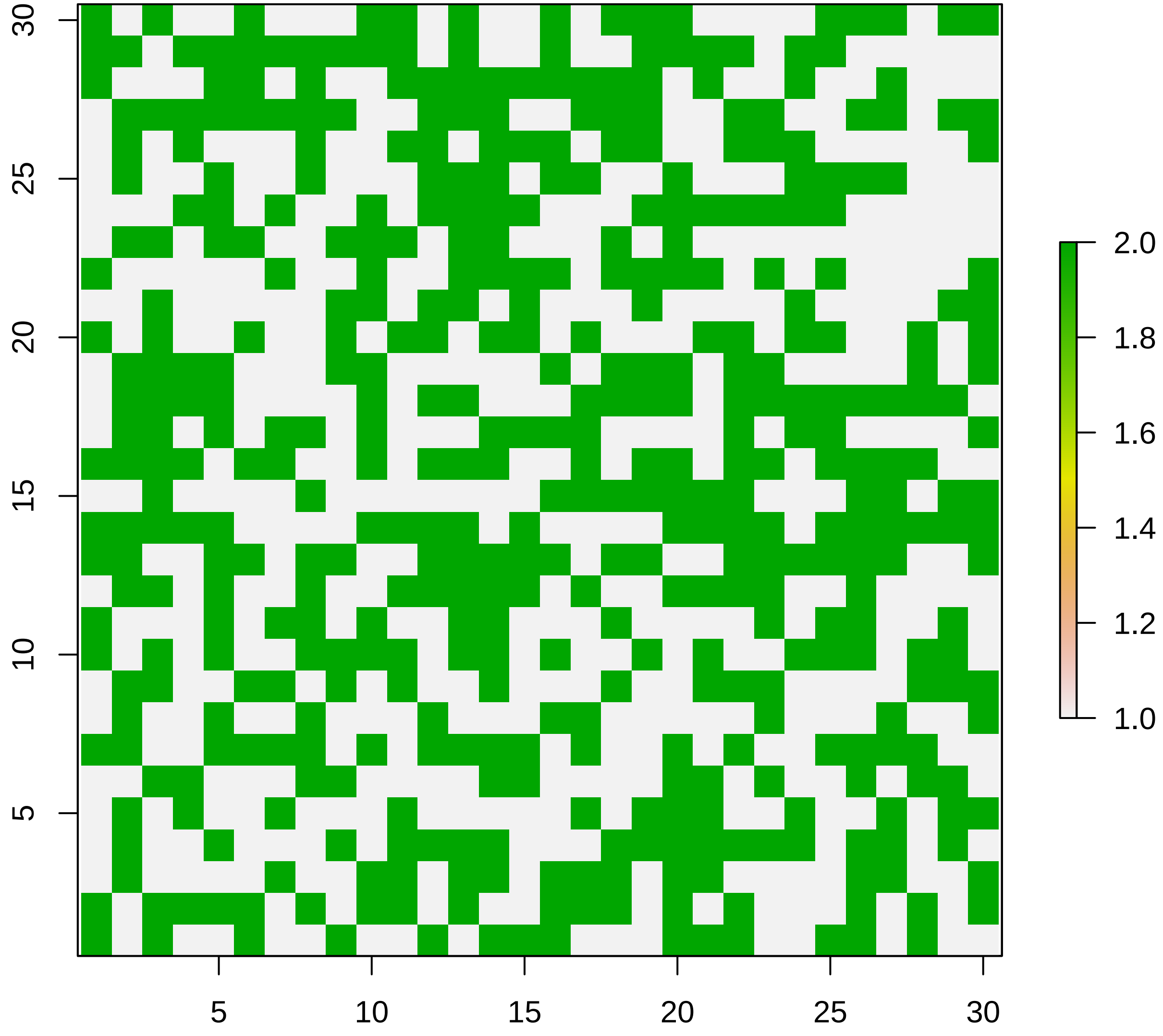

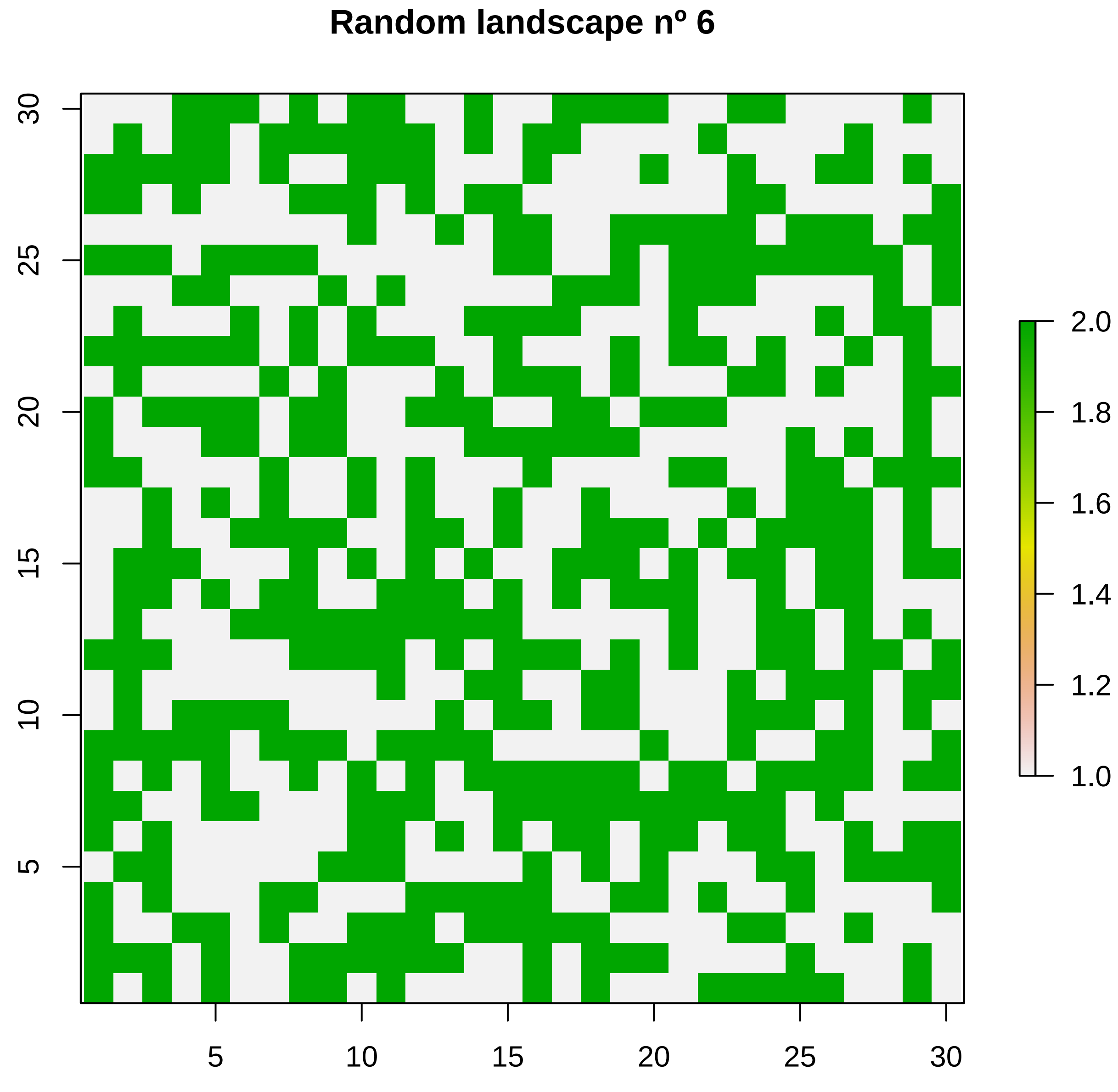
Random landscape nº 6
5
10
15
20
25
30
5
10
15
20
25
30
2.0
1.8
1.6
1.4
1.2
1.0

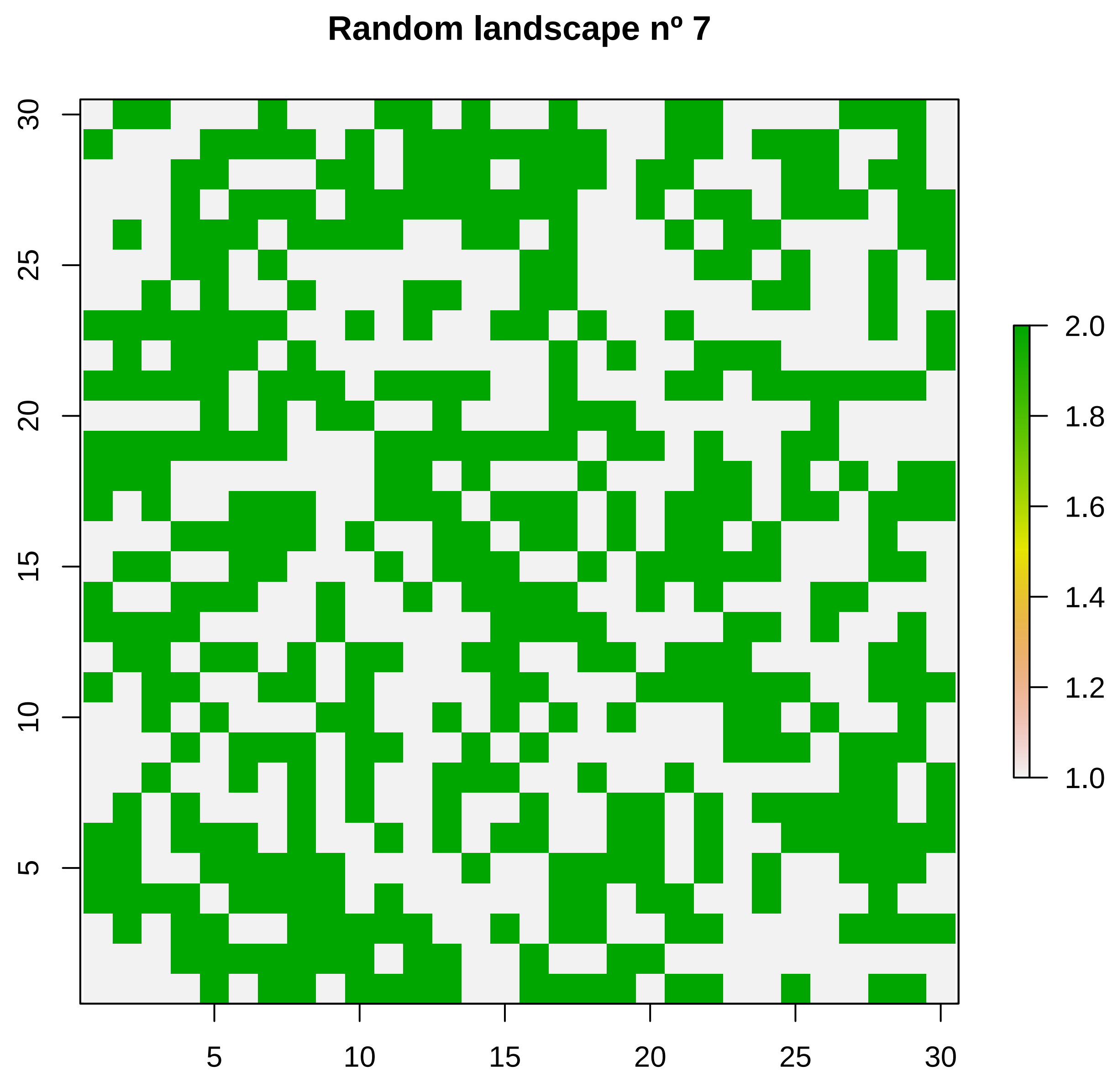

Random landscape nº 7
30
25
20
15
10
5
5
10
15
20
25
30
2.0
1.8
1.6
1.4
1.2
1.0

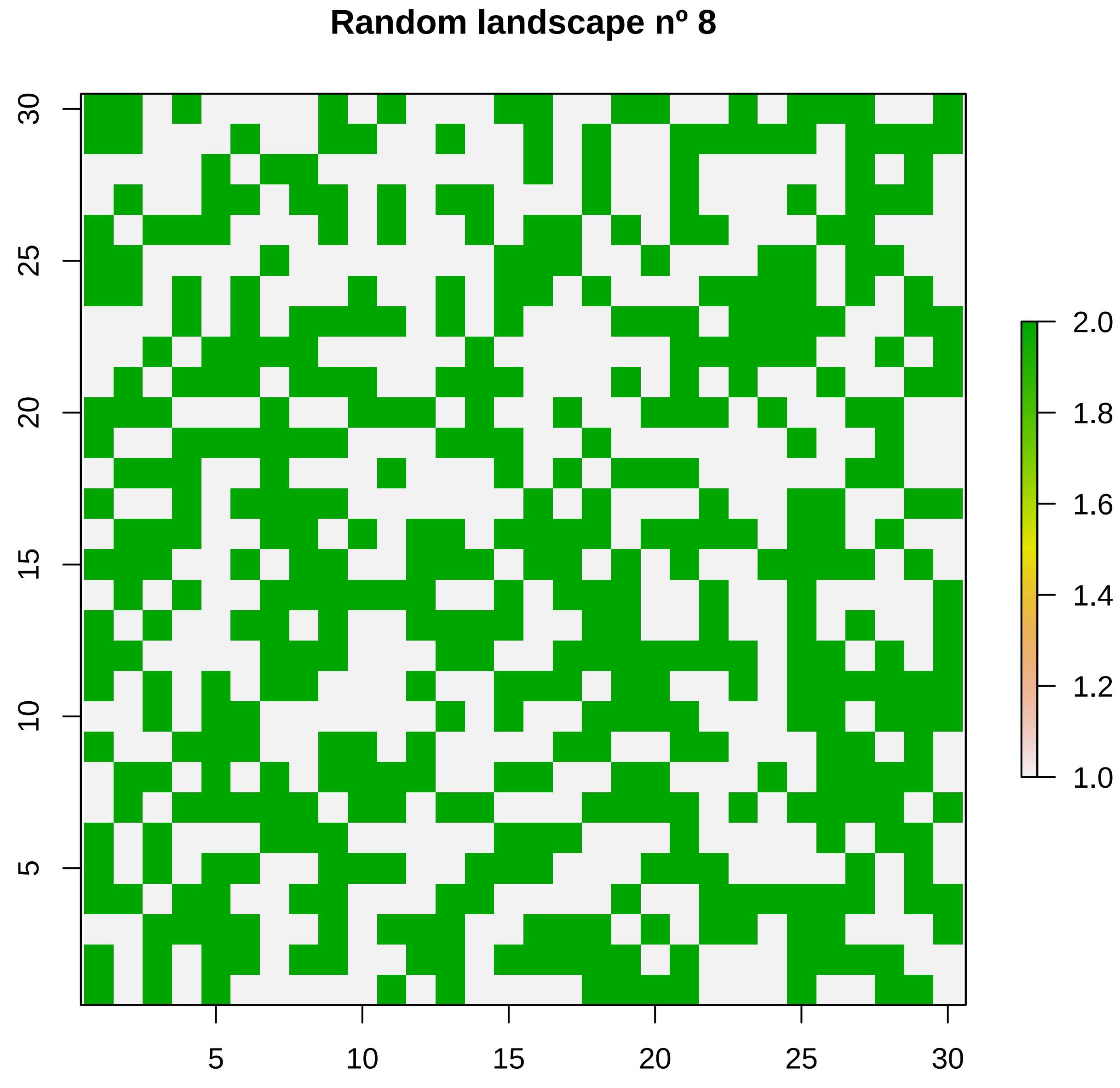
Random landscape nº 8
5
10
15
20
25
30
1.0
1.2
1.4
1.6
1.8
2.0

#### Random landscape nº 9

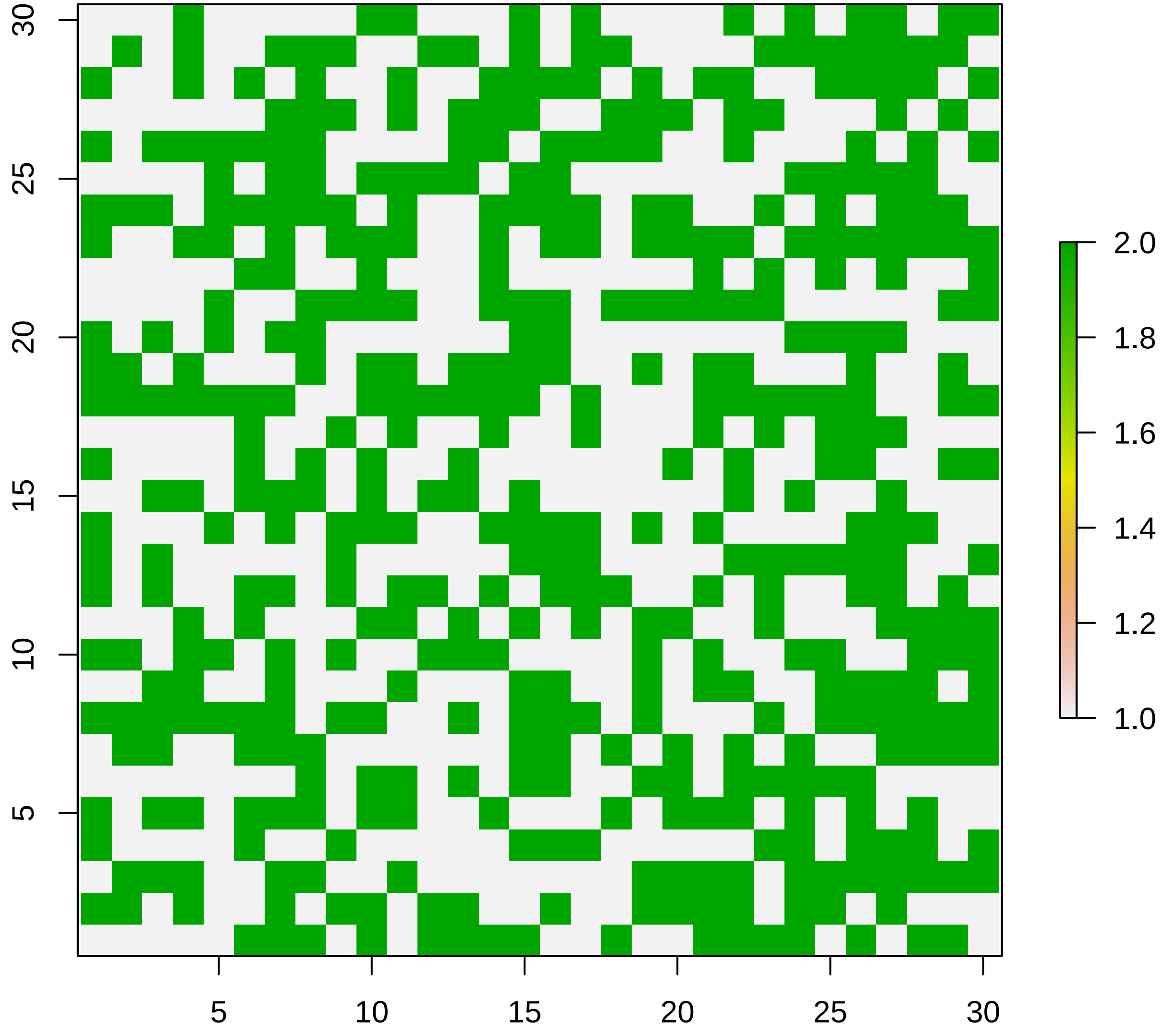

**Random landscape nº 10**

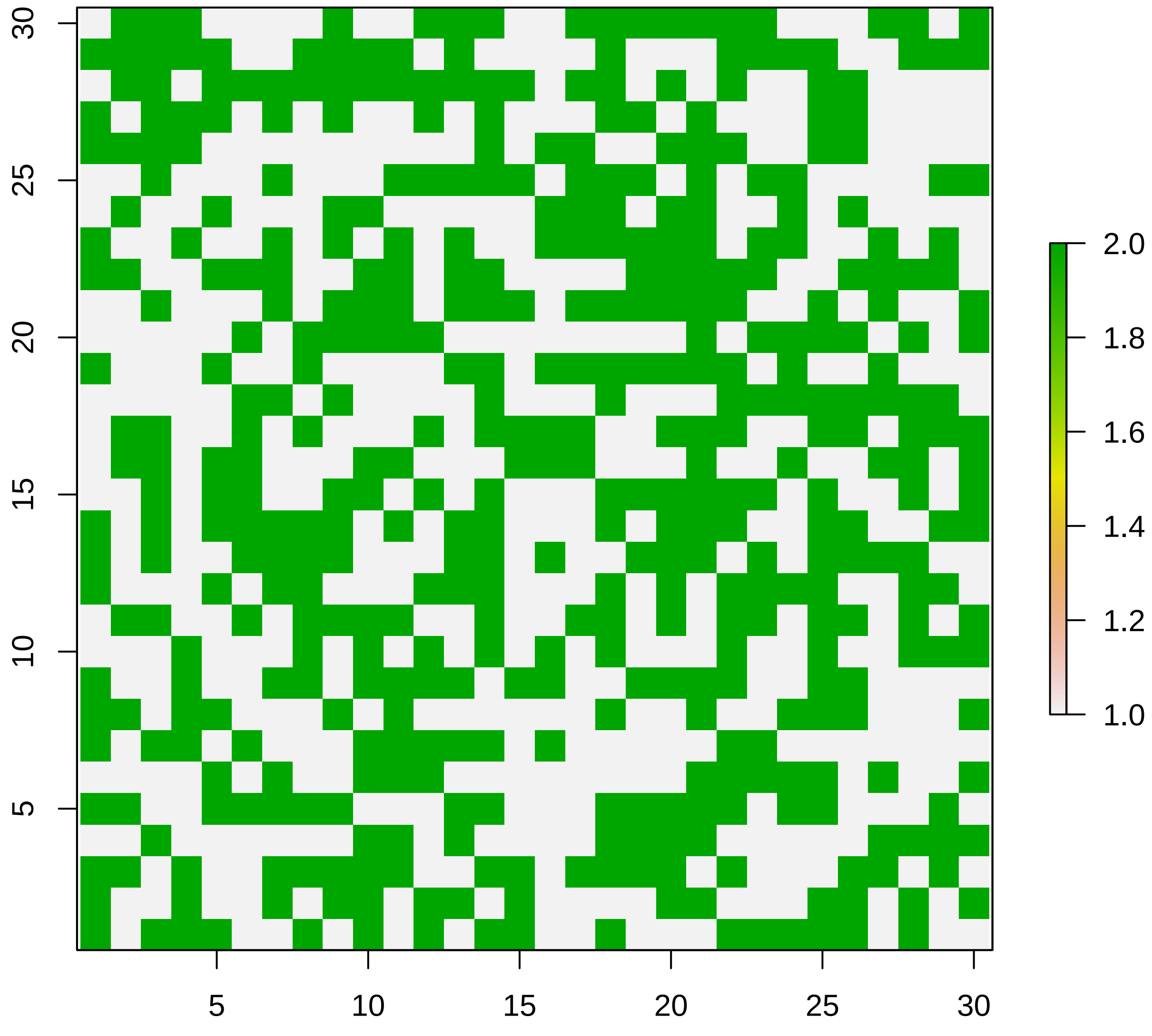

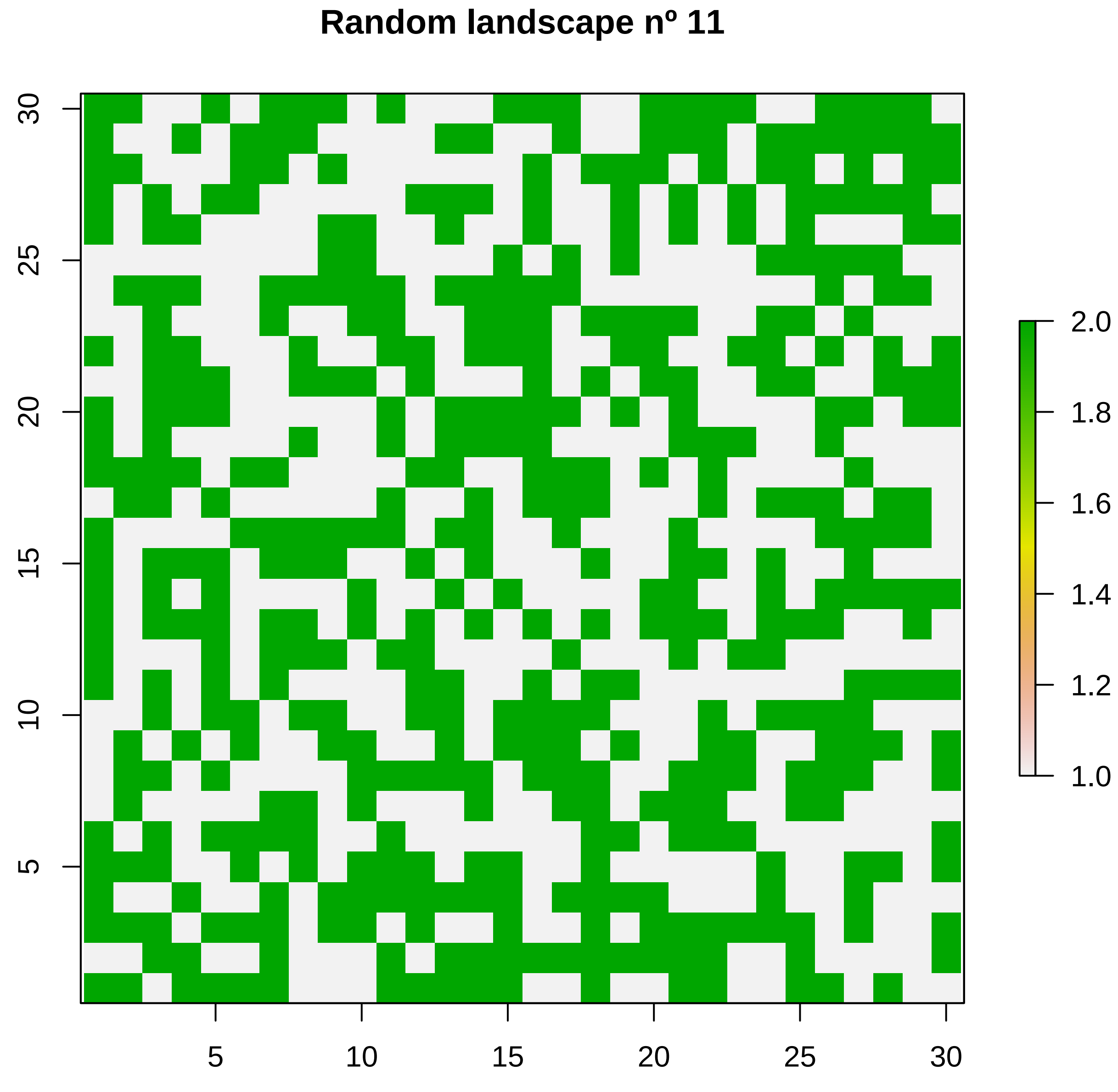

Random landscape nº 11
30
25
20
15
10
5
5
10
15
20
25
30
2.0
1.8
1.6
1.4
1.2
1.0

### Random landscape nº 12

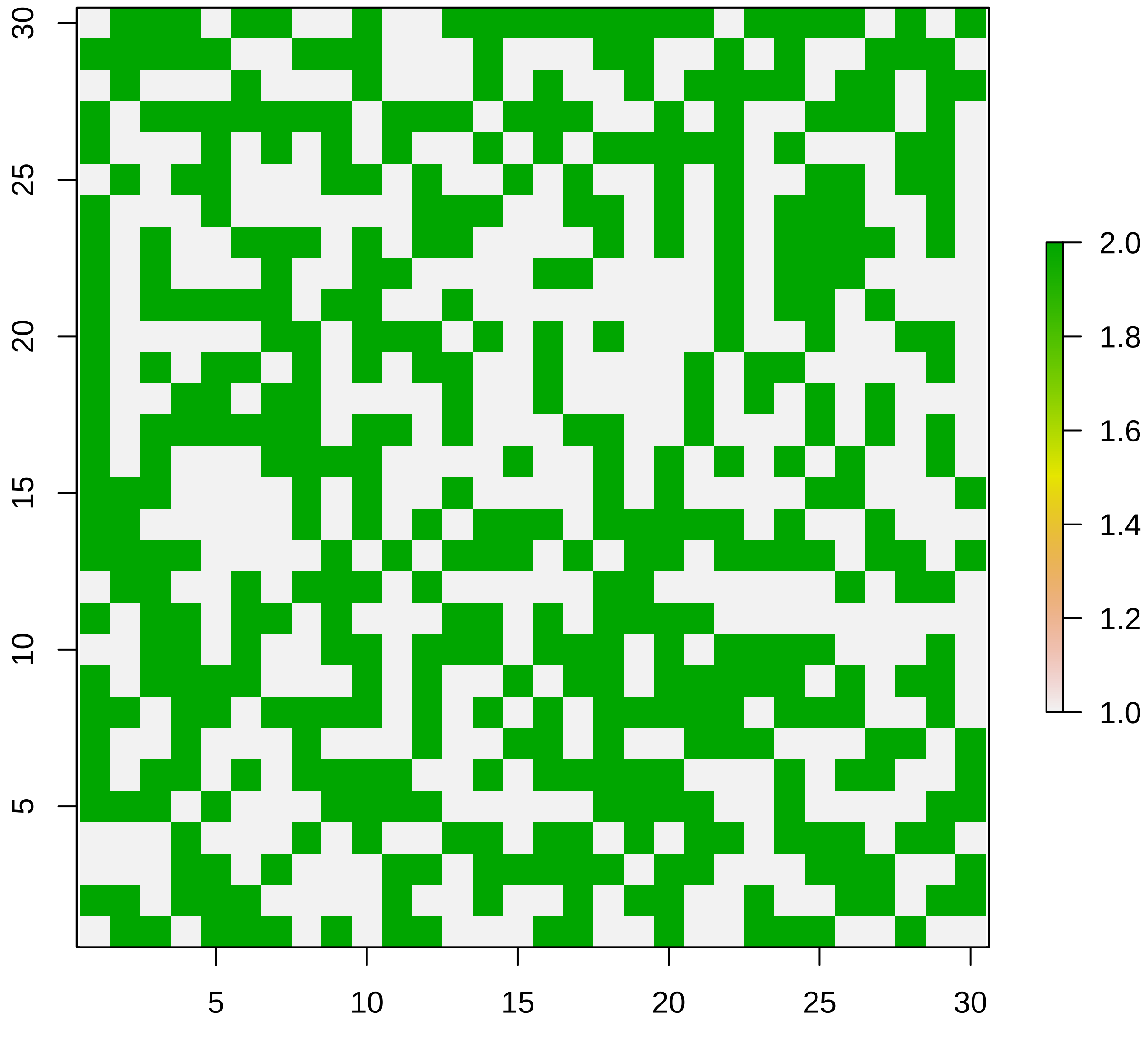

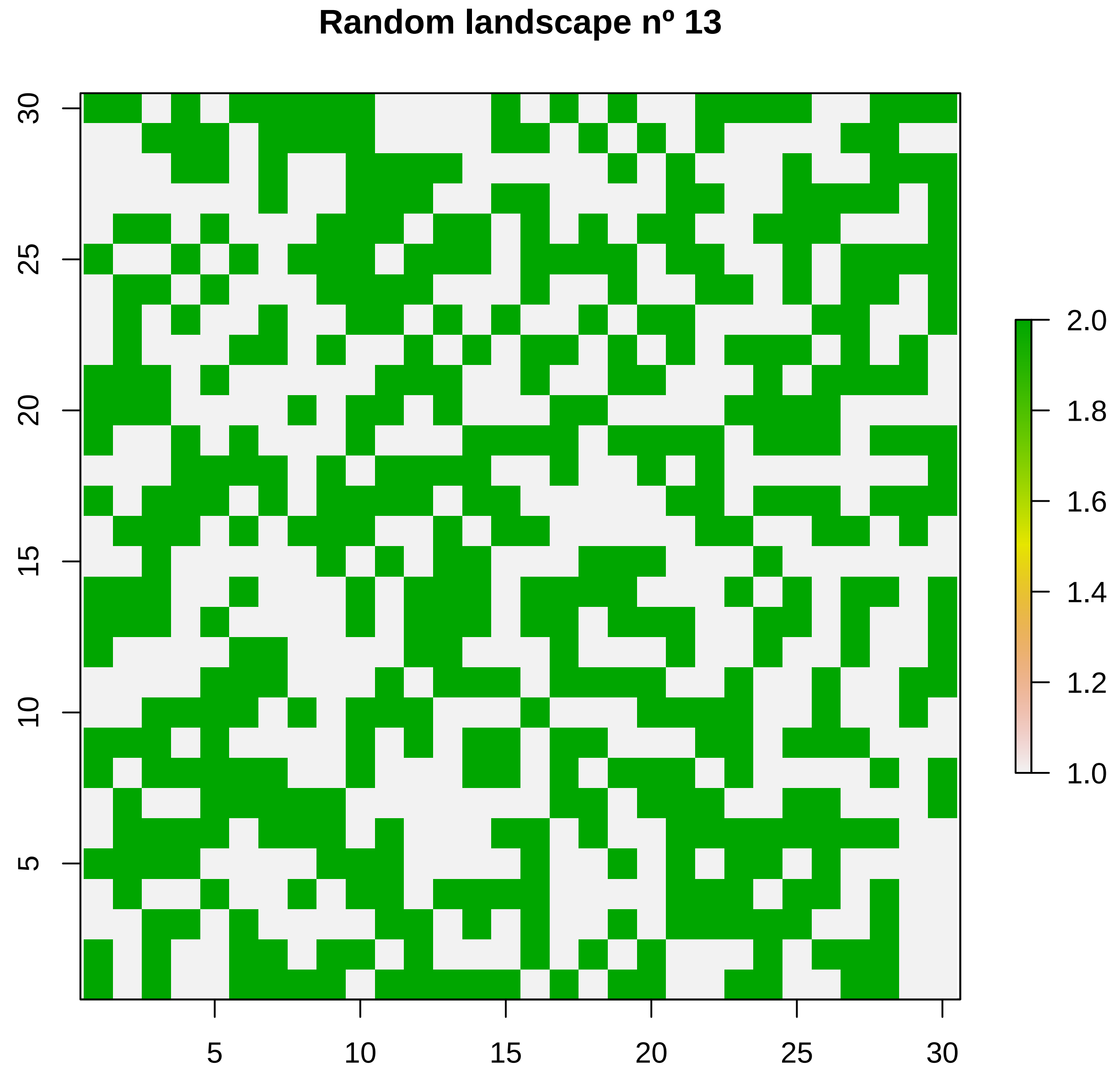
Random landscape nº 13
5
10
15
20
25
30
5
10
15
20
25
30
1.0
1.2
1.4
1.6
1.8
2.0

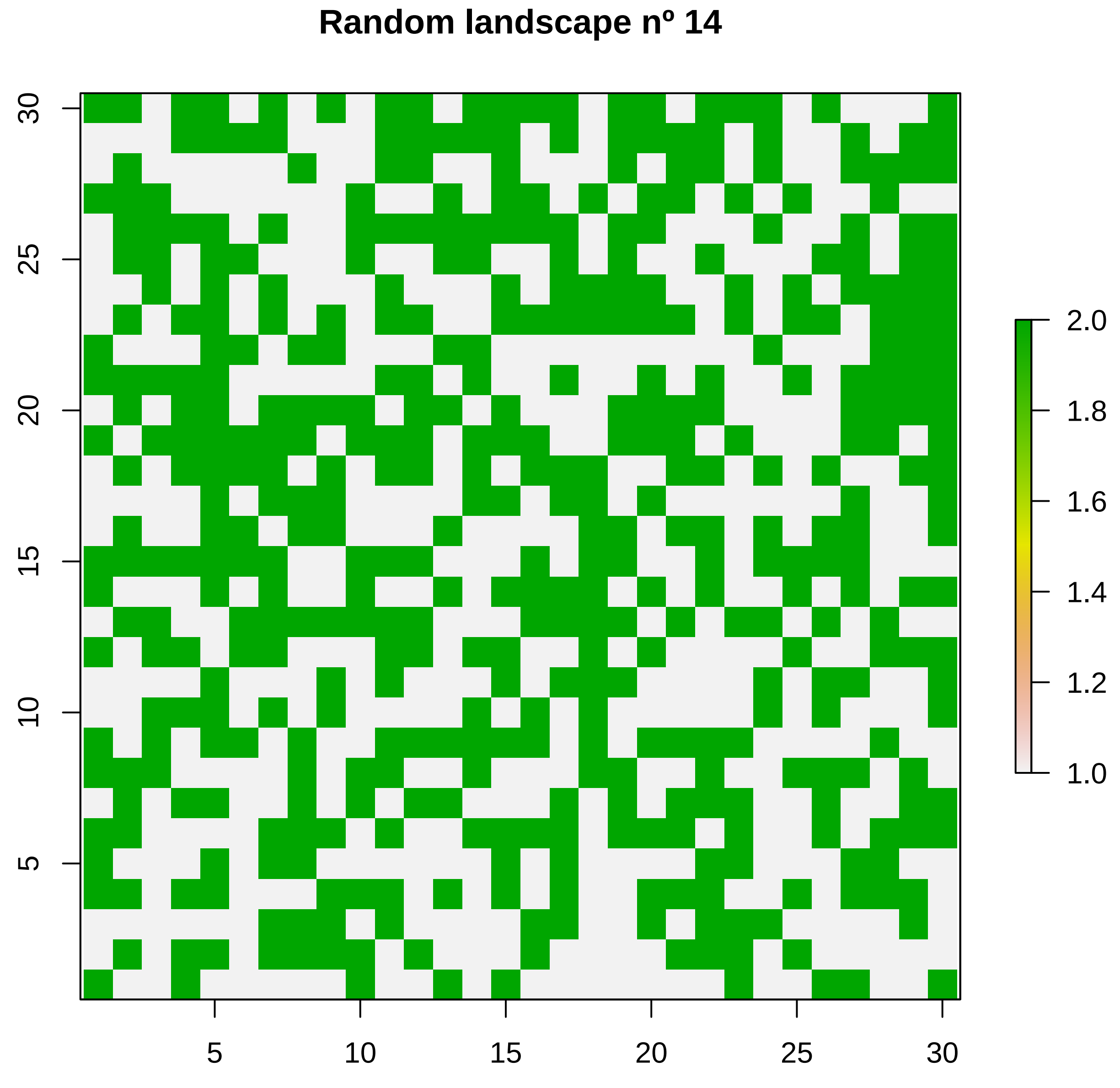
Random landscape nº 14
5
10
15
20
25
30
5
10
15
20
25
30
1.0
1.2
1.4
1.6
1.8
2.0

#### Random landscape nº 15

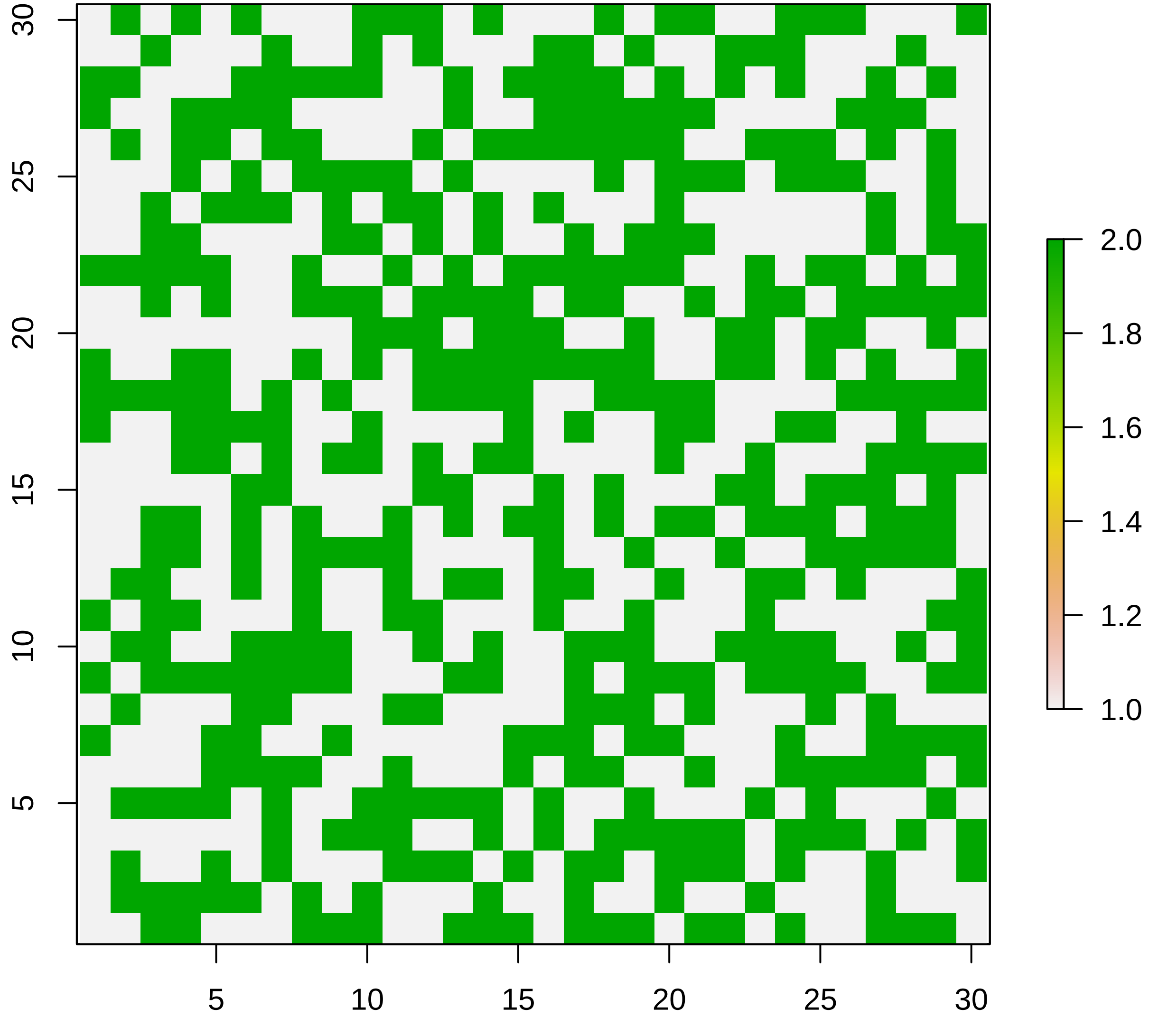

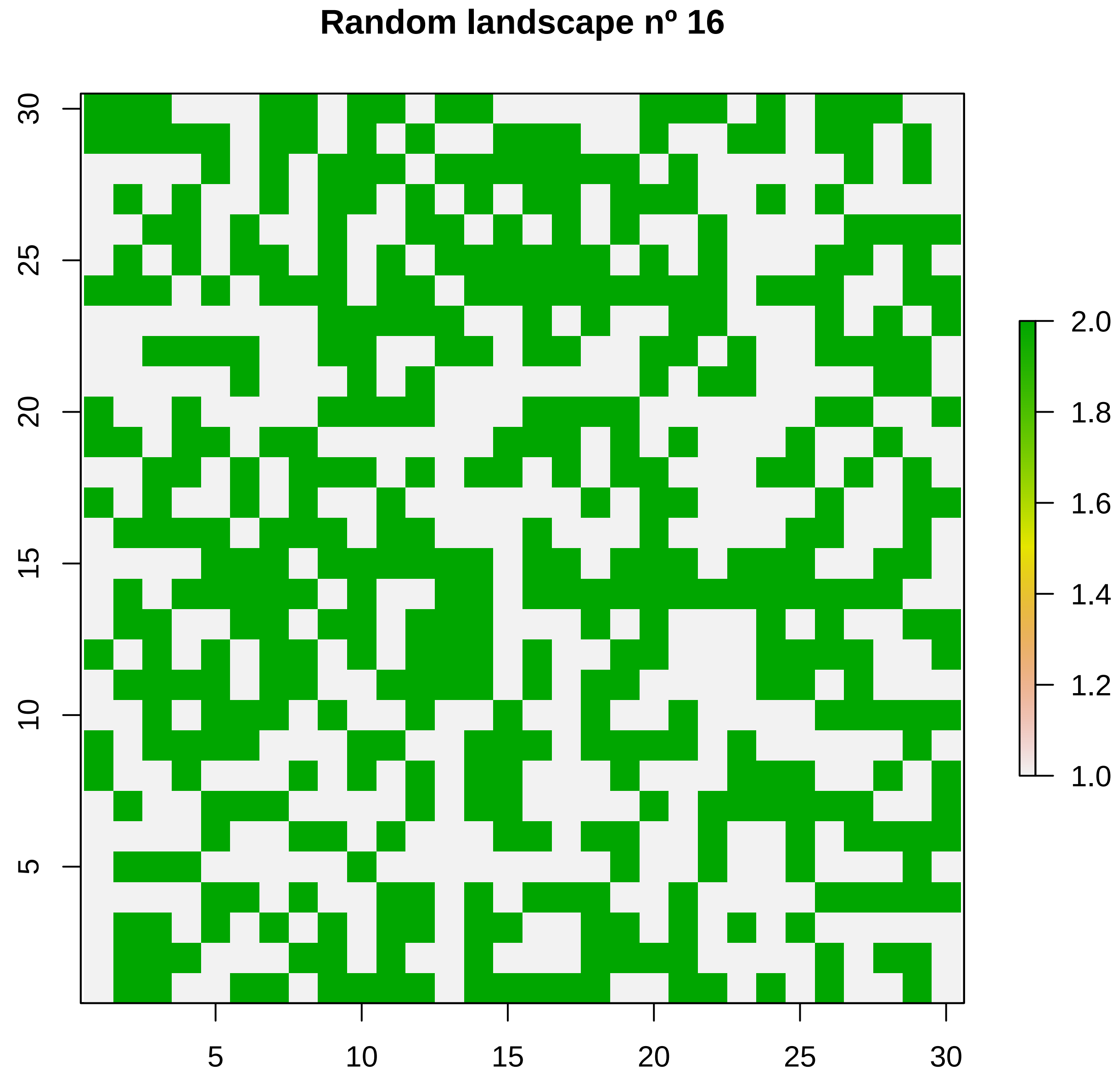

Random landscape nº 16
5
10
15
20
25
30
5
10
15
20
25
30
2.0
1.8
1.6
1.4
1.2
1.0

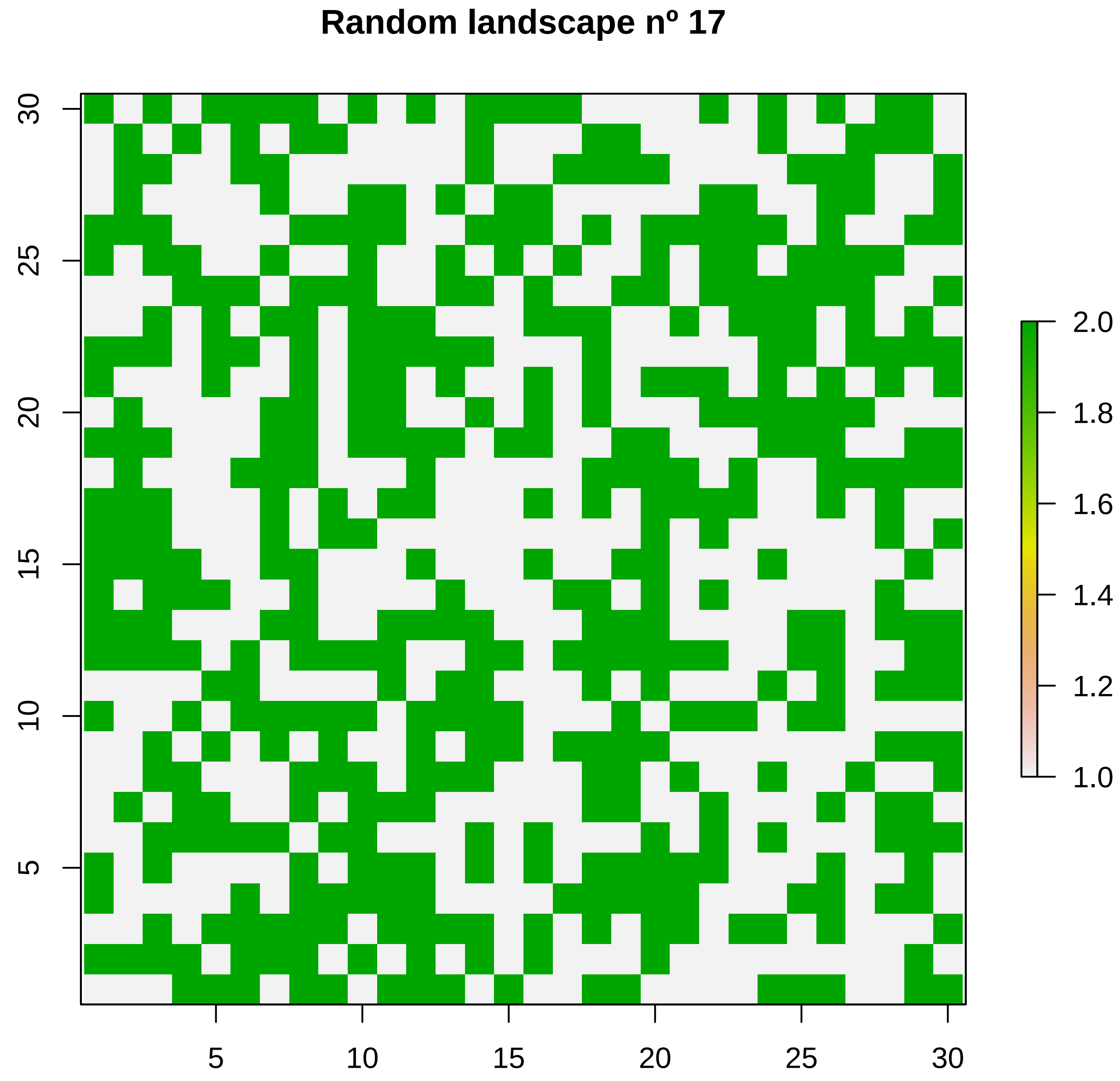

Random landscape nº 17
30
25
20
15
10
5
5
10
15
20
25
30
2.0
1.8
1.6
1.4
1.2
1.0

#### Random landscape nº 18

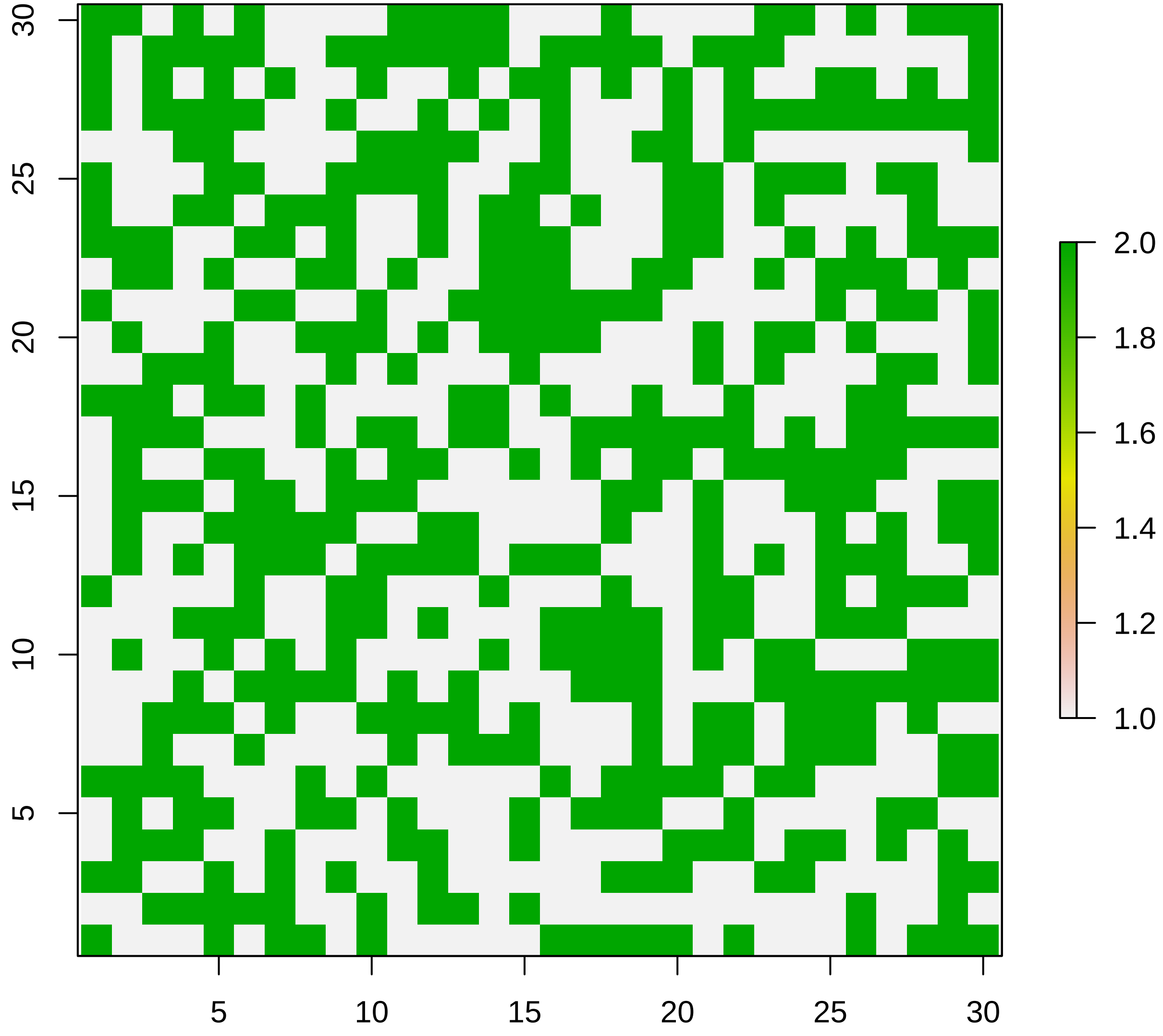

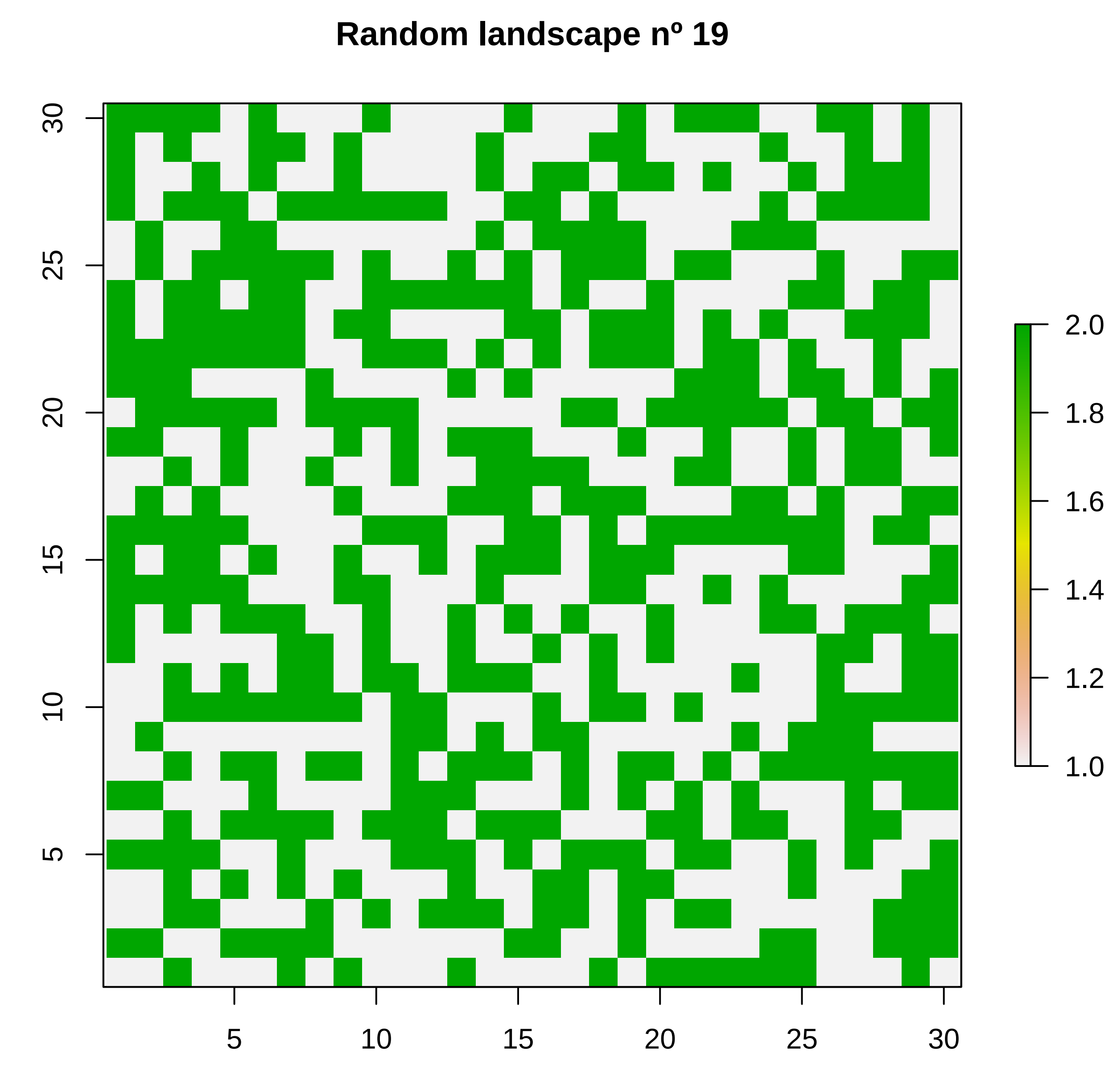

Random landscape nº 19
30
25
20
15
10
5
5
10
15
20
25
30
2.0
1.8
1.6
1.4
1.2
1.0

**Random landscape nº 20**

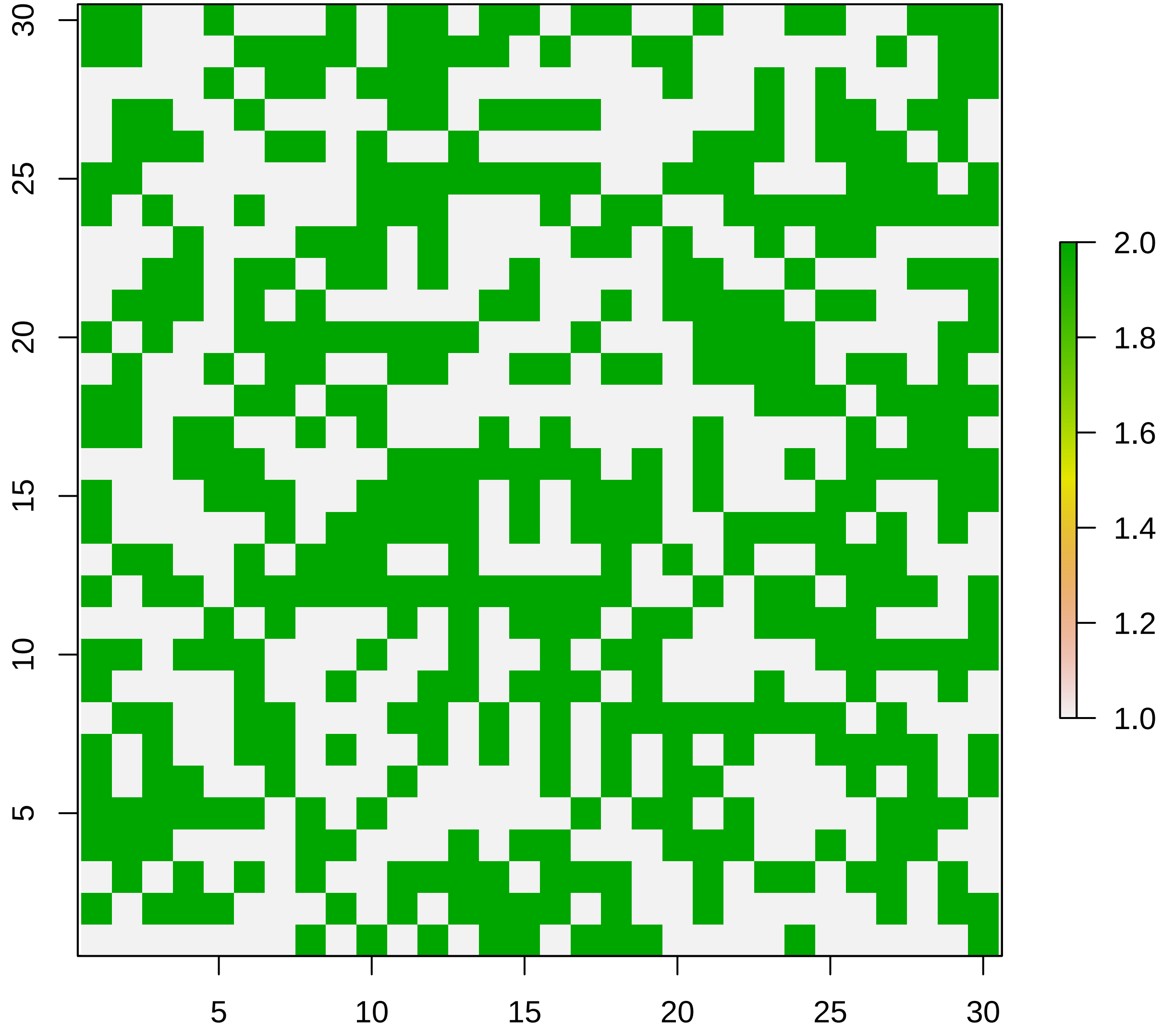
