## Supplementary material for "To disperse or compete? Coevolution of traits leads to a limited number of reproductive strategies": Dynamics for Homogeneous, Random and Aggregated landscapes with strong trade-off and high mutation rate

### Supplementary file S2:

Evolutionary dynamics of investment (E), propagule size (So), and number of propagules (E/So) over time (x-axis) for the simulations in Homogeneous, Random and Aggregated landscapes. Investment and propagule size are represented on the left y-axis, while number of propagules is on the right y-axis. Simulations were performed with mutation rate = 0.05 and strong size-dispersal trade-off.

#### Homogeneous landscapes:

A)

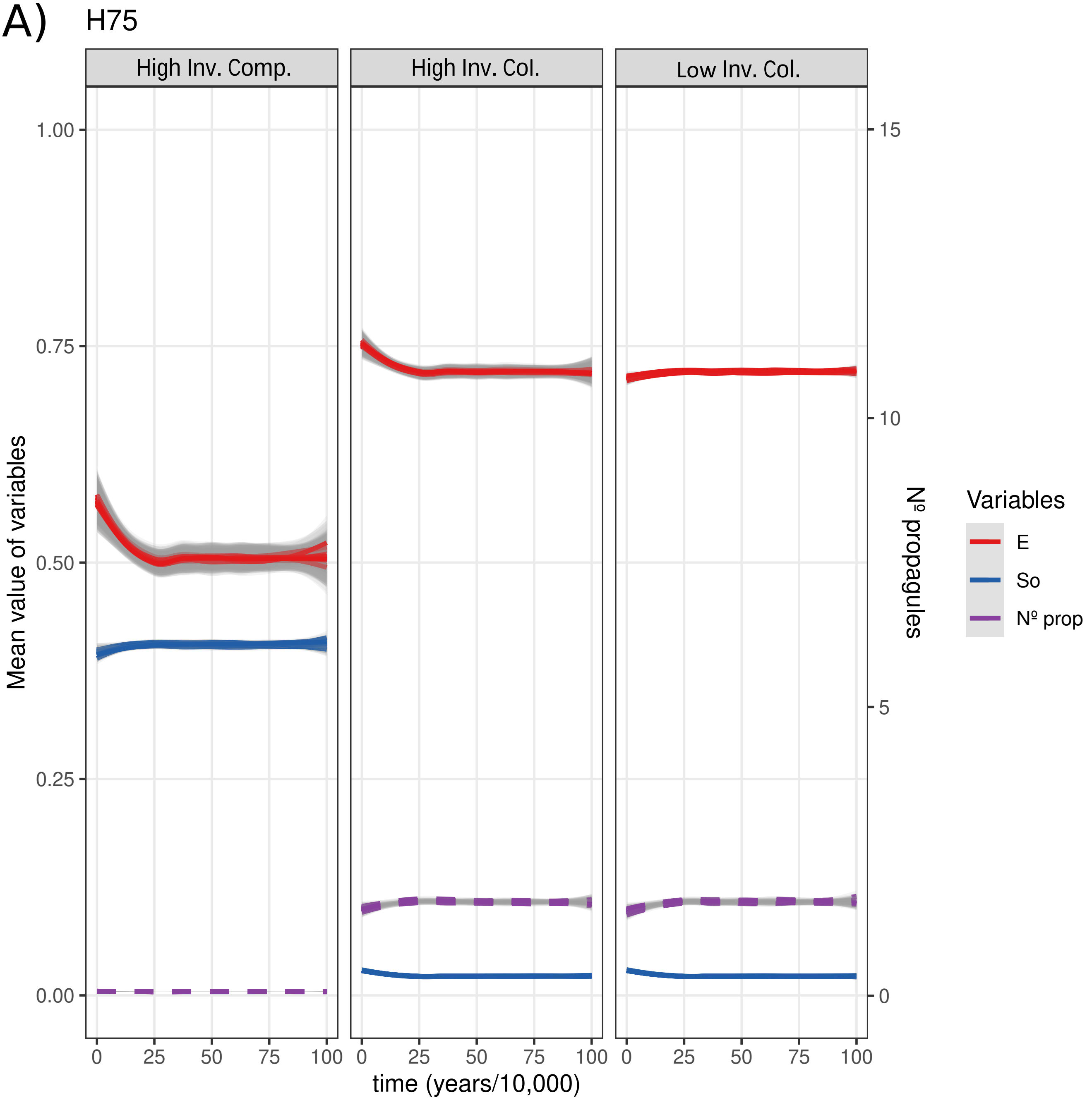


Figure A) Homogeneous landscape with K' = 75. Titles of the subplots refer to starting conditions. Each line corresponds to the mean of all values (~900) obtained from a given simulation, and thus we show 20 lines (20 simulations) for each starting conditions and variable (E, So and Nº propagules), with the corresponding 95% CI (in grey).

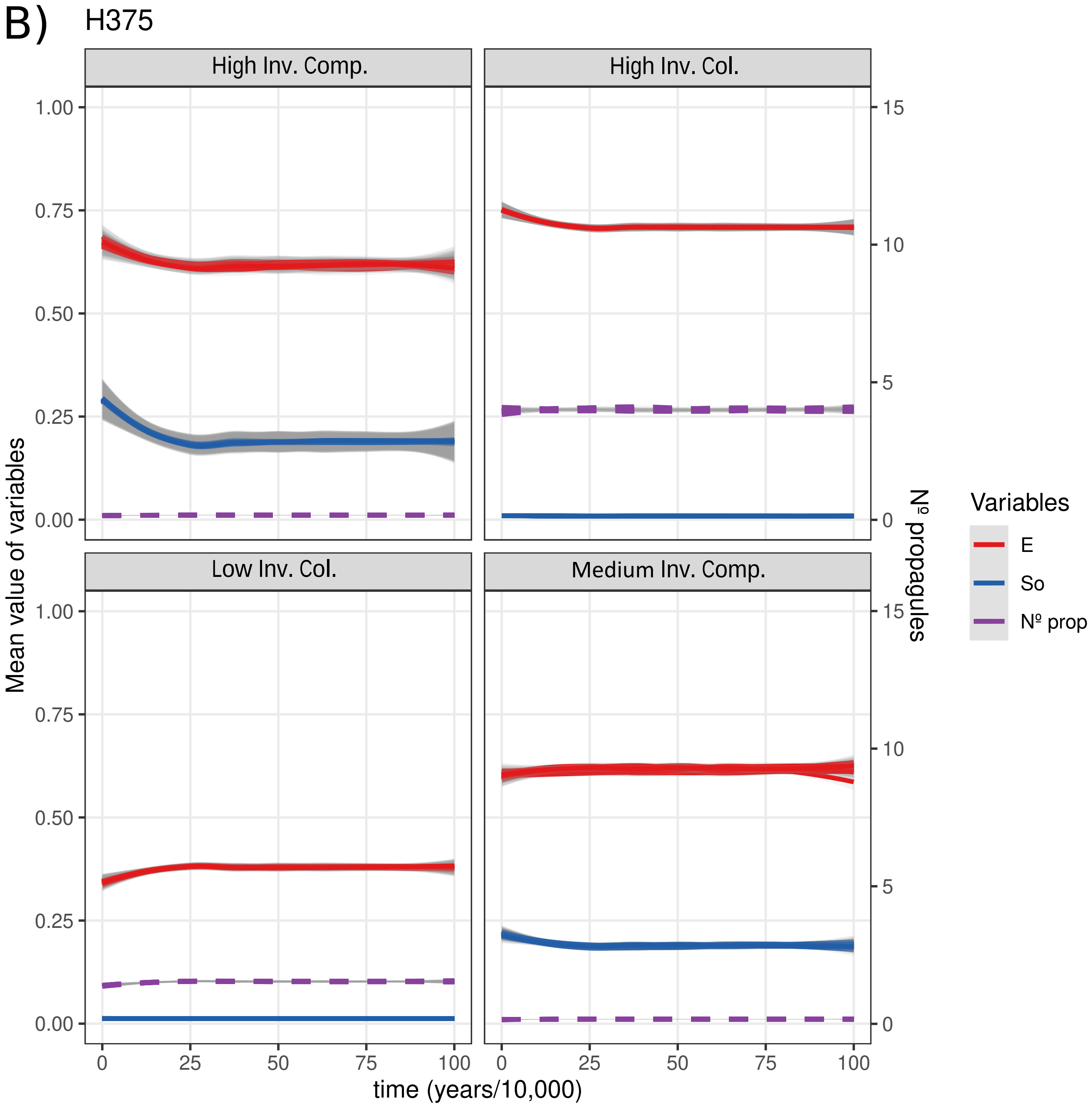


Figure B) Homogeneous landscape with K’ = 375. Titles of the subplots refer to starting conditions. Each line corresponds to the mean of all values (~900) obtained from a given simulation, and thus we show 20 lines (20 simulations) for each starting conditions and variable (E, SO and Nº propagules), with the corresponding 95% CI (in grey).

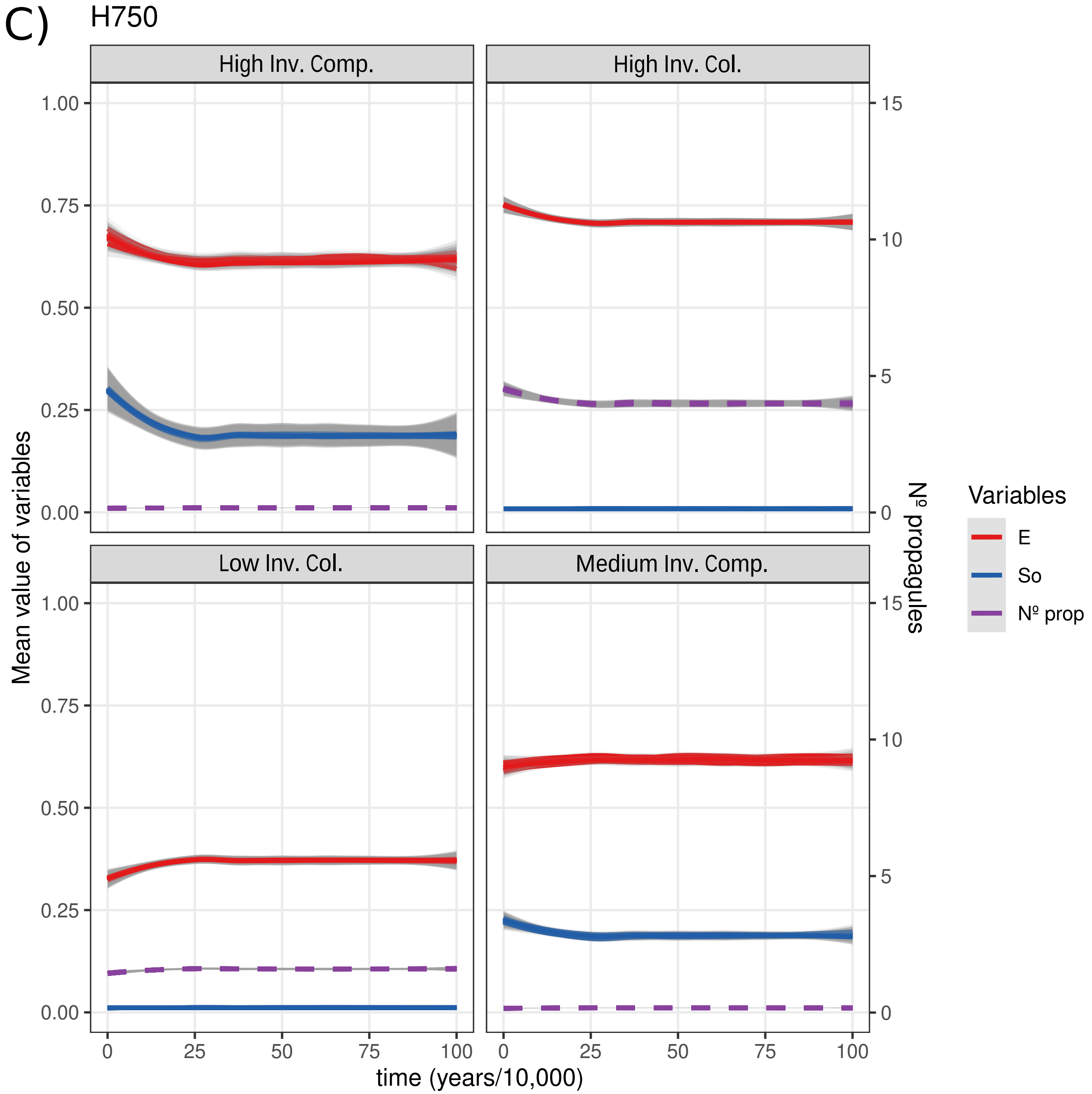


Figure C) Homogeneous landscape with K’ = 750. Titles of the subplots refer to starting conditions. Each line corresponds to the mean of all values (~900) obtained from a given simulation, and thus we show 20 lines (20 simulations) for each starting conditions and variable (E, SO and Nº propagules), with the corresponding 95% CI (in grey).

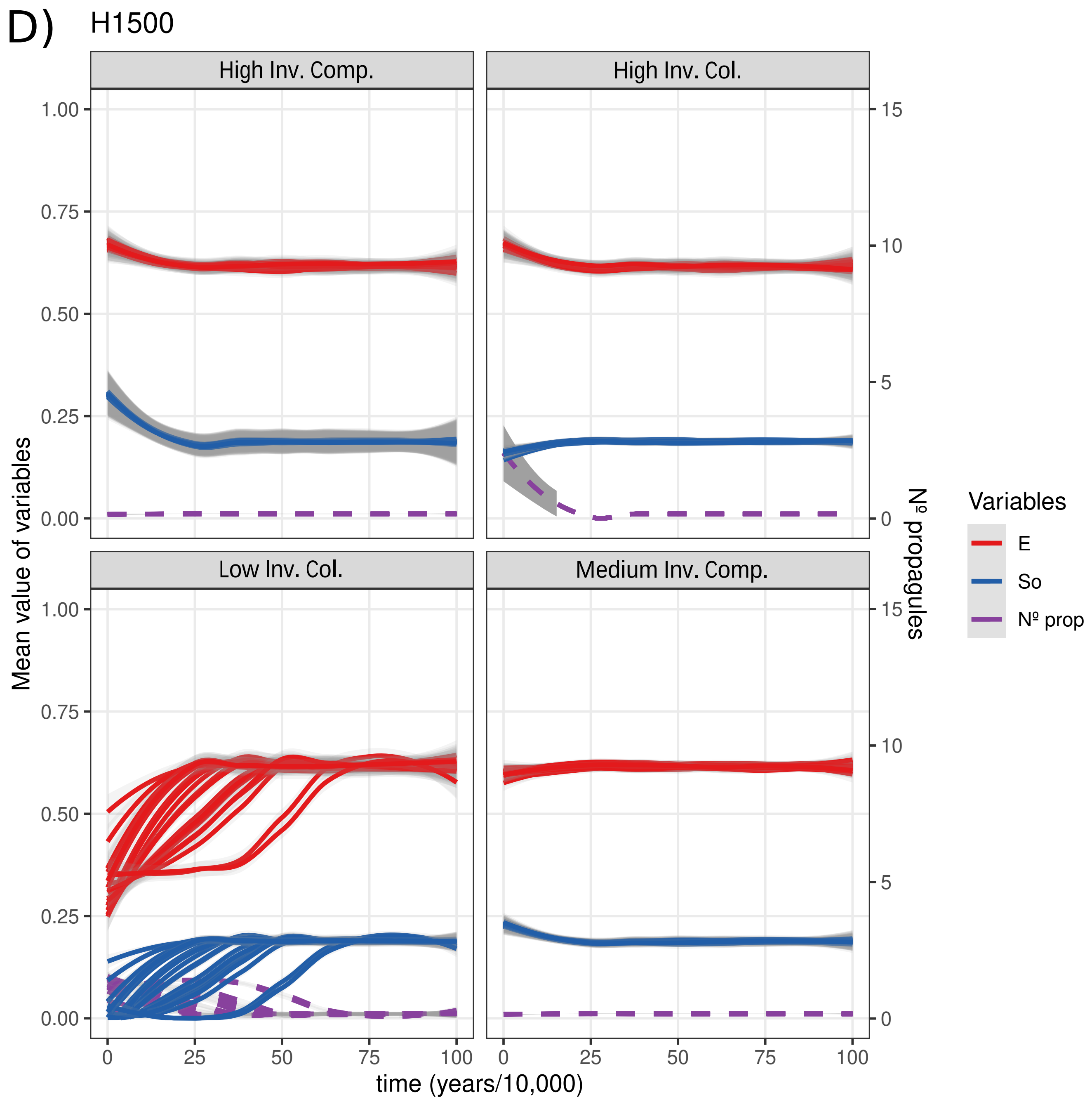


Figure D) Homogeneous landscape with K’ = 1500. Titles of the subplots refer to starting conditions. Each line corresponds to the mean of all values (~900) obtained from a given simulation, and thus we show 20 lines (20 simulations) for each starting conditions and variable (E, SO and Nº propagules), with the corresponding 95% CI (in grey).

### Random landscapes:

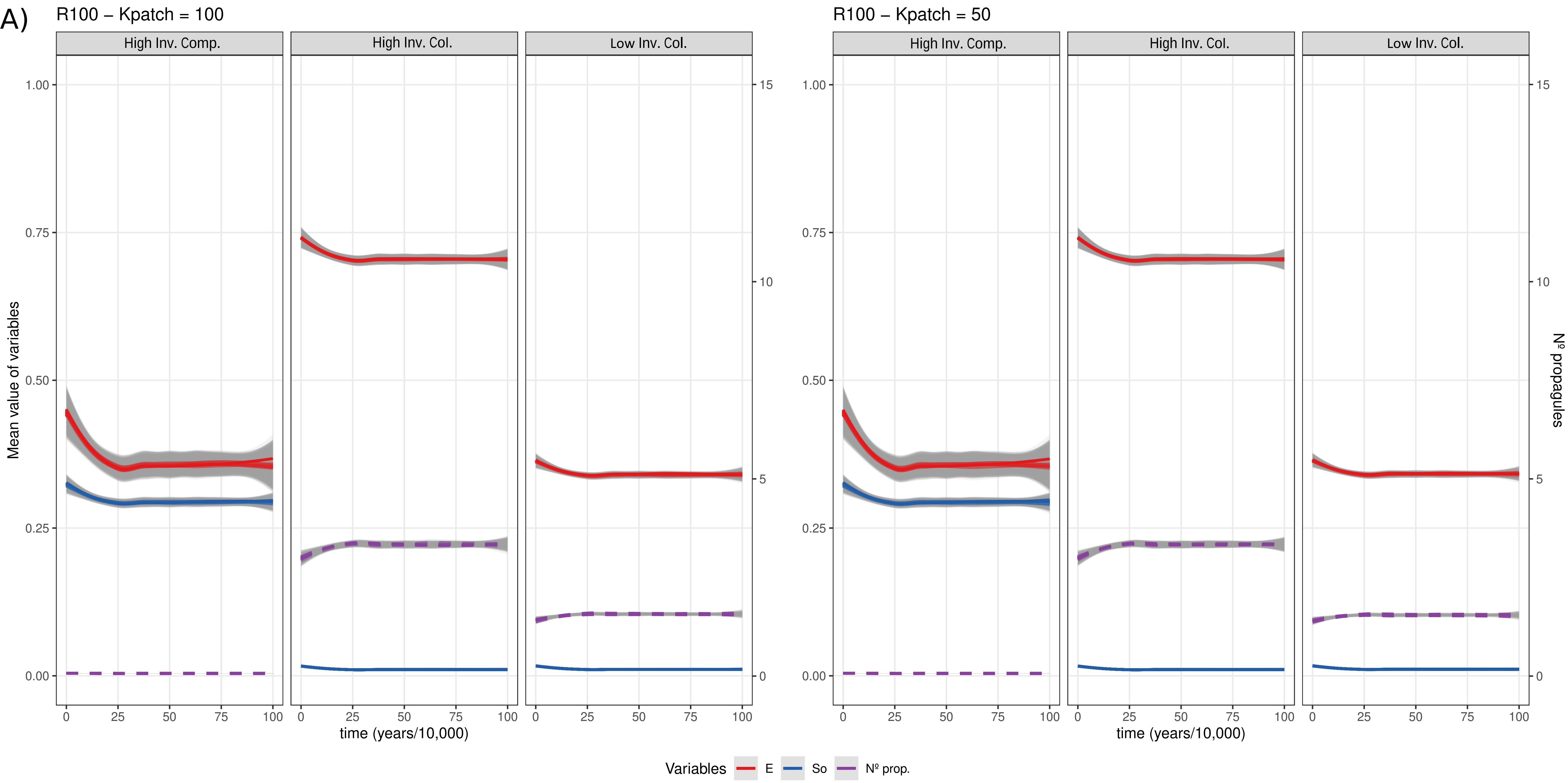


Figure A) Random landscape with K = 100. Results have been divided depending on rich or poor patches. This is indicated on the top as Kpatch, with rich patches (on the left) having a value = K, and poor patches (on the right) a value of K/2. Titles of the subplots refer to starting conditions. Each line corresponds to the mean of all values (~450) obtained from a given simulation, and thus we show 20 lines (20 simulations) for each starting conditions and variable (E, So and Nº propagules), with the corresponding 95% CI (in grey).

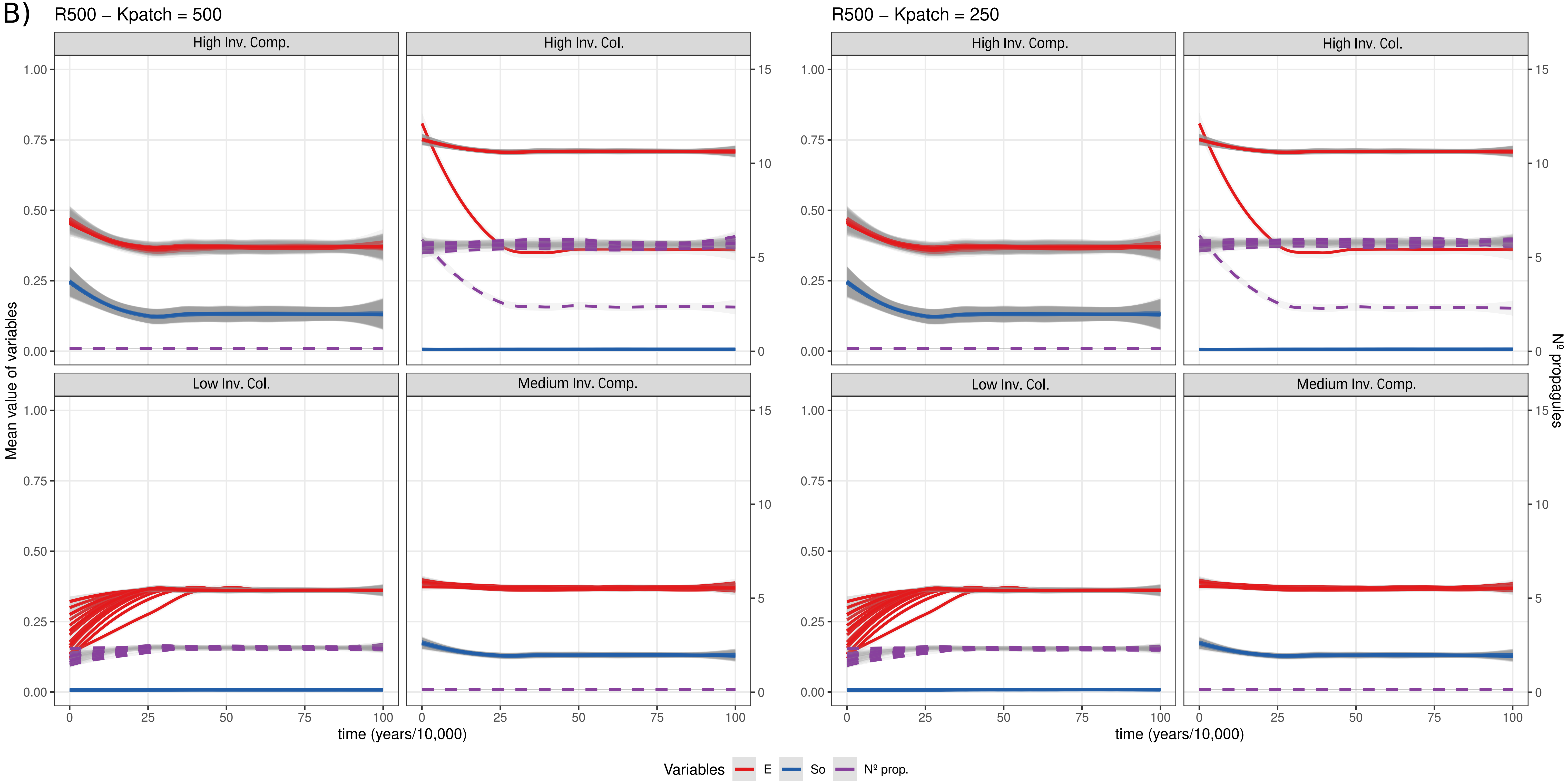


Figure B) Random landscape with K = 500. Results have been divided depending on rich or poor patches. This is indicated on the top as Kpatch, with rich patches (on the left) having a value = K, and poor patches (on the right) a value of K/2. Titles of the subplots refer to starting conditions. Each line corresponds to the mean of all values (~450) obtained from a given simulation, and thus we show 20 lines (20 simulations) for each starting conditions and variable (E, So and Nº propagules), with the corresponding 95% CI (in grey).

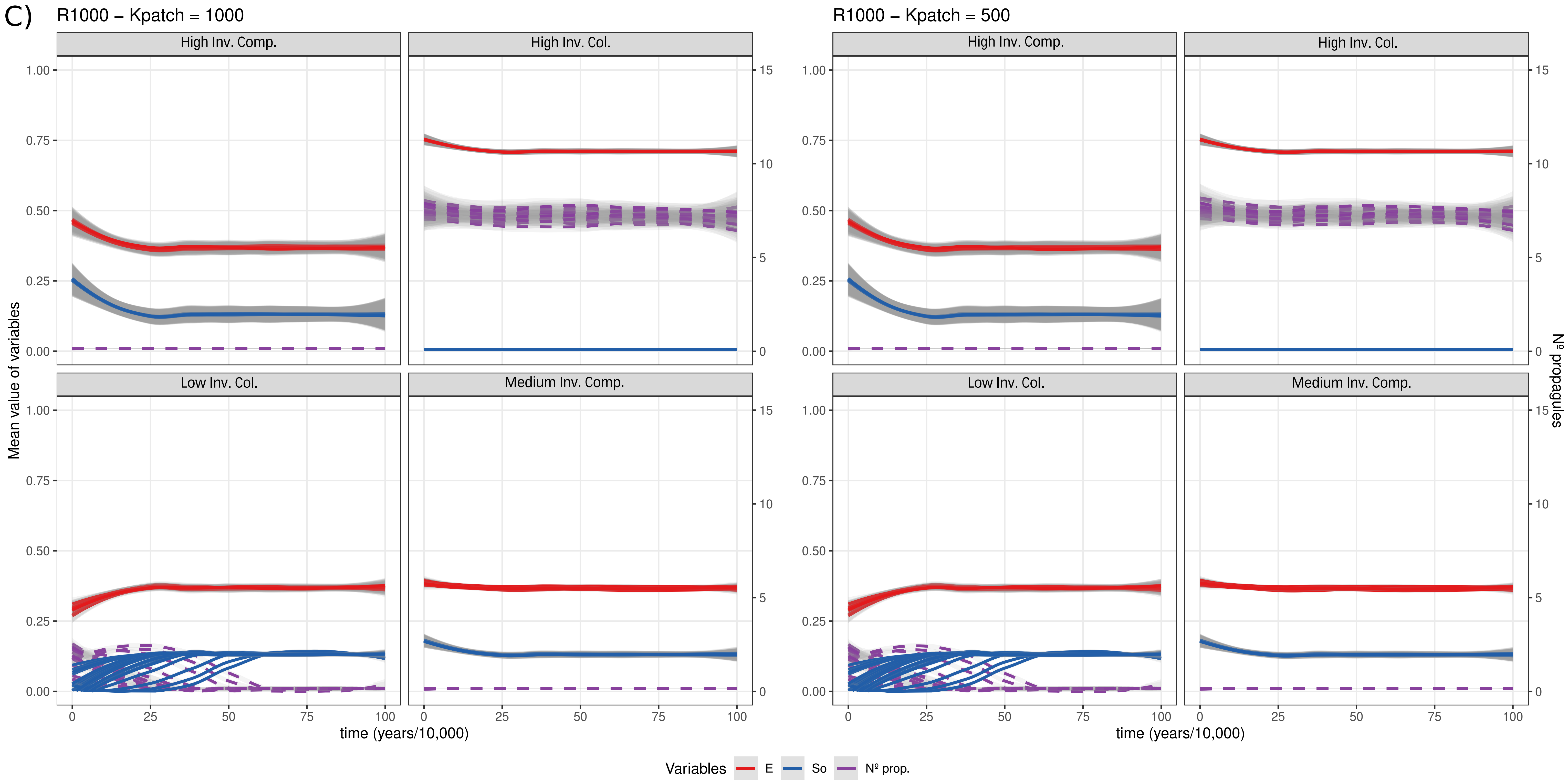


Figure C) Random landscape with K = 1000. Results have been divided depending on rich or poor patches. This is indicated on the top as Kpatch, with rich patches (on the left) having a value = K, and poor patches (on the right) a value of K/2. Titles of the subplots refer to starting conditions. Each line corresponds to the mean of all values (~450) obtained from a given simulation, and thus we show 20 lines (20 simulations) for each starting conditions and variable (E, So and Nº propagules), with the corresponding 95% CI (in grey).

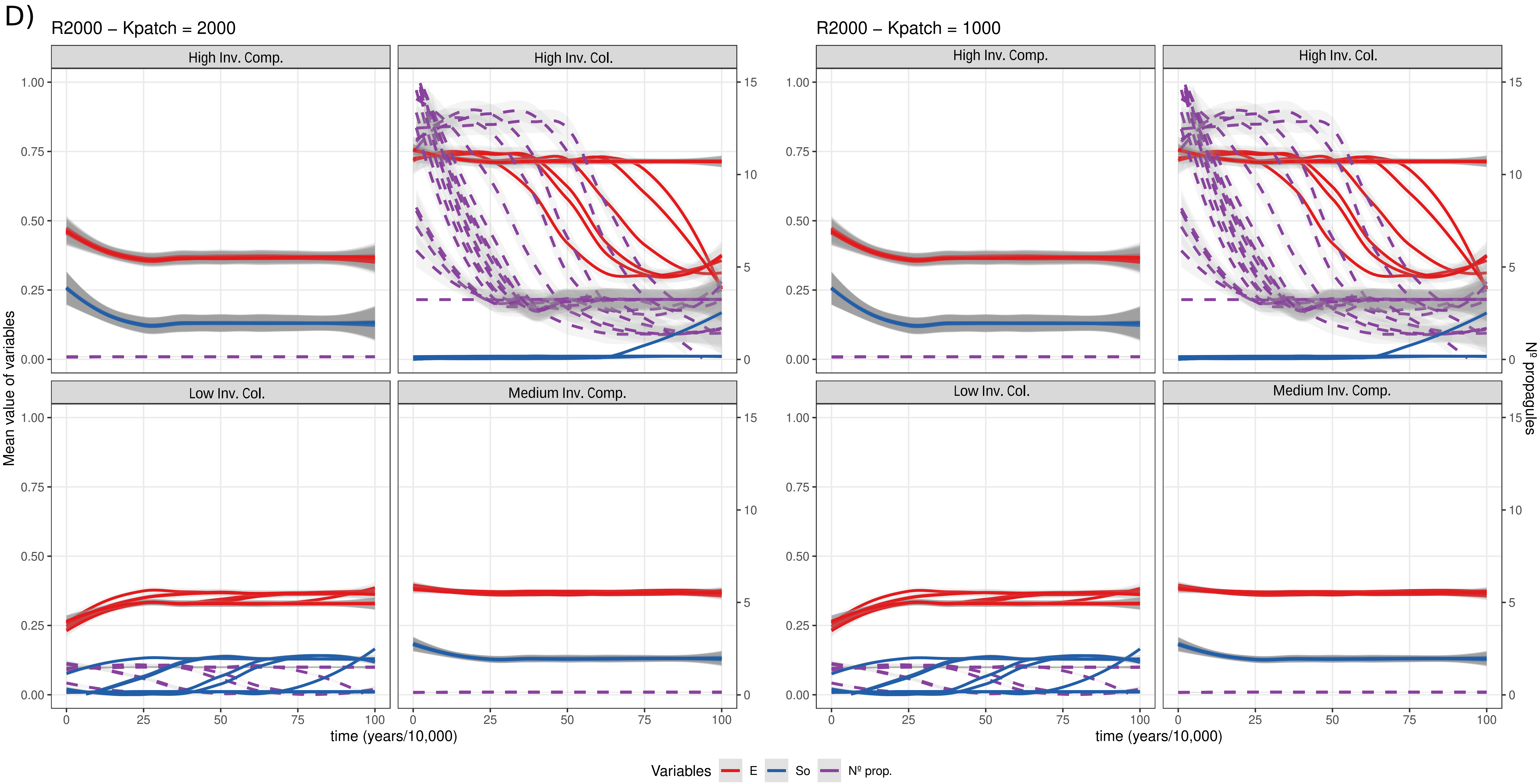


Figure D) Random landscape with K = 2000. Results have been divided depending on rich or poor patches. This is indicated on the top as Kpatch, with rich patches (on the left) having a value = K, and poor patches (on the right) a value of K/2. Titles of the subplots refer to starting conditions. Each line corresponds to the mean of all values (~450) obtained from a given simulation, and thus we show 20 lines (20 simulations) for each starting conditions and variable (E, So and Nº propagules), with the corresponding 95% CI (in grey).

### Aggregated landscapes:

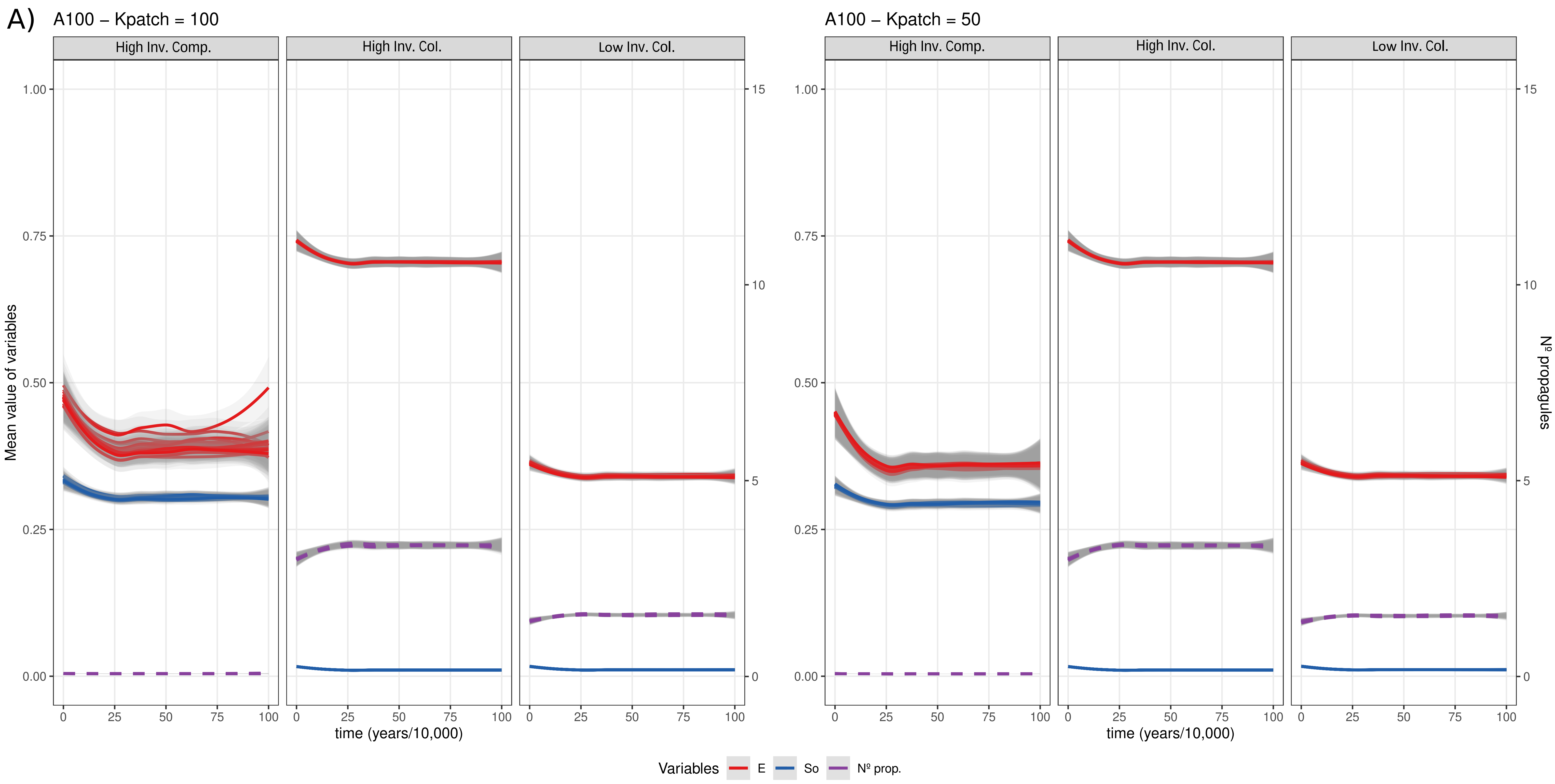


Figure A) Aggregated landscape with K = 100. Results have been divided depending on rich or poor patches. This is indicated on the top as Kpatch, with rich patches (on the left) having a value = K, and poor patches (on the right) a value of K/2. Titles of the subplots refer to starting conditions. Each line corresponds to the mean of all values (~450) obtained from a given simulation, and thus we show 20 lines (20 simulations) for each starting conditions and variable (E, So and Nº propagules), with the corresponding 95% CI (in grey).

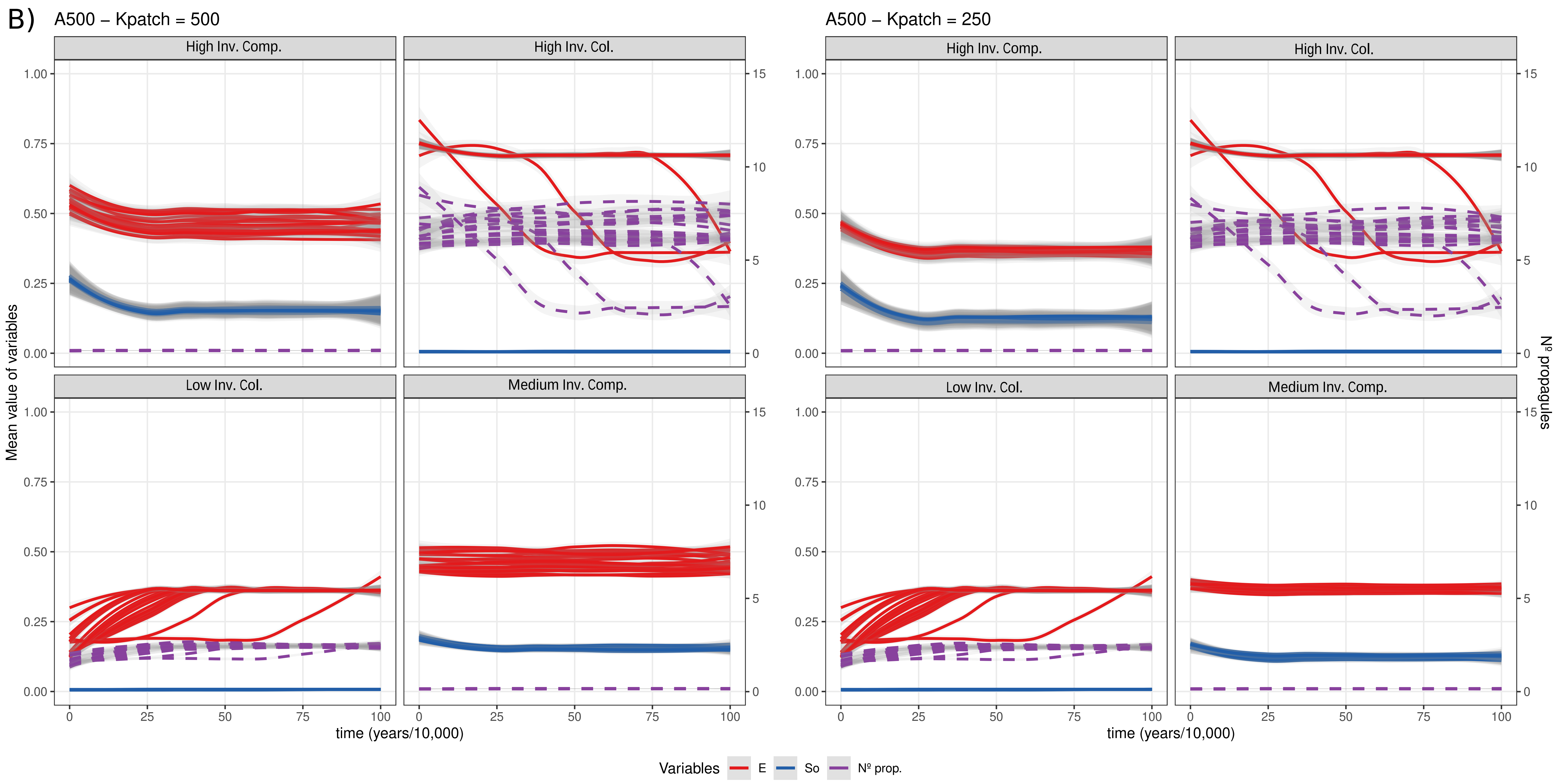


Figure B) Aggregated landscape with K = 500. Results have been divided depending on rich or poor patches. This is indicated on the top as Kpatch, with rich patches (on the left) having a value = K, and poor patches (on the right) a value of K/2. Titles of the subplots refer to starting conditions. Each line corresponds to the mean of all values (~450) obtained from a given simulation, and thus we show 20 lines (20 simulations) for each starting conditions and variable (E, So and Nº propagules), with the corresponding 95% CI (in grey).

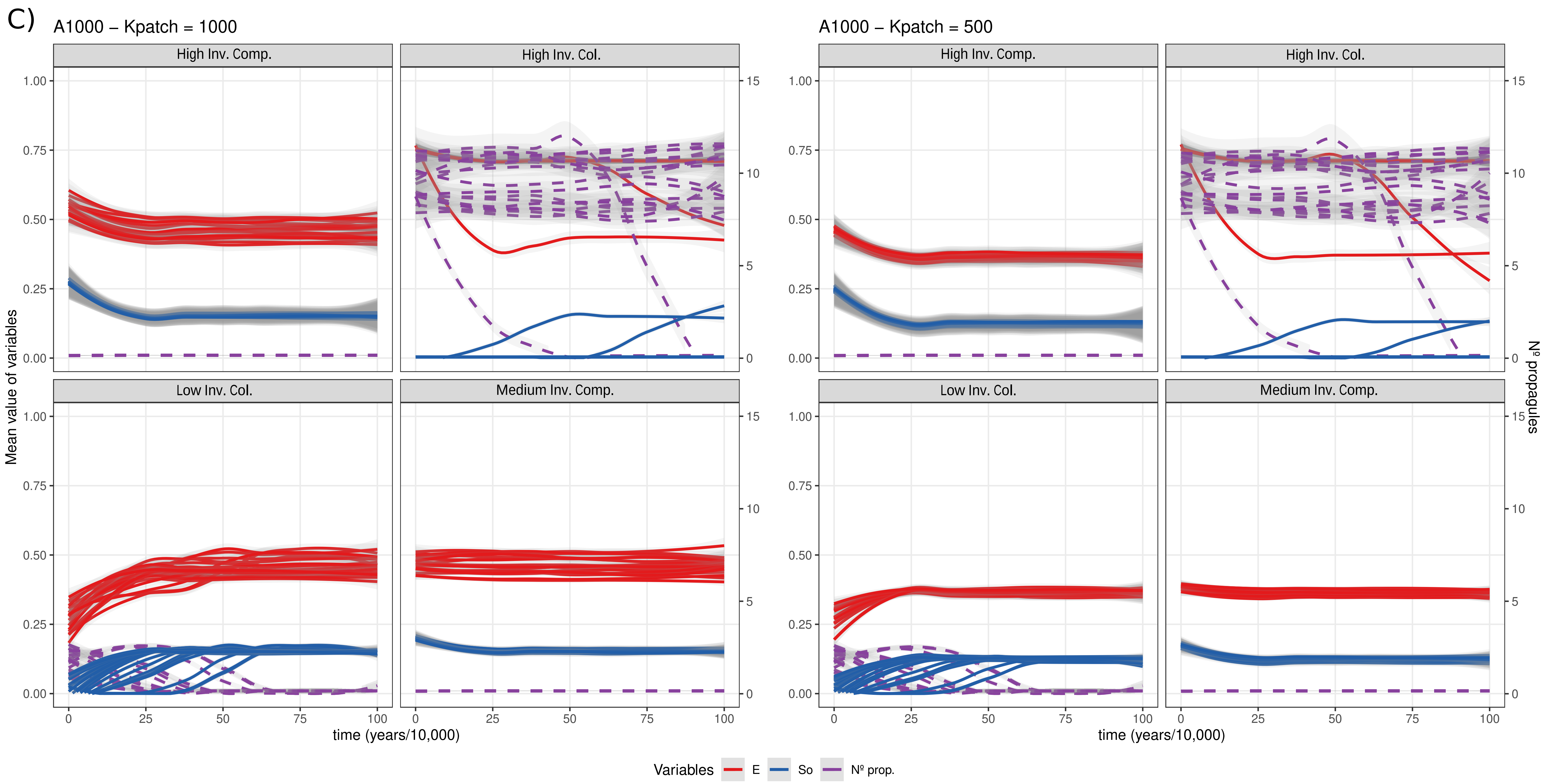


Figure C) Aggregated landscape with K = 1000. Results have been divided depending on rich or poor patches. This is indicated on the top as Kpatch, with rich patches (on the left) having a value = K, and poor patches (on the right) a value of K/2. Titles of the subplots refer to starting conditions. Each line corresponds to the mean of all values (~450) obtained from a given simulation, and thus we show 20 lines (20 simulations) for each starting conditions and variable (E, So and Nº propagules), with the corresponding 95% CI (in grey).

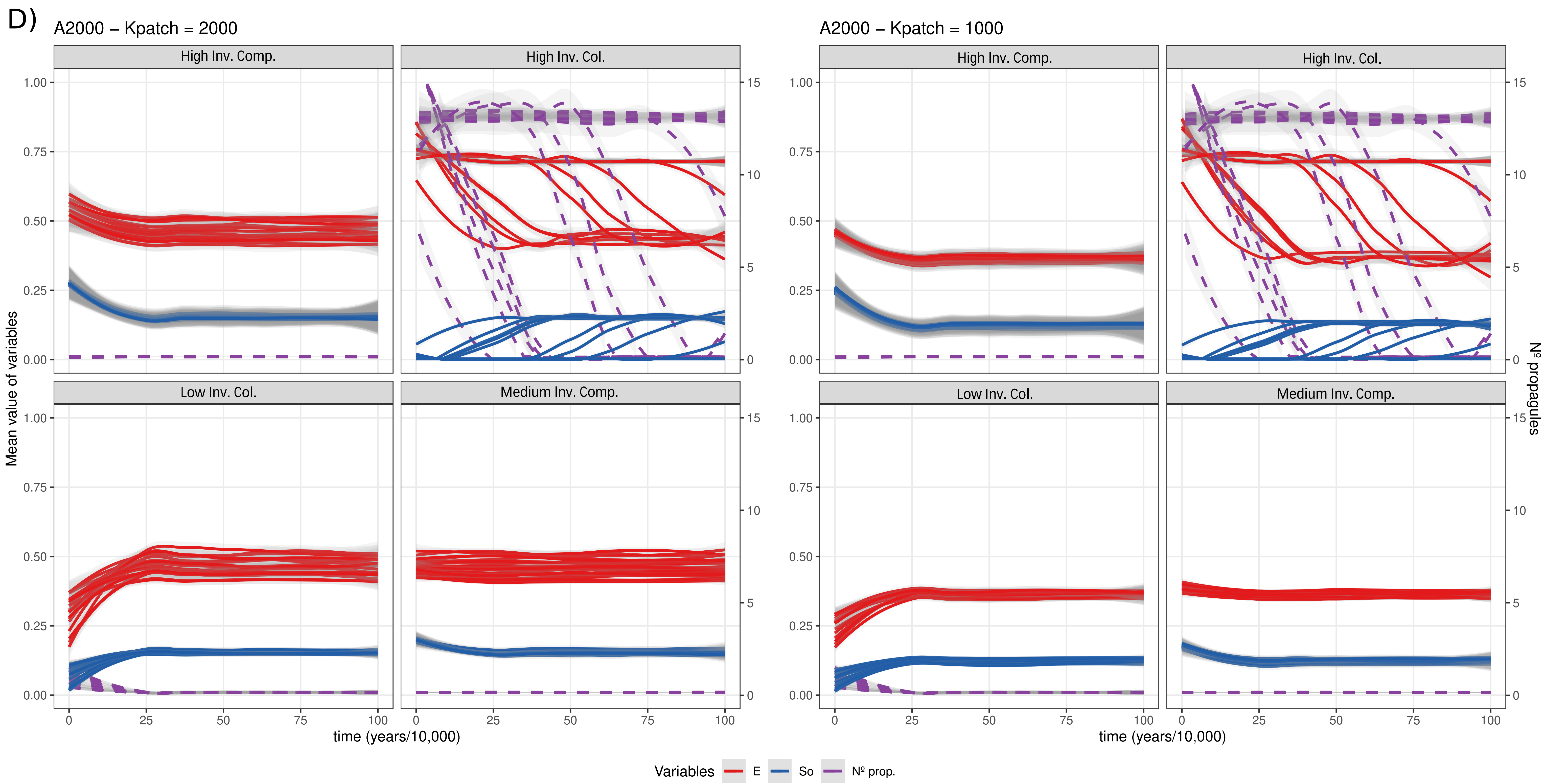


Figure D) Aggregated landscape with K = 2000. Results have been divided depending on rich or poor patches. This is indicated on the top as Kpatch, with rich patches (on the left) having a value = K, and poor patches (on the right) a value of K/2. Titles of the subplots refer to starting conditions. Each line corresponds to the mean of all values (~450) obtained from a given simulation, and thus we show 20 lines (20 simulations) for each starting conditions and variable (E, So and Nº propagules), with the corresponding 95% CI (in grey).
