## Supplementary material for "To disperse or compete? Coevolution of traits leads to a limited number of reproductive strategies": Dynamics for Homogeneous, Random and Aggregated landscapes with strong trade-off and low mutation rate

### Supplementary file S3:

Final equilibrium strategies in Homogeneous, Random and Aggregated landscapes for different resource levels (100, 500, 1000 and 2000), and slow (0.001) mutation rate. Simulations were performed with a strong size-dispersal trade-off.

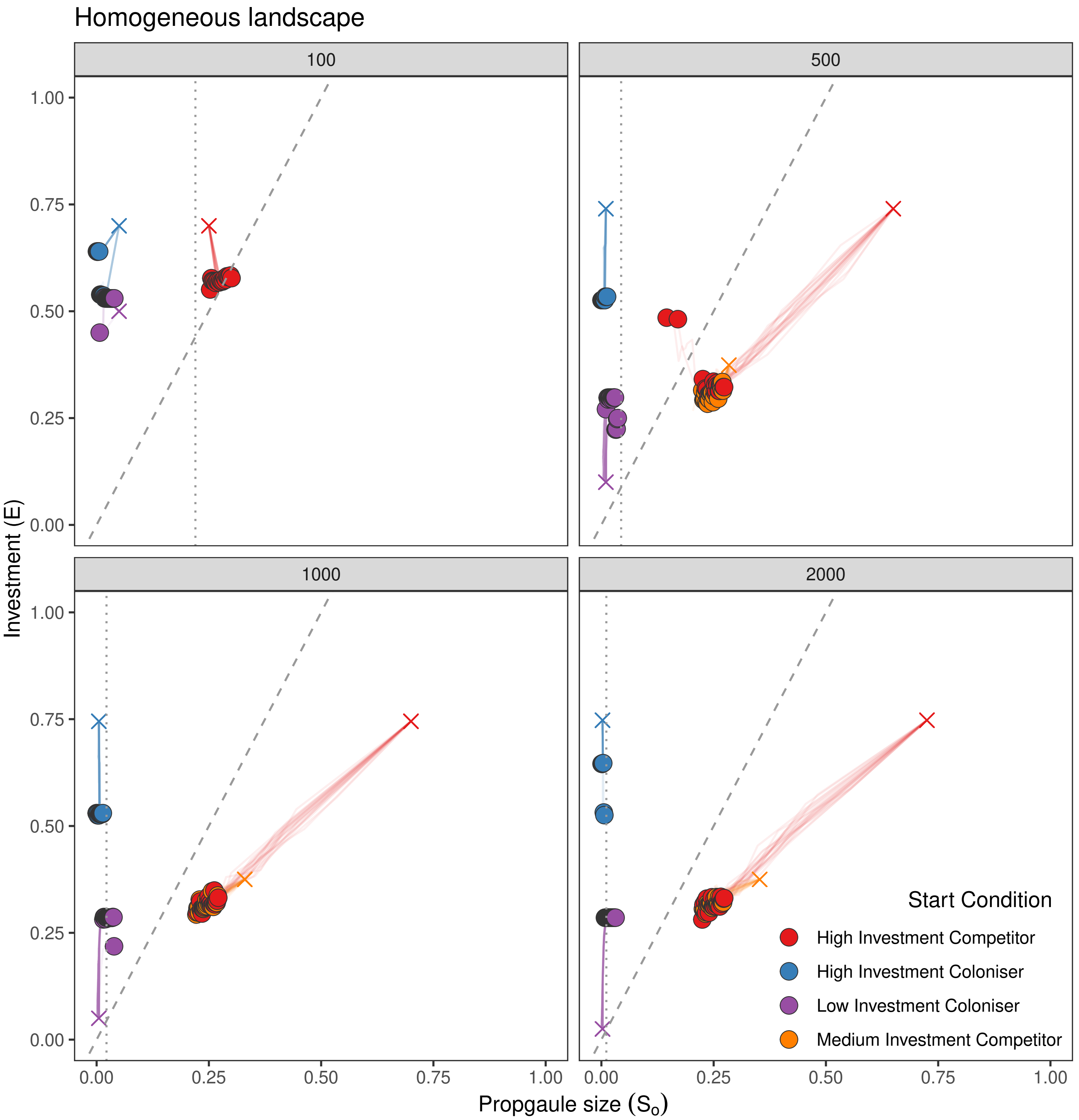


**Figure 1. Evolution of strategies in Homogeneous landscapes (strong size-dispersal trade-off; mutation rate = 0.001)**

Filled circles indicate the final (= mean of all colonies in a simulation) investment (E) and propagule size (So) for different starting conditions for the Homogeneous landscape at different resource levels (K = 100, 500, 1000, 2000). Note that axes (E and So) are written as proportions of the resource level K for that landscape. Starting conditions are indicated with a cross and lines are the temporal dynamics of E and So during simulations (values captured every $10^4$ time-steps). Each coloured straight line corresponds to the mean value of E/So of all colonies in a given simulation. The dashed line separates regions of trait combinations producing more than one propagule (left of line) or a single propagule (right of line). The dotted line indicates the limit between propagules with high dispersal (left of line) and propagules with low dispersal.The position of points has been randomly shifted slightly to aid visualisation.

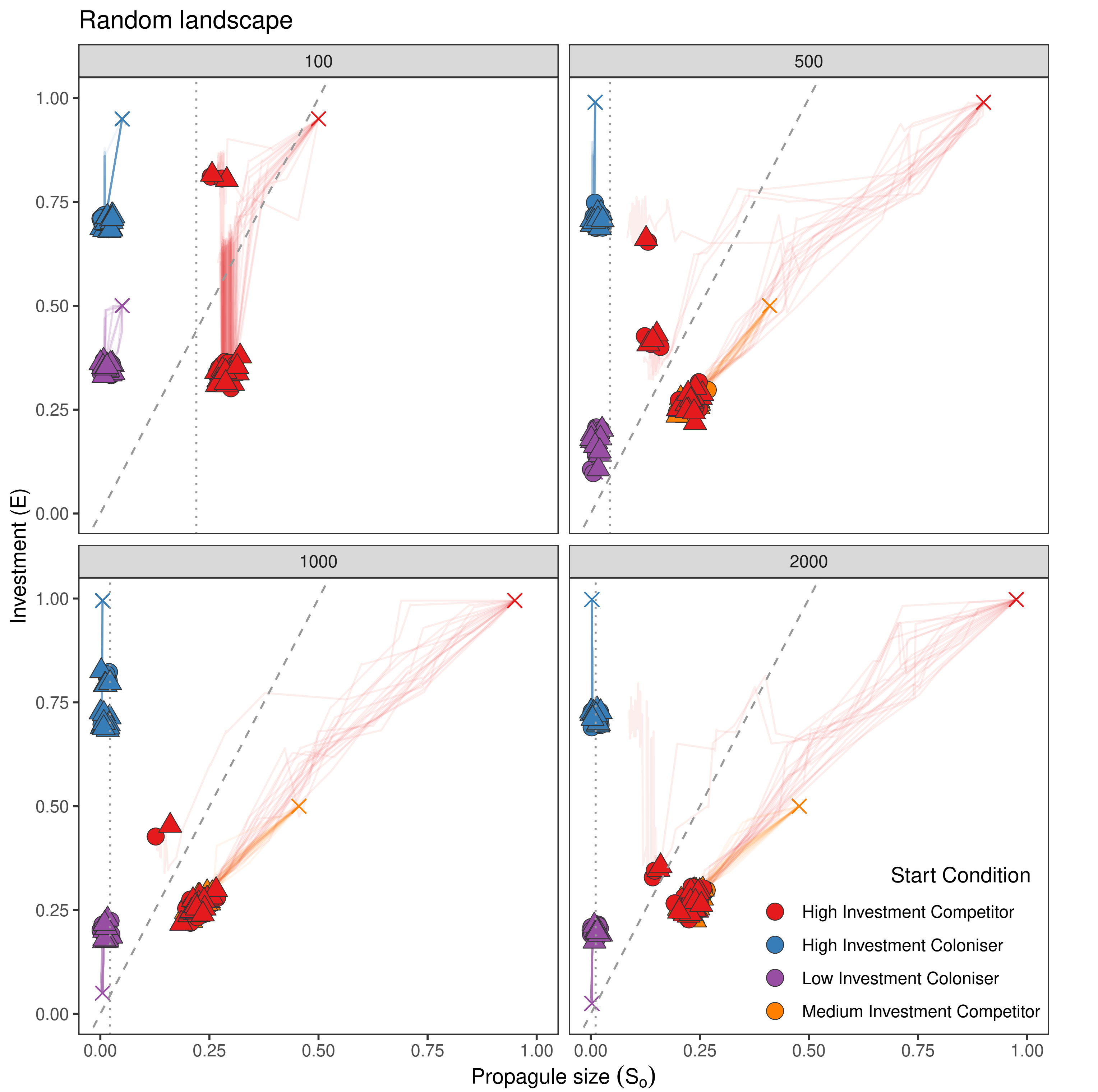


**Figure 2. Evolution of strategies in Random landscapes (strong size-dispersal trade-off; mutation rate = 0.001)**

Filled circles/triangles indicate the final (= mean of all colonies in a simulation) investment (E) and propagule size (So) for different starting conditions for the Random landscape at different resource levels (K = 100, 500, 1000, 2000). Circles indicate colonies in rich patches and triangles indicate colonies in poor patches. Note that axes (E and So) are written as proportions of the resource level K for that landscape. Starting conditions are indicated with a cross and lines are the temporal dynamics of E and So during simulations (values captured every $10^4$ time-steps). Each coloured straight line corresponds to the mean value of E/So of all colonies in a given simulation. The dashed line separates regions of trait combinations producing more than one propagule (left of line) or a single propagule (right of line). The dotted line indicates the limit between propagules with high dispersal (left of line) and propagules with low dispersal.The position of points has been randomly shifted slightly to aid visualisation.

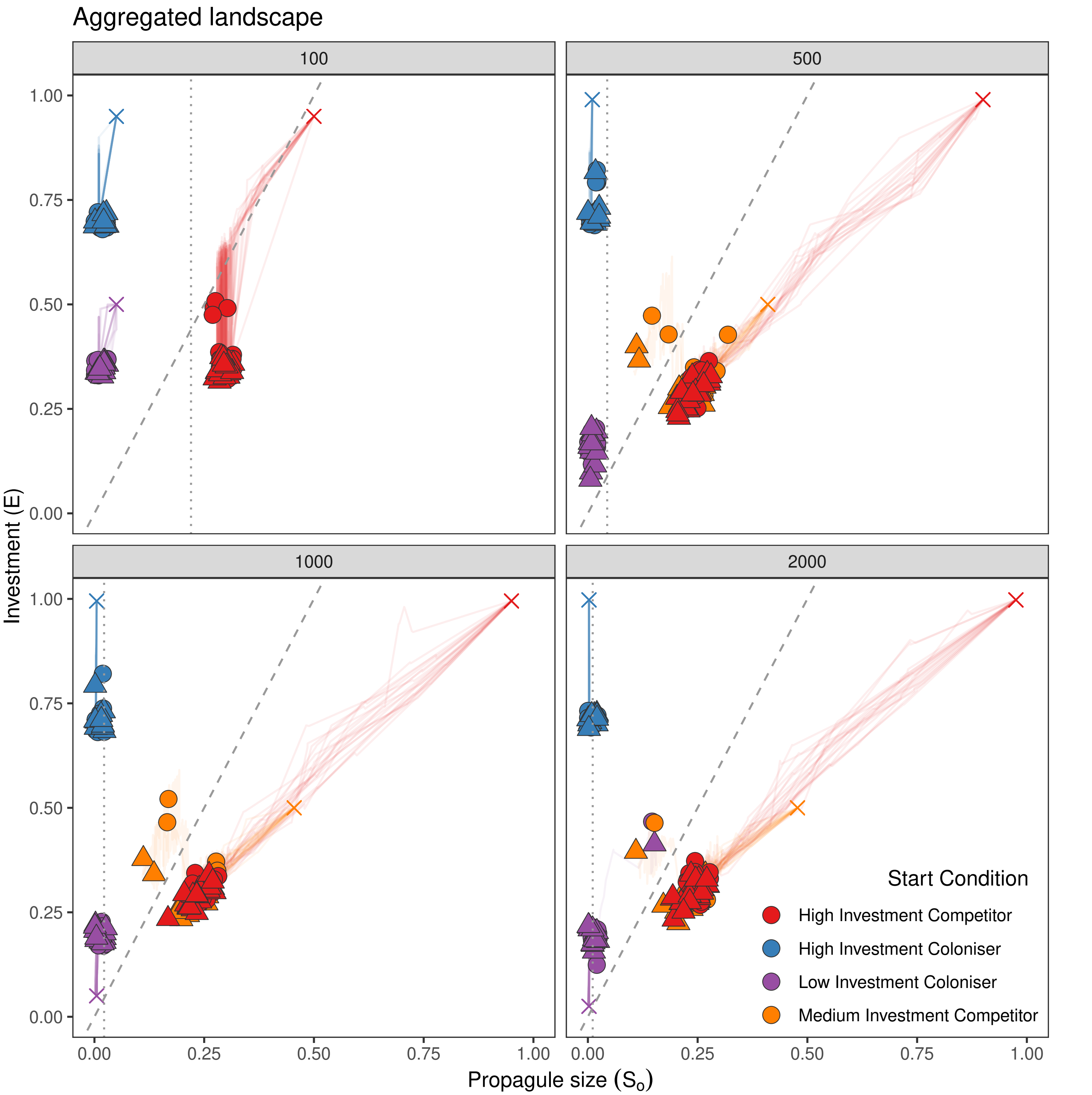


**Figure 3. Evolution of strategies in Aggregated landscapes (strong size-dispersal trade-off; mutation rate = 0.001)**

Filled circles/triangles indicate the final (= mean of all colonies in a simulation) investment (E) and propagule size (So) for different starting conditions for the Aggregated landscape at different resource levels (K = 100, 500, 1000, 2000). Circles indicate colonies in rich patches and triangles indicate colonies in poor patches. Note that axes (E and So) are written as proportions of the resource level K for that landscape. Starting conditions are indicated with a cross and lines are the temporal dynamics of E and So during simulations (values captured every $10^4$ time-steps). Each coloured straight line corresponds to the mean value of E/So of all colonies in a given simulation. The dashed line separates regions of trait combinations producing more than one propagule (left of line) or a single propagule (right of line). The dotted line indicates the limit between propagules with high dispersal (left of line) and propagules with low dispersal.The position of points has been randomly shifted slightly to aid visualisation.
