## Supplementary material for "To disperse or compete? Coevolution of traits leads to a limited number of reproductive strategies": Dynamics for Homogeneous, Random and Aggregated landscapes with weak trade-off

### Supplementary file S4:

Final equilibrium strategies in Homogeneous, Random and Aggregated landscapes for different resource levels (100, 500, 1000 and 2000), and different mutation rates (fast = 0.05 ; slow = 0.001). Simulations were performed with a weak size-dispersal trade-off.

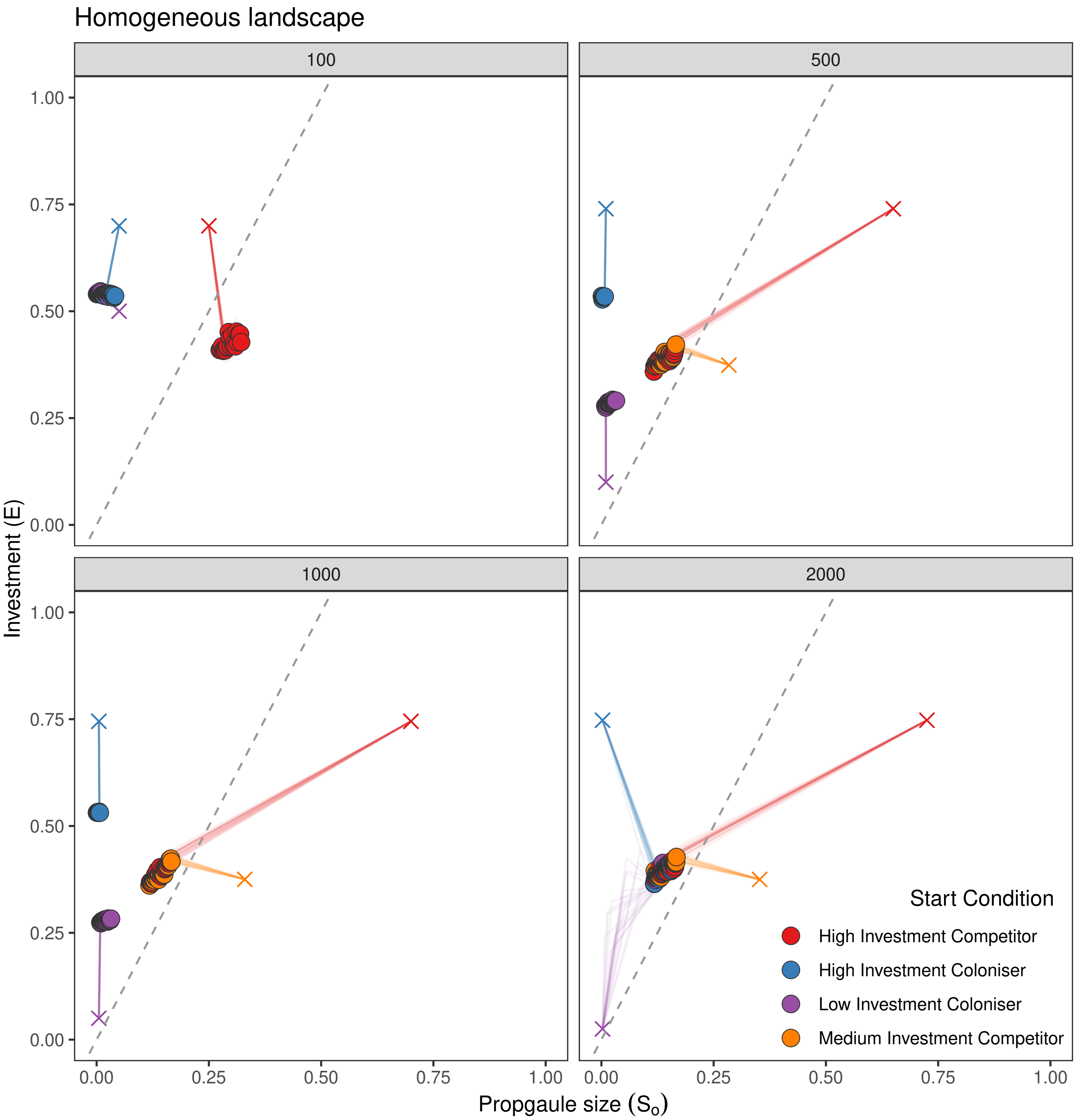


**Figure 1. Evolution of strategies in Homogeneous landscapes (weak size-dispersal trade-off; mutation rate = 0.05)**
Filled circles indicate the final (= mean of all colonies in a simulation) investment (E) and propagule size (So) for different starting conditions for the Homogeneous landscape at different resource levels (K = 100, 500, 1000, 2000). Note that axes (E and So) are written as proportions of the resource level K for that landscape. Starting conditions are indicated with a cross and lines are the temporal dynamics of E and So during simulations (values captured every $10^4$ time-steps). Each coloured straight line corresponds to the mean value of E/So of all colonies in a given simulation. The dashed line separates regions of trait combinations producing more than one propagule (left of line) or a single propagule (right of line). The position of points has been randomly shifted slightly to aid visualisation.

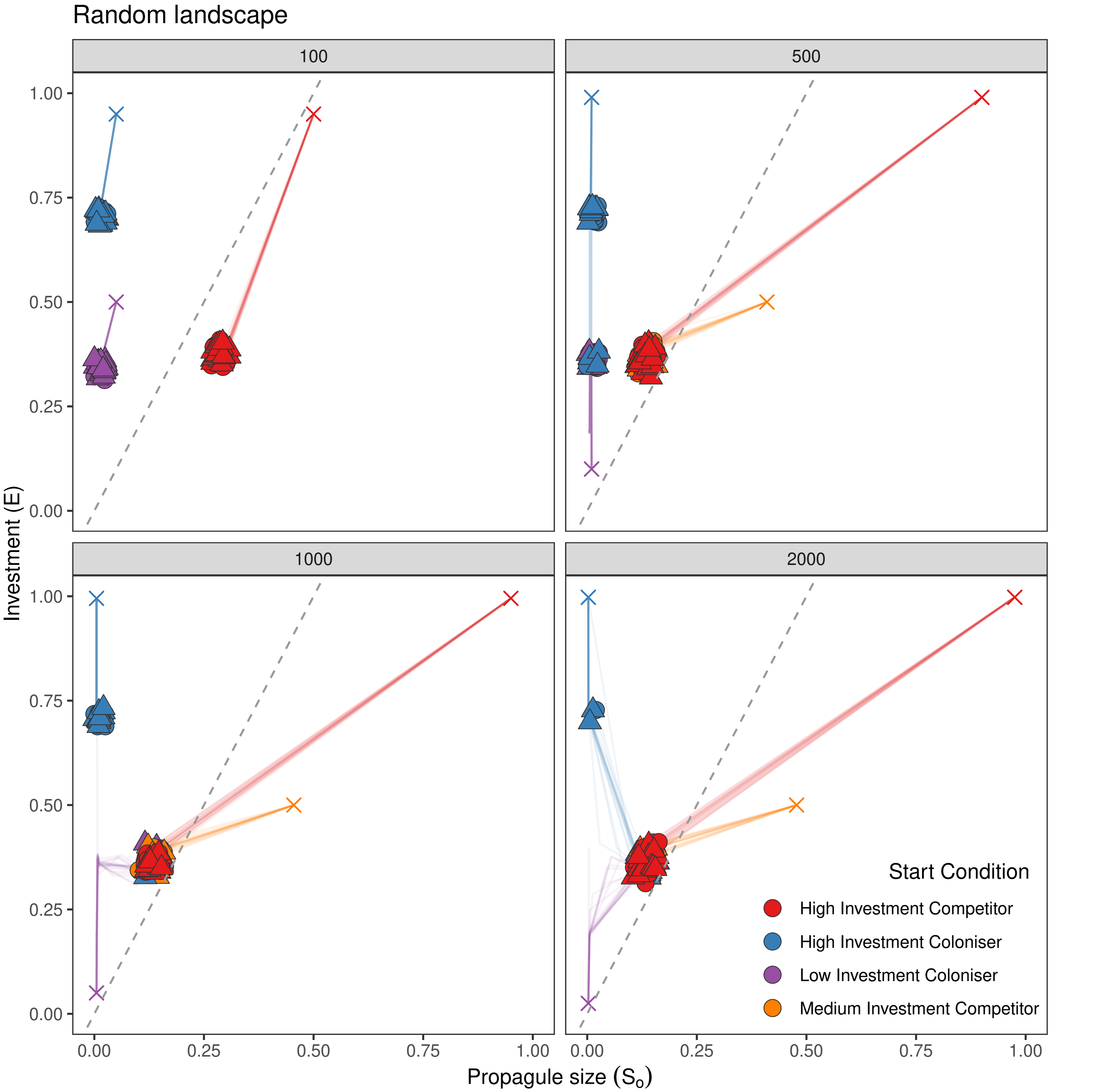


**Figure 2. Evolution of strategies in Random landscapes (weak size-dispersal trade-off; mutation rate = 0.05)**
Filled circles/triangles indicate the final (= mean of all colonies in a simulation) investment (E) and propagule size (So) for different starting conditions for the Random landscape at different resource levels (K = 100, 500, 1000, 2000). Circles indicate colonies in rich patches and triangles indicate colonies in poor patches. Note that axes (E and So) are written as proportions of the resource level K for that landscape. Starting conditions are indicated with a cross and lines are the temporal dynamics of E and So during simulations (values captured every $10^4$ time-steps). Each coloured straight line corresponds to the mean value of E/So of all colonies in a given simulation. The dashed line separates regions of trait combinations producing more than one propagule (left of line) or a single propagule (right of line). The position of points has been randomly shifted slightly to aid visualisation.

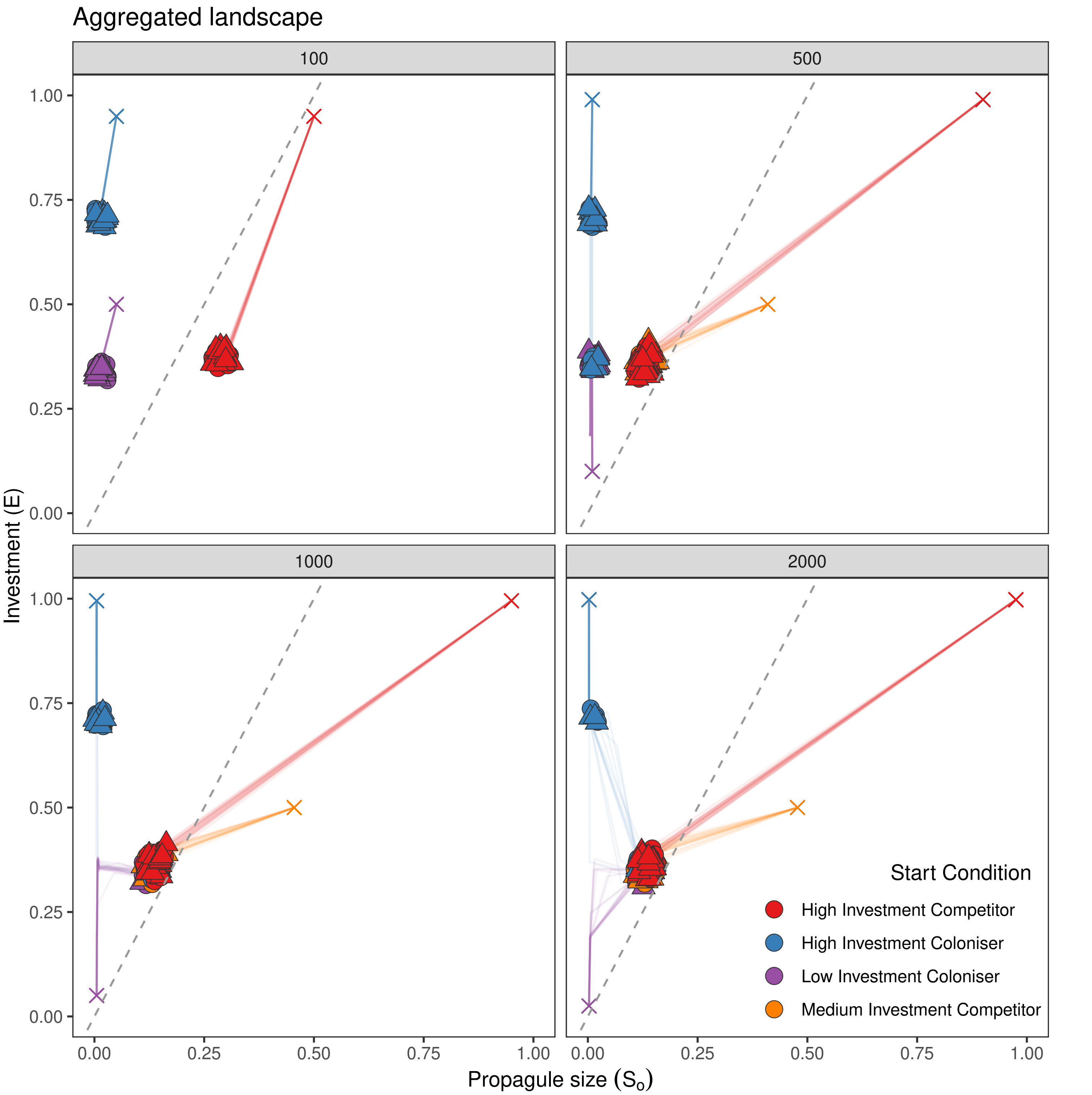


**Figure 3. Evolution of strategies in Aggregated landscapes (weak size-dispersal trade-off; mutation rate = 0.05)**
Filled circles/triangles indicate the final (= mean of all colonies in a simulation) investment (E) and propagule size (So) for different starting conditions for the Aggregated landscape at different resource levels (K = 100, 500, 1000, 2000). Circles indicate colonies in rich patches and triangles indicate colonies in poor patches. Note that axes (E and So) are written as proportions of the resource level K for that landscape. Starting conditions are indicated with a cross and lines are the temporal dynamics of E and So during simulations (values captured every $10^4$ time-steps). Each coloured straight line corresponds to the mean value of E/So of all colonies in a given simulation. The dashed line separates regions of trait combinations producing more than one propagule (left of line) or a single propagule (right of line). The position of points has been randomly shifted slightly to aid visualisation.

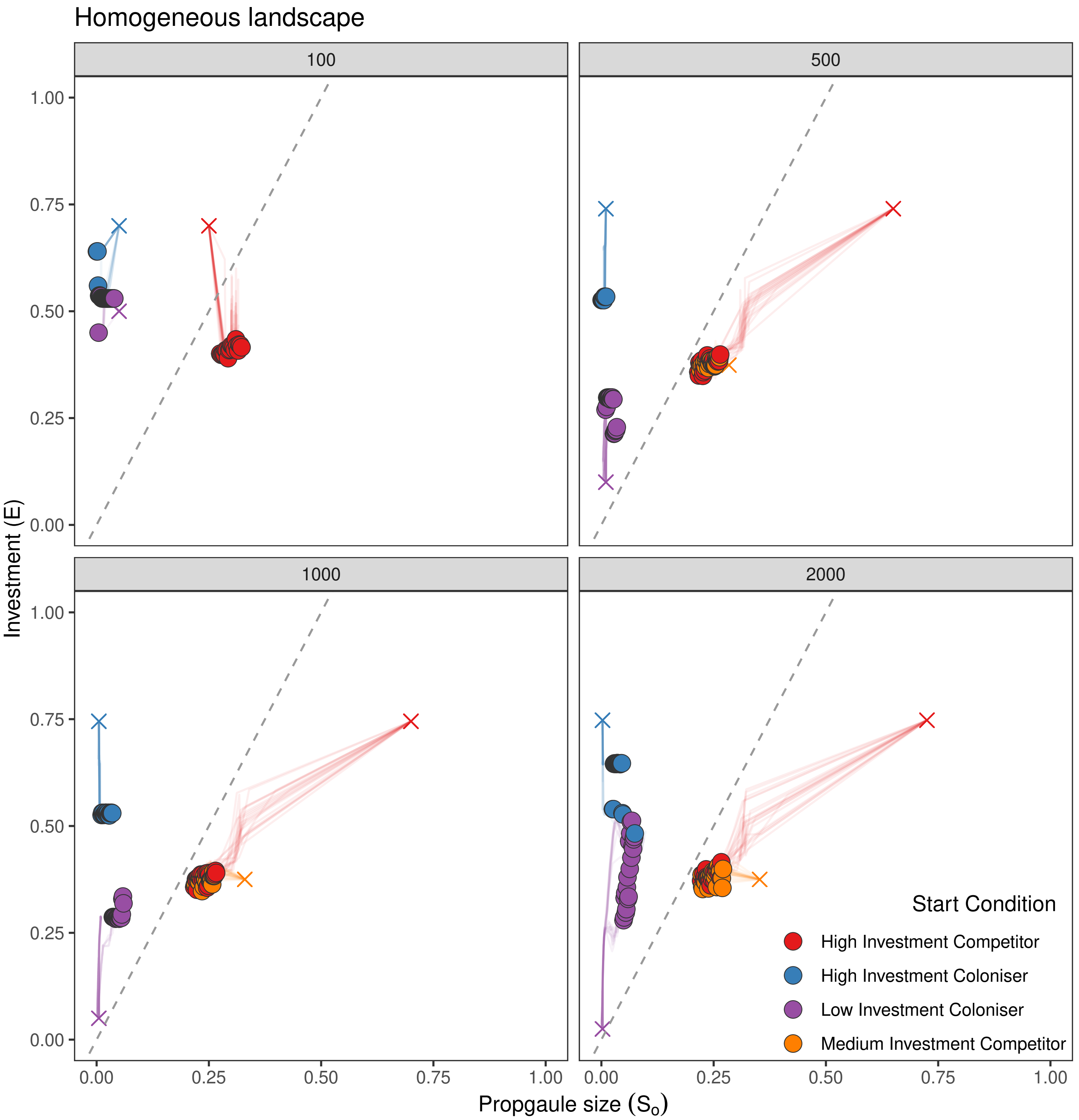


**Figure 4. Evolution of strategies in Homogeneous landscapes (weak size-dispersal trade-off; mutation rate = 0.001)**
Filled circles indicate the final (= mean of all colonies in a simulation) investment (E) and propagule size (So) for different starting conditions for the Homogeneous landscape at different resource levels (K = 100, 500, 1000, 2000). Note that axes (E and So) are written as proportions of the resource level K for that landscape. Starting conditions are indicated with a cross and lines are the temporal dynamics of E and So during simulations (values captured every $10^4$ time-steps). Each coloured straight line corresponds to the mean value of E/So of all colonies in a given simulation. The dashed line separates regions of trait combinations producing more than one propagule (left of line) or a single propagule (right of line). The position of points has been randomly shifted slightly to aid visualisation.

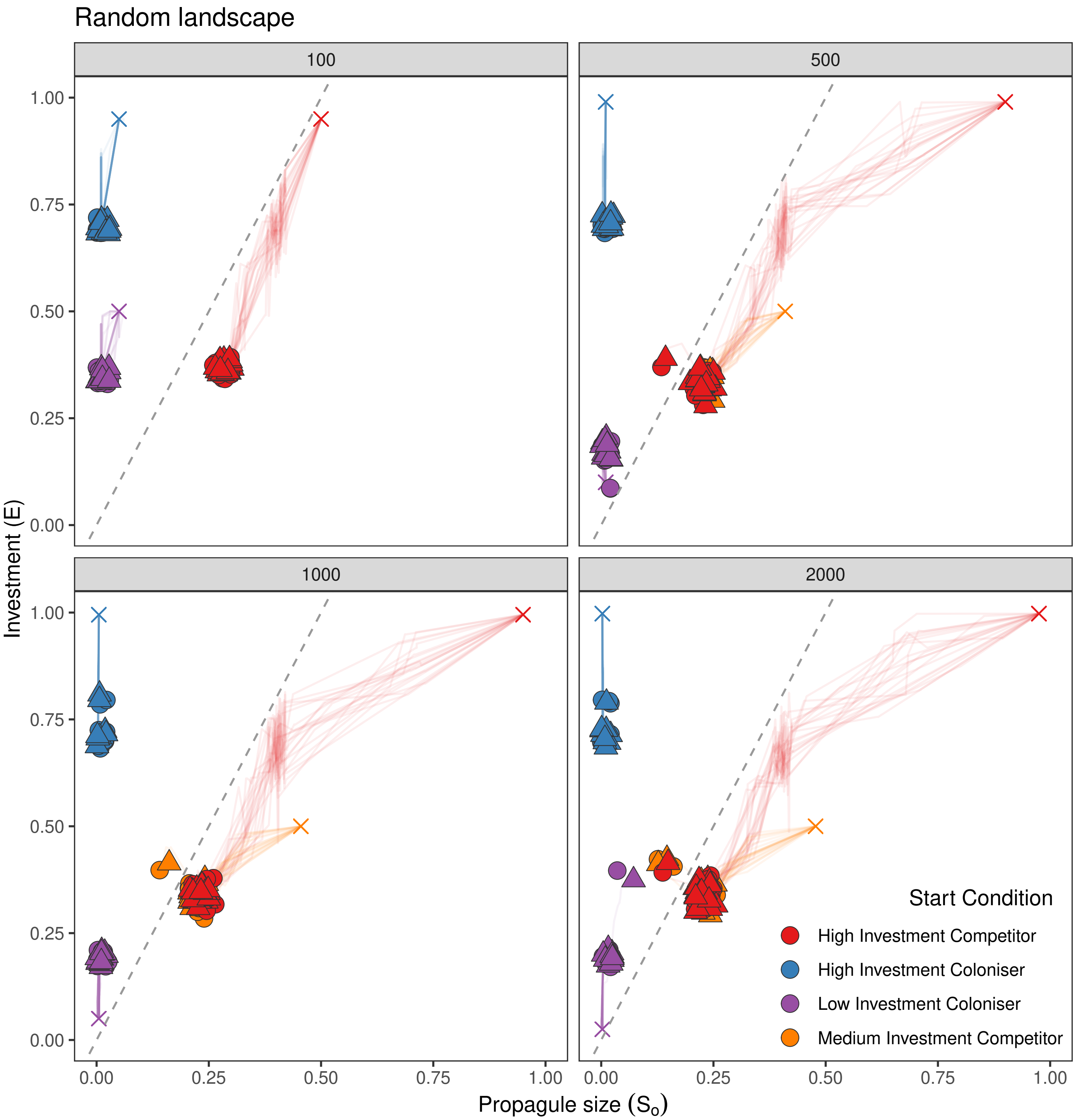


**Figure 5. Evolution of strategies in Random landscapes (weak size-dispersal trade-off; mutation rate = 0.001)**

Filled circles/triangles indicate the final (= mean of all colonies in a simulation) investment (E) and propagule size (So) for different starting conditions for the Random landscape at different resource levels (K = 100, 500, 1000, 2000). Circles indicate colonies in rich patches and triangles indicate colonies in poor patches. Note that axes (E and So) are written as proportions of the resource level K for that landscape. Starting conditions are indicated with a cross and lines are the temporal dynamics of E and So during simulations (values captured every $10^4$ time-steps). Each coloured straight line corresponds to the mean value of E/So of all colonies in a given simulation. The dashed line separates regions of trait combinations producing more than one propagule (left of line) or a single propagule (right of line). The position of points has been randomly shifted slightly to aid visualisation.

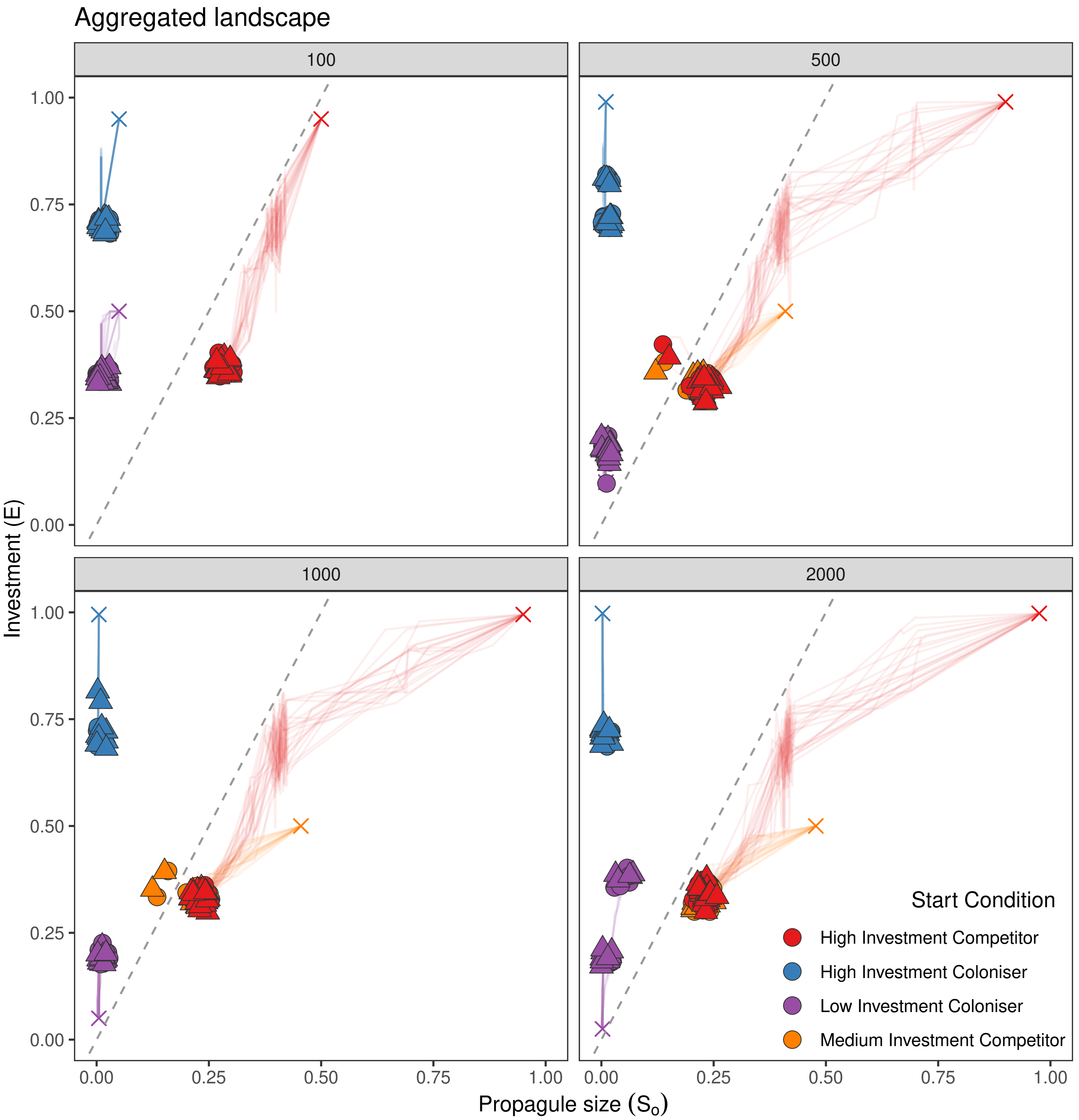


**Figure 6. Evolution of strategies in Aggregated landscapes (weak size-dispersal trade-off; mutation rate = 0.001)**
Filled circles/triangles indicate the final (= mean of all colonies in a simulation) investment (E) and propagule size (So) for different starting conditions for the Aggregated landscape at different resource levels (K = 100, 500, 1000, 2000). Circles indicate colonies in rich patches and triangles indicate colonies in poor patches. Note that axes (E and So) are written as proportions of the resource level K for that landscape. Starting conditions are indicated with a cross and lines are the temporal dynamics of E and So during simulations (values captured every $10^4$ time-steps). Each coloured straight line corresponds to the mean value of E/So of all colonies in a given simulation. The dashed line separates regions of trait combinations producing more than one propagule (left of line) or a single propagule (right of line). The position of points has been randomly shifted slightly to aid visualisation.
