## Supplementary material for "To disperse or compete? Coevolution of traits leads to a limited number of reproductive strategies": Invasion analysis

### Supplementary file S5:

Outcome of the invasion analysis in Homogeneous, Random and Aggregated landscapes for different resource levels (100, 500, 1000 and 2000) and strong/weak size-dispersal trade-off. Axes titles refer to equilibrium strategies.

#### Strong size-dispersal trade-off:

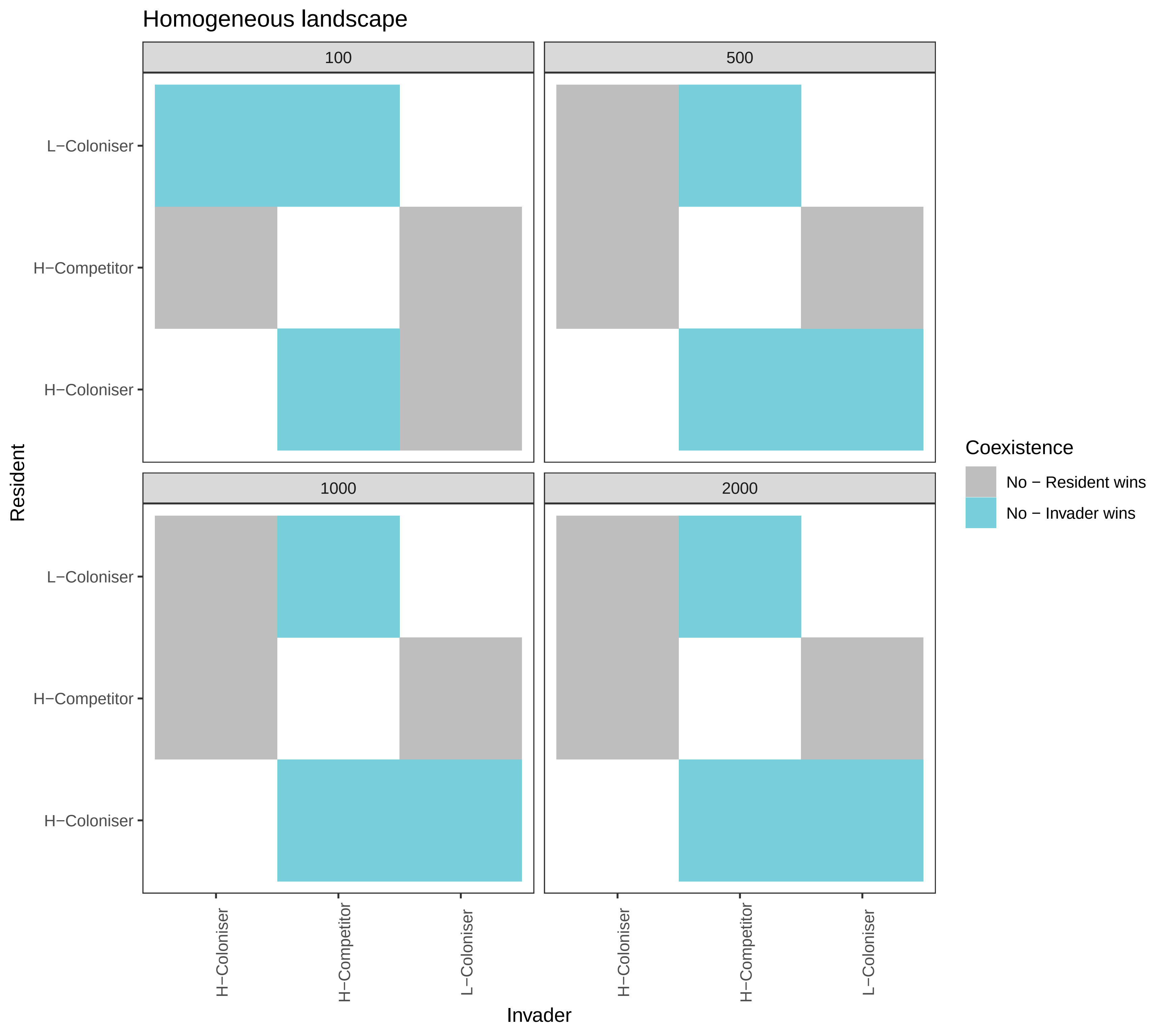

Homogeneous landscape
100
500
1000
2000
L−Coloniser
H−Competitor
H−Coloniser
Resident
Invader
Coexistence
No − Resident wins
No − Invader wins

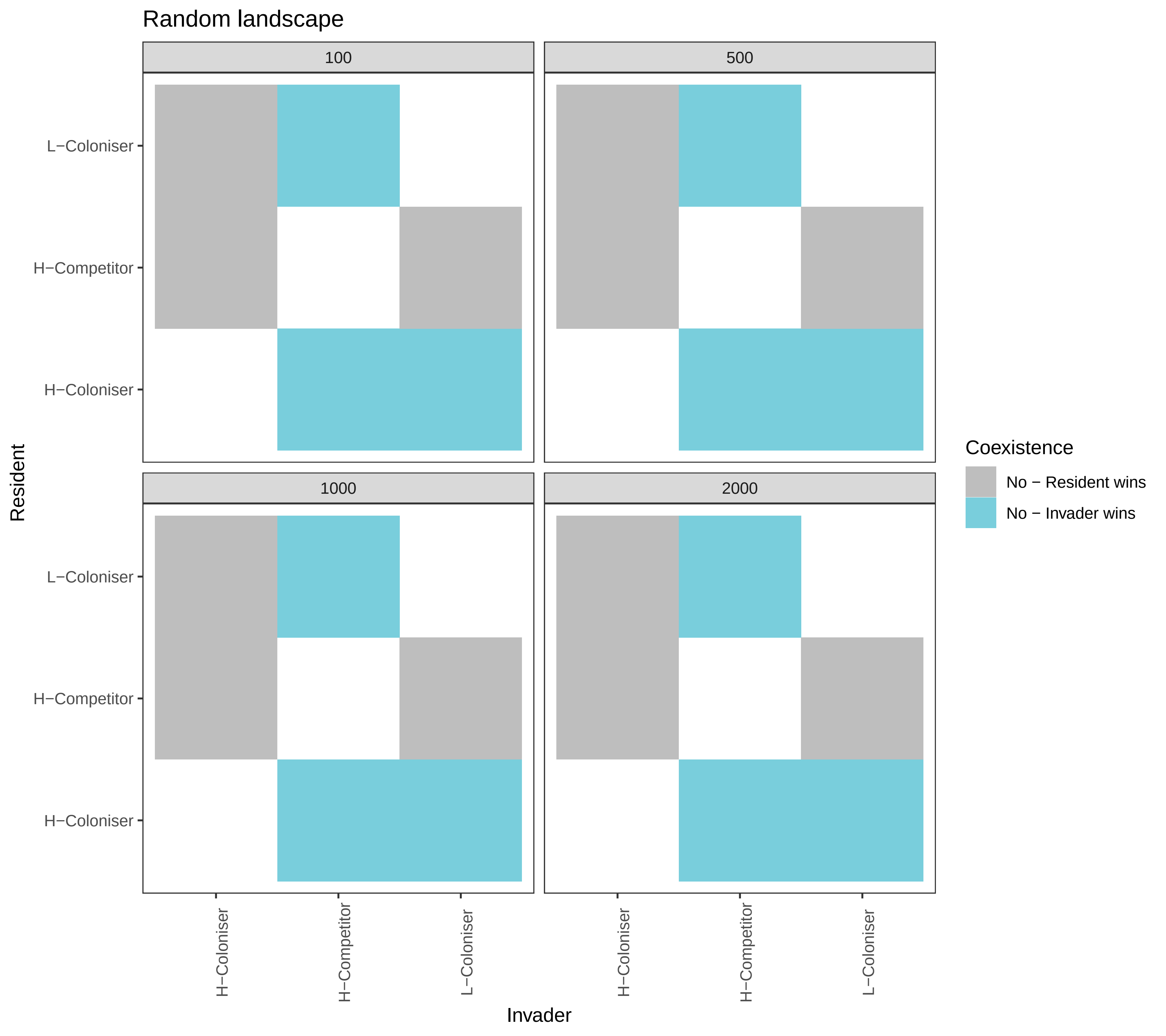

Random landscape
100
500
1000
2000
L−Coloniser
H−Competitor
H−Coloniser
Resident
Invader
Coexistence
No − Resident wins
No − Invader wins

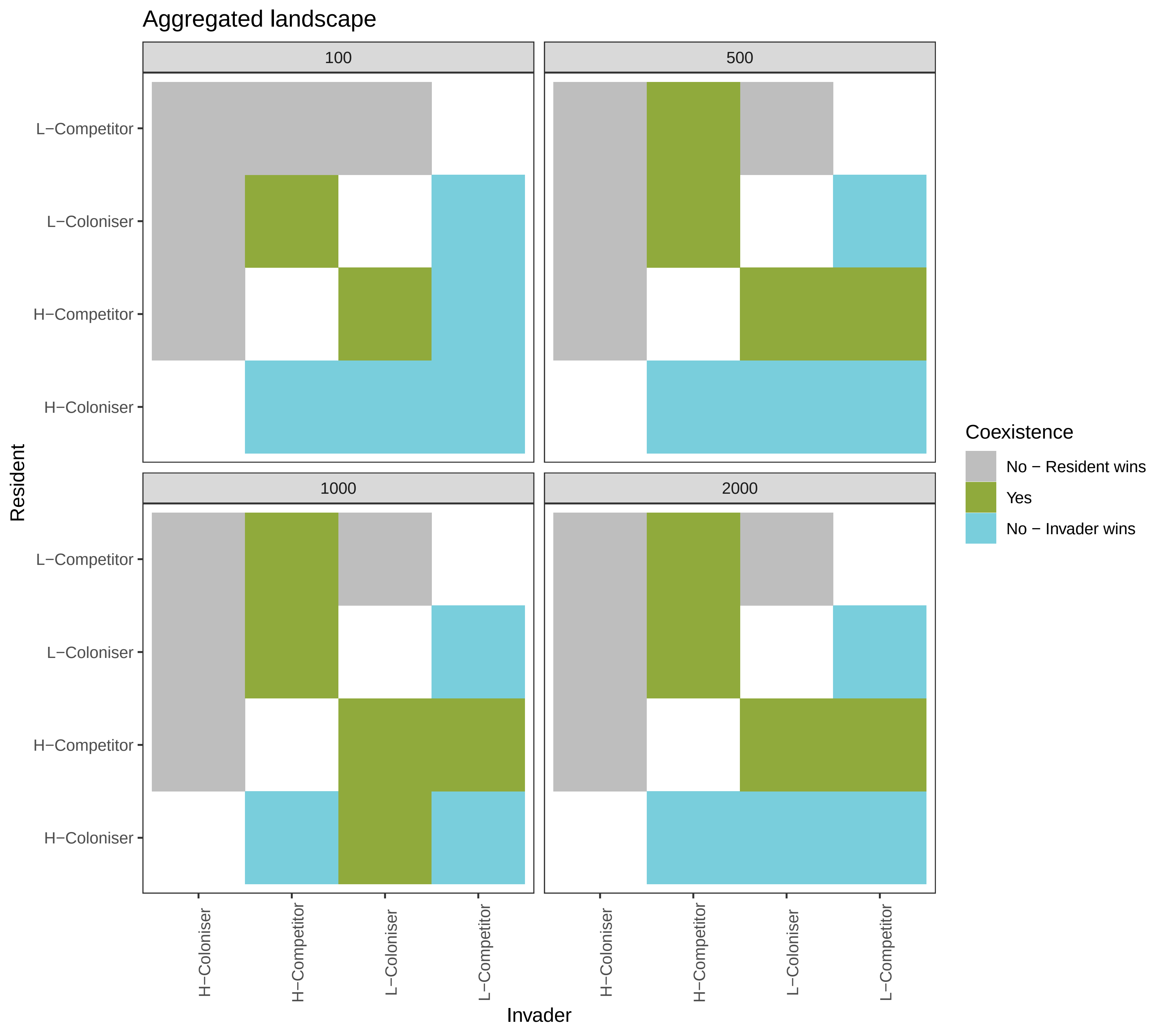
Aggregated landscape
100
500
1000
2000
L−Competitor
L−Coloniser
H−Competitor
H−Coloniser
Resident
H−Coloniser
H−Competitor
L−Coloniser
L−Competitor
Invader
Coexistence
No − Resident wins
Yes
No − Invader wins

#### Weak size-dispersal trade-off:

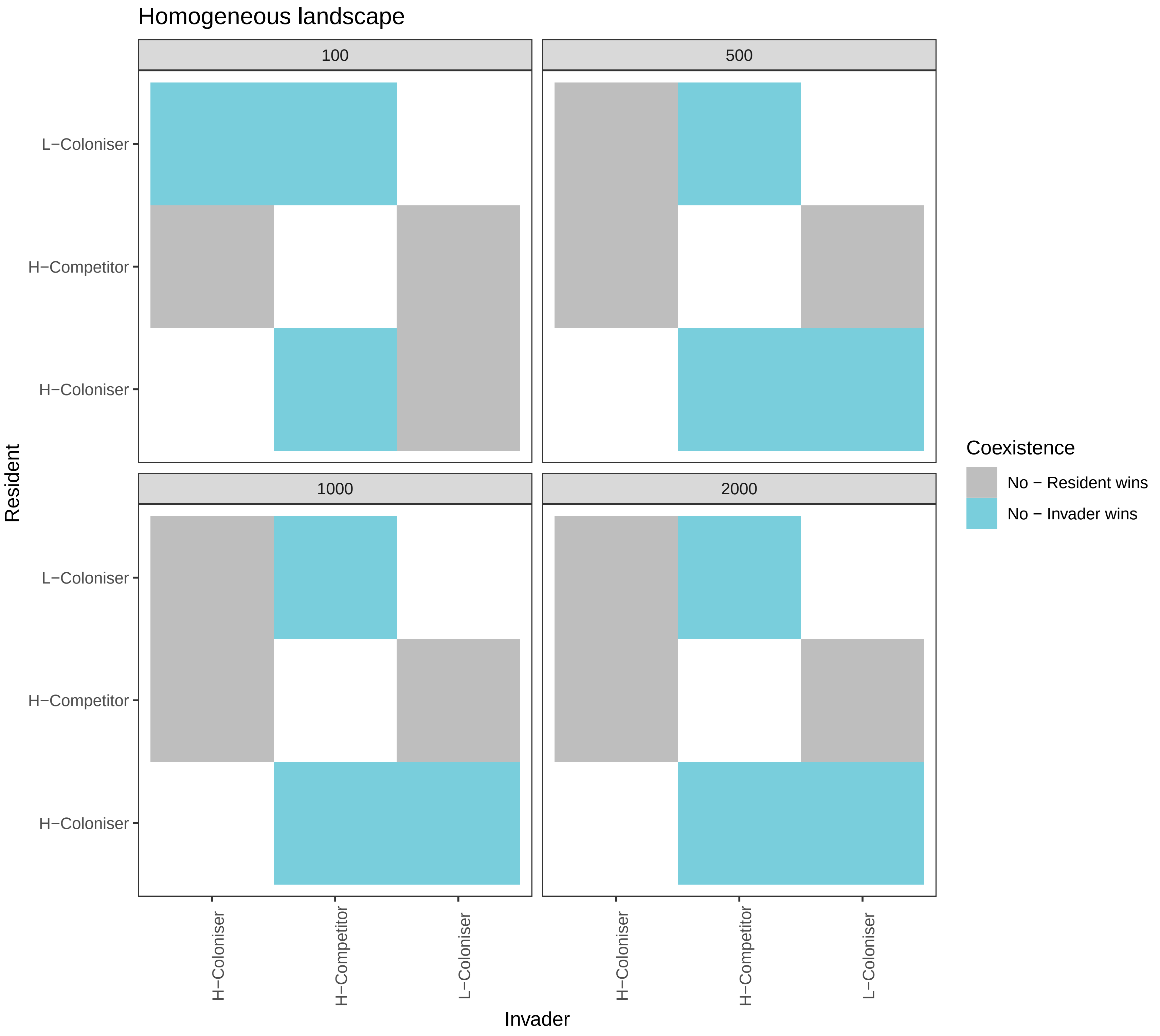
Homogeneous landscape
100
500
1000
2000
L−Coloniser
H−Competitor
H−Coloniser
Resident
Invader
Coexistence
No − Resident wins
No − Invader wins

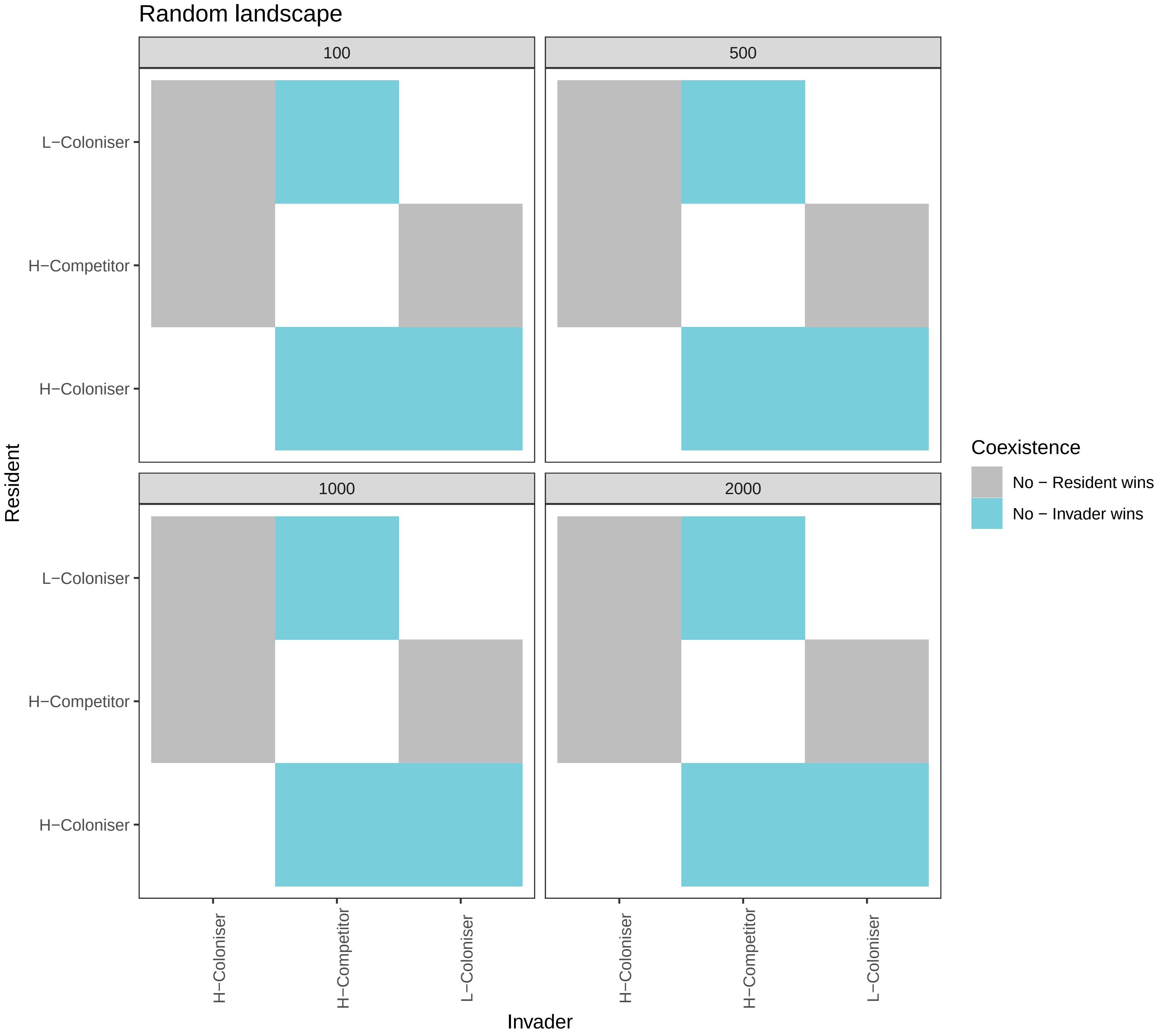
Random landscape
100
500
1000
2000
L−Coloniser
H−Competitor
H−Coloniser
Resident
H−Coloniser
H−Competitor
L−Coloniser
Invader
Coexistence
No − Resident wins
No − Invader wins

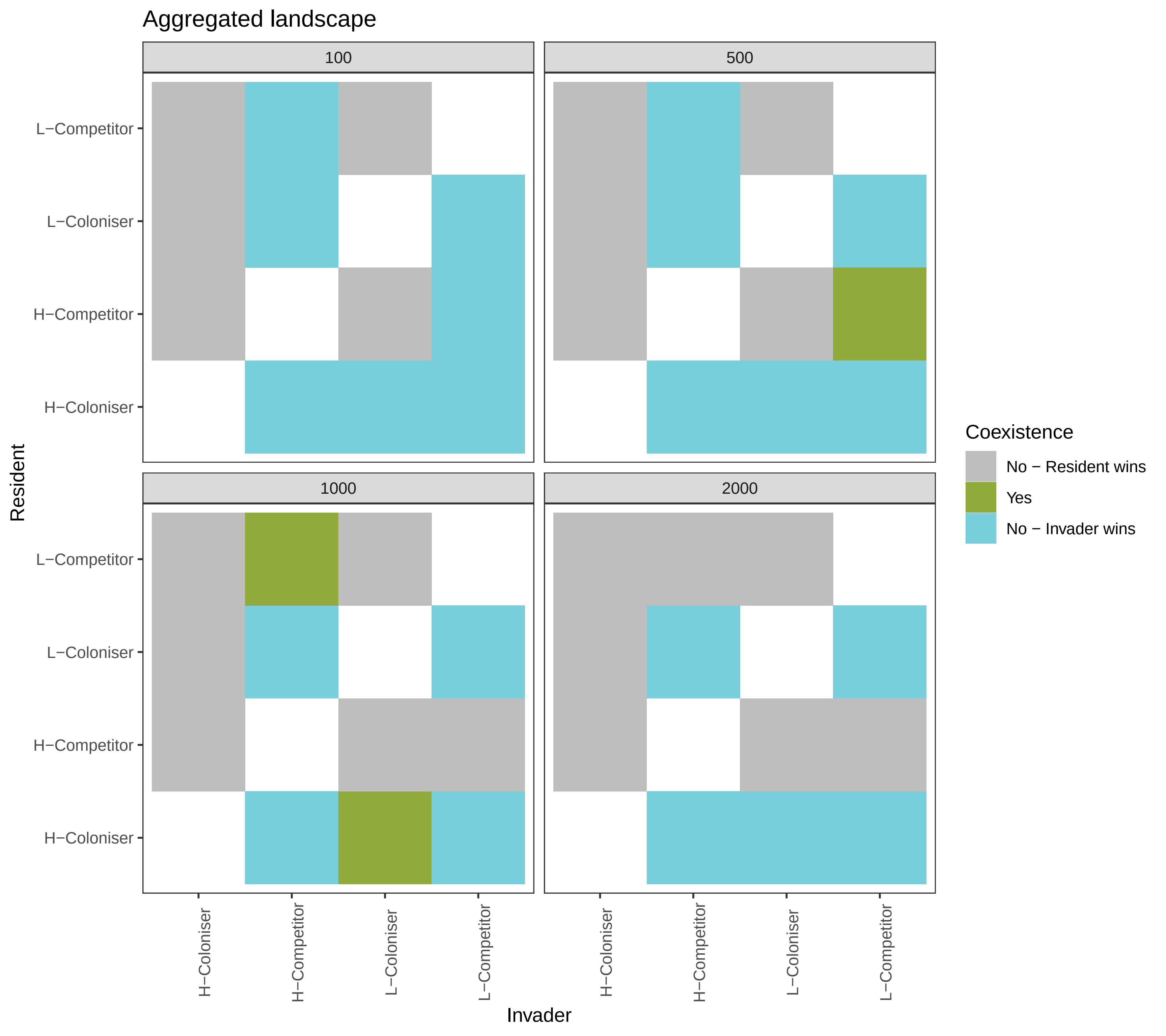
Aggregated landscape
100
500
1000
2000
L−Competitor
L−Coloniser
H−Competitor
H−Coloniser
Resident
H−Coloniser
H−Competitor
L−Coloniser
L−Competitor
Invader
Coexistence
No − Resident wins
Yes
No − Invader wins
